# Unexpected Specificity within Dynamic Transcriptional Protein-Protein Complexes

**DOI:** 10.1101/2020.05.29.124271

**Authors:** Matthew J. Henley, Brian M. Linhares, Brittany S. Morgan, Tomasz Cierpicki, Carol A. Fierke, Anna K. Mapp

**Affiliations:** Life Sciences Institute, University of Michigan, Ann Arbor, MI 48109; Program in Chemical Biology, University of Michigan, Ann Arbor, MI 48109; Department of Biophysics, University of Michigan, Ann Arbor, MI 48109; Department of Pathology, University of Michigan, Ann Arbor, MI 48109; Department of Chemistry, Texas A&M University, College Station, TX 77843; Department of Chemistry, University of Michigan, Ann Arbor, MI 48109

**Author notes:** **Significance** Transcriptional activators represent a molecular recognition enigma. Their function in transcription initiation requires selective engagement of coactivators yet the prevailing molecular recognition models propose this occurs via nonspecific intermolecular contacts. Here, mechanistic analysis of several related activator•coactivator complexes resolves this conundrum. In contrast to the expectations from nonspecific recognition models, even small sequence changes in the activators cause activator•coactivator complexes to undergo significant conformational redistribution, driven by specific intermolecular interactions and conformational changes in the coactivator itself. These unappreciated *specific* recognition mechanisms rationalize the high sequence variability of functional activators, opening new questions about the relationship between recognition and function. <u>Dedication:</u> We dedicate this work to Professor Laura L. Kiessling, on the occasion of her 60_th_ birthday. Her pioneering work on molecular recognition principles within transiently associated macromolecular complexes has been an inspiration.

## Abstract

A key functional event in eukaryotic gene activation is the formation of dynamic protein-protein interaction networks between transcriptional activators and transcriptional coactivators. Seemingly incongruent with the tight regulation of transcription, many biochemical and biophysical studies suggest that activators use nonspecific hydrophobic and/or electrostatic interactions to bind to coactivators, with few if any specific contacts. Here a mechanistic dissection of a set of representative dynamic activator•coactivator complexes, comprised of the ETV/PEA3 family of activators and the coactivator Med25, reveals a different molecular recognition model. The data demonstrate that small sequence variations within an activator family significantly redistribute the conformational ensemble of the complex while not affecting overall affinity, and distal residues within the activator—not often considered as contributing to binding—play a key role in mediating conformational redistribution. The ETV/PEA3•Med25 ensembles are directed by specific contacts between the disordered activator and the Med25 interface, which is facilitated by structural shifts of the coactivator binding surface. Taken together, these data highlight the critical role coactivator plasticity plays in recognition of disordered activators, and indicates that molecular recognition models of disordered proteins must consider the ability of the binding partners to mediate specificity.

## Introduction

Protein-protein interactions (PPIs) formed between transcriptional activators and coactivators play a critical role in gene expression; these PPIs underpin co-localization of transcriptional machinery components and stimulate transcription initiation.(1–8) A prevailing view of activator-coactivator PPIs is that they are largely non-specific and, further, that the selectivity necessary for appropriate gene expression comes from other sources such as activator-DNA interactions and/or co-localization.(3, 9–14) Indeed, there is considerable data suggesting activator•coactivator complexes form via almost entirely nonspecific intermolecular interactions, from early experiments demonstrating that a wide range of natural and non-natural amphipathic molecules interact with coactivators to more recent structural studies indicating no fixed activator•coactivator binding mode (Fig. 1).(1, 9–11, 15–18)

**Figure 1.**
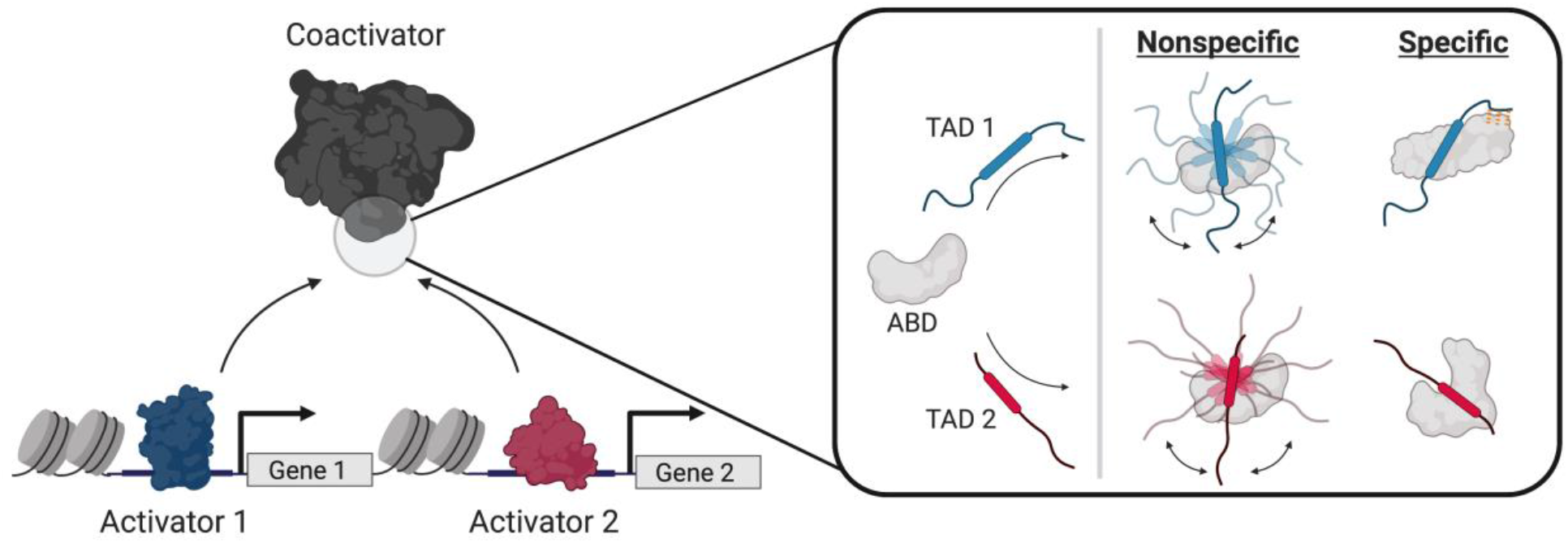
Recognition models of activator function. Left: Transcriptional activators regulate gene activity via protein-protein interactions with coactivators. Right: Comparison of nonspecific (3, 11) and specific (this work) models of activator•coactivator recognition. Nonspecific models propose that the transcriptional activation domains (TADs) of unique activators bind the activator binding domains (ABDs) of coactivators via nonspecific intermolecular interactions, forming complexes without fixed orientation or structure via “sequence-independent” recognition.(11) Rounded boxes in the TADs represent the binding motif.

Nonspecific recognition models, while attractive in their simplicity, are inconsistent with the critical functional role that individual activator•coactivator PPIs play in gene expression. There are several examples of transcriptional activators that depend on interactions with specific activator binding domains (ABDs) of coactivators for function,(19–23) e.g., the SREBP family of activators require the KIX ABD of the coactivator ARC105 to regulate fatty acid homeostasis(21) even though other coactivators such as CBP and p300 have structurally similar KIX motifs. Further, the biophysical studies that investigate how ABDs recognize diverse activators most often utilize qualitative equilibrium approaches(9–11) that are blind to critical mechanistic information(24–26) due to equilibrium averaging. It is therefore an open question whether there are other molecular recognition mechanisms at play; this would account for the diversity of functional activator sequences as well as the observed selectivity of activators *in vivo*. Because activator•coactivator complexes often represent promising therapeutic targets, developing a more detailed understanding of the molecular recognition mechanisms of these crucial PPIs is also essential for the development of small molecule modulators.(4, 27–29)

Here, we take a critical look at activator•coactivator molecular recognition by mechanistically dissecting a representative set of dynamic complexes formed between the ABD of Mediator subunit Med25 and the amphipathic transcriptional activation domains (TADs) of the ETV/PEA3 family of Ets transcriptional activators (ETV1, ETV4, and ETV5).(23, 30–32) Previous biophysical studies indicated that the interaction of Med25 with family member ETV5 appears to be a prototypical nonspecific TAD•ABD complex: it occurs over a shallow surface, is driven by electrostatic and hydrophobic interactions, and forms a dynamic complex that is recalcitrant to structure determination.(30, 32) We utilize a mechanistic and structural approach that combines quantitative data regarding activator•coactivator conformational states obtained via transient kinetic analysis with structural information obtained through mutagenesis and NMR spectroscopy. Our data reveal that the conformational ensembles of ETV/PEA3•Med25 PPIs are strikingly sensitive to slight changes in TAD sequence, despite being dynamic complexes with several well-populated conformational sub-states at equilibrium. Furthermore, the mechanisms by which conformational sensitivity involve the ability of ordered and disordered regions of the TAD to participate in finite sets of *specific* interactions with the Med25 interface, as well as conformational changes in Med25 that remodel the TAD binding site. Together, these results reveal an unappreciated degree of specificity in the formation of activator•coactivator complexes that is in direct contrast to the prevailing nonspecific recognition models of these essential PPIs (Fig. 1).(3, 11)

## Results

### ETV/PEA3•Med25 PPIs as a Model System for Dynamic TAD•ABD Interactions

ETV/PEA3•Med25 interactions represent an ideal system to study dynamic TAD•ABD interactions for several reasons. First, previous studies indicated that interaction of ETV/PEA3 family member ETV5 with the Med25 ABD is typical of a dynamic TAD•ABD complex(30, 32); the binding surface is shallow, both acidic and hydrophobic amino acids of the TAD determine affinity and activity, and multiple bound conformational states have been detected by both NMR and kinetic analyses. Second, our recent studies of the ETV5•Med25 complex showed that the bound conformational ensemble is directly accessible by transient kinetic analysis.(32) The models of TAD•ABD molecular recognition can thus be dissected in this system without the loss of critical conformational information to equilibrium averaging. Third, the ETV/PEA3 family of transcription factors serves as an excellent natural system to test the relationship between TAD sequence and recognition; they contain almost identical arrangements of acidic and hydrophobic residues across the TAD sequence, especially within the helical binding region that undergoes coupled folding and binding with Med25,(30) but the identity of specific residues varies slightly (Fig. 2A). This system can therefore be used to test whether TAD•Med25 interactions are truly nonspecific and insensitive to variations in the TAD sequence(11), or if these interactions are affected by TAD sequence changes and thus have a degree of specificity to formation. Finally, the Med25 ABD is ligandable by small molecules(32), and therefore conclusions from mechanistic studies can be directly applied to guide and assess optimization of small molecule modulators of TAD•Med25 complex formation.

**Figure 2.**
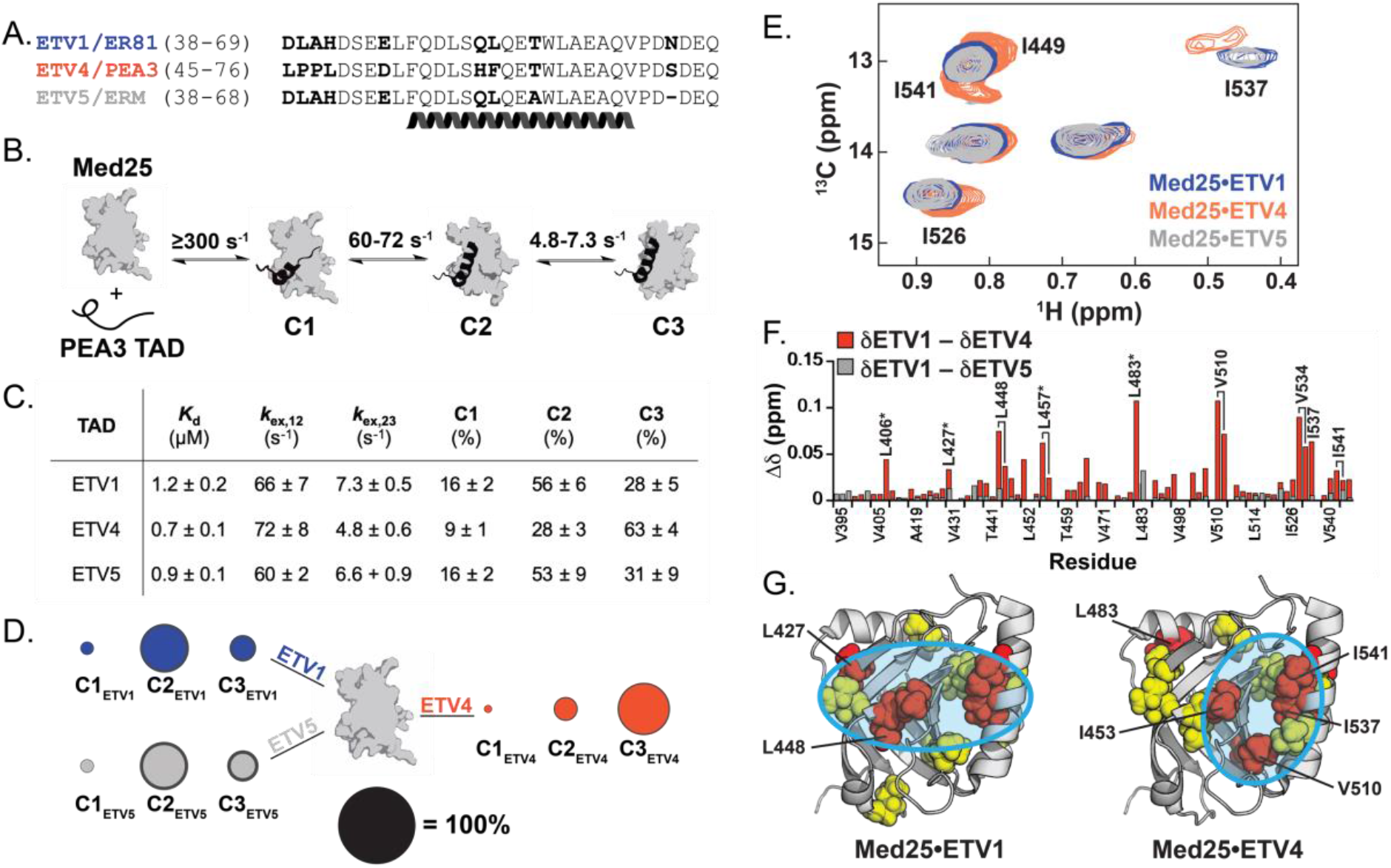
ETV/PEA3 activators differentially engage with Med25. A) Alignment of ETV/PEA3 family activation domains. The helix denotes the residues that undergo coupled folding and binding with Med25, as determined by NMR chemical shift analysis.*(30)* B) Mechanism of binding of ETV/PEA3 activators to Med25, determined here for ETV1 and ETV4, and previously for ETV5.*(32)* The range of exchange rates between analogous steps for ETV/PEA3 TADs are shown. C) Table of relevant binding parameters for each ETV/PEA3 TAD, including the equilibrium affinity, exchange rates between C1 and C2 (*k*_ex,12_) and between C2 and C3 (*k*_ex,23_), and equilibrium populations of each state. All values represent the average of 3-4 biological replicates, and the error is the standard deviation. D) Equilibrium populations of the three ETV/PEA3•Med25 conformations scaled relative to the diameter of the black circle. Standard deviations of the values are shown as the dark gray outer circle. E) Overlay of the Ile Cδ region of the _1_H,_13_C-HSQC of Med25 in complex with 1.1 equivalents unlabeled ETV1 (blue), ETV4 (orange), or ETV5 (grey). Peaks that have chemical shift differences between complexes are labeled. Note: I449 and I541 form a single overlapped peak for the ETV1 and ETV5 complexes. Full spectra are in the supporting information. F) Chemical shift differences of Med25 methyl resonances between ETV1 and ETV4 (orange) and ETV1 and ETV5 (grey). G) Chemical shift perturbations induced by binding of ETV/PEA3 activators plotted on the structure of Med25 (PDB ID 2XNF)*(34)*. Yellow = 0.040–0.080 ppm, red > 0.080 ppm. Cyan circles highlight general distinctions in perturbation patterns. Several residues with chemical shift differences >0.030 ppm between the ETV1•Med25 (or ETV5•Med25) and ETV4•Med25 complexes are labeled.

### Small Sequence Differences Between ETV/PEA3 Family Members Lead to Conformationally Distinct PPIs with Med25

We first examined whether the slight sequence variations across the ETV/PEA3 family TADs affect the conformational ensembles of the individual ETV/PEA3•Med25 complexes. Stopped-flow fluorescence transient kinetic experiments were performed to directly evaluate conformational dynamics and equilibria, using TADs synthesized with the solvatochromic fluorophore 4-DMN conjugated to the *N*-terminus.(33) We previously demonstrated with this approach that the ETV5•Med25 complex forms in a minimal three step linear mechanism(32): after an initial rapid association event that mostly occurs in the instrument dead-time (~2-4 ms), the complex undergoes two sequential conformational changes (Fig 2B). At equilibrium, all three bound conformations of the ETV5•Med25 complex are well populated due to relatively small conformational equilibrium constants.(32)

Application of the same experimental conditions to ETV1 and ETV4 indicated that the kinetic binding mechanism is conserved; for all ETV/PEA3 TADs a rapid binding step followed by two conformational change steps was observed. Each of these individual steps occurred with similar exchange rate constants (*k*_ex_; the sum of the forward and reverse rate constants) for each ETV/PEA3•Med25 complex (Fig. 2B,C); this suggests that the steps represent analogous conformational transitions in each complex. In addition, the equilibrium binding affinity between ETV/PEA3 TADs varied less than two-fold (0.7-1.2 μM, Fig. 2C), consistent with the expectation from nonspecific models that minor substitutions in the TAD will not affect the overall stability of the activator•coactivator complex.

Despite a conserved binding mechanism and similar overall affinities, calculation of equilibrium conformational populations from the kinetic data revealed clear differences between the engagement modes of ETV/PEA3 family members (Fig. 2C,D; for raw data and detailed kinetic analysis, see SI Discussion of Kinetic Analysis). While the populations of analogous conformational states of the ETV1•Med25 and ETV5•Med25 complexes were essentially identical, the ETV4•Med25 complex populated the three analogous conformations in a unique manner (Fig 2C,D). Critically, this shift in conformational equilibria does not correlate with predicted structural propensity differences between the ETV/PEA3 TADs (SI Fig. S1), which suggests that variable residues between ETV1/ETV5 and ETV4 alter the TAD•Med25 conformational ensemble via intermolecular interactions made in the bound state.

We next examined differences in the engagement modes of ETV/PEA3 TADs via NMR spectroscopy. Unique TADs with nonspecific engagement modes are expected to produce essentially identical binding signatures in NMR spectra of the partner ABD,(11) therefore comparison of chemical shift perturbation (CSP) patterns of labeled Med25 bound to individual ETV/PEA3 TADs provides an orthogonal source of insight into the specificity of the interaction. Side-chain methyl _1_H,_13_C-HSQC experiments were used as a primary method to enable direct detection of effects on both surface and buried residues of Med25.

Comparative analysis of Med25 _1_H,_13_C-HSQC spectra bound to different ETV/PEA3 family members was consistent with the expected engagement differences between the ETV/PEA3 TADs: spectra of ETV1- and ETV4-bound Med25 exhibited several large differences in CSP patterns, whereas the spectra of Med25 bound to ETV1 and ETV5 were essentially indistinguishable (Fig. 2E,F; full spectral overlay shown in SI Fig. S19). Inspection of the CSP data plotted on the structure of Med25 indicated that all ETV/PEA3 family members bind to a previously identified(30–32) core binding site formed between the central β-barrel and the *C*-terminal α3 helix (Fig. 2G), but ETV1/ETV5 and ETV4 produce unique perturbation patterns in the binding surface (Fig. 2G, cyan circles). In addition, several resonances representing buried and/or allosteric residues displayed significant CSP differences between the ETV1- and ETV4-bound complexes, suggesting that the conformation of the Med25 ABD may also be different between these complexes (Fig. 2F, starred). The NMR data therefore supports the conclusion from kinetics experiments that ETV4 has a unique Med25 engagement mode as compared to ETV1 and ETV5; together, these biophysical and structural experiments suggest a model where one or more of the variable residues in the ETV/PEA3 TADs make distinct *specific* intermolecular interactions with the Med25 surface.

### Ordered and Disordered Regions of the ETV/PEA3 TADs Dictate Conformational Differences between ETV/PEA3•Med25 PPIs

We next identified the TAD residues that bias ETV/PEA3•Med25 PPIs towards distinct conformational sub-states using a mutagenesis approach. This effort focused on residues that are conserved between ETV1 and ETV5, but not ETV4. Two regions of interest were evident (Fig. 3A, boxed): 1) a two amino acid ‘variable motif’ in the helical binding region consisting of a polar residue followed by a hydrophobic residue (QL in ETV1/ETV5 and HF in ETV4), and 2) the four amino acid *N*-terminus of the TAD (DLAH in ETV1/ETV5 and LPPL in ETV4), a region predicted to be entirely disordered for all ETV/PEA3 TADs (SI Fig. S1 and Ref. 30). A small library of mutant TADs varying residues in the two regions of interest of the ETV4 sequence—either the Leu or Phe residue in the variable motif and the DLAH or LPPL *N*-terminus—was synthesized and assessed in stopped-flow kinetic assays (Fig. 3B; values for all variants in SI Tables S2 and S3). The polar residue (Gln/His) in the variable motif was also tested but had no effect on conformational populations. Importantly, the equilibrium binding affinity varied only slightly across the mutants (Fig. 3D).

**Figure 3.**
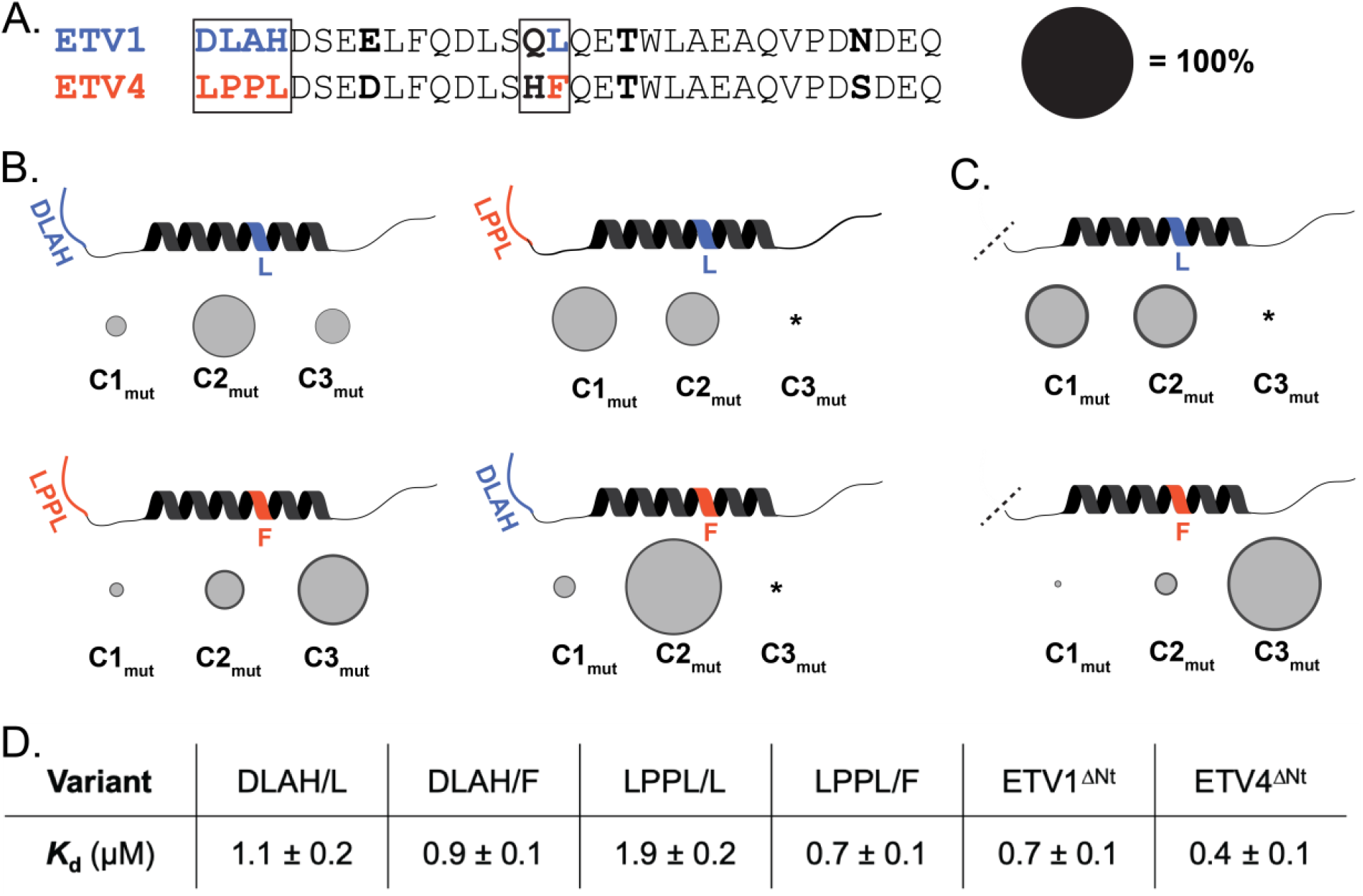
Variable residues in the disordered *N*-terminus and the helical binding region mediate differences in ETV/PEA3•Med25 conformational behavior. A) Alignment of ETV1 and ETV4 activators with regions that were selected for mutational analysis boxed. Regions/residues that affected the conformational ensemble are color coded to ETV1 (blue) or ETV4 (orange). Effects of the Gln/His residues in the variable motif were also tested but did not affect conformational populations and are thus omitted in B. and C. for clarity. Populations of conformational states in B. and C. are scaled relative to the diameter of the black circle. B) Results from kinetics experiments of mutant TADs, for native (left) and non-native (right) combinations of variable *N*-termini and helical binding regions. Variants were made based on the ETV4 sequence. The data shown is the average across all the variants tested from each group, with the error (dark grey outer circle) representing the standard deviation. C) Results from kinetics experiments with ETV1_ΔNt_ (left) and ETV4_ΔNt_ (right). D) Average equilibrium *K*_d_ values of variants tested. *Conformer was undetectable in kinetics experiments (see SI *Discussion of Kinetic Data Analysis* for further details).

Consistent with the hypothesis that one or both of the variable regions dictate the conformational differences between the ETV/PEA3•Med25 complexes, ETV/PEA3 variants with ‘native’ combinations of the *N*-terminus and the hydrophobic residue of the variable motif exhibited similar conformational ensembles to the natural TADs (Fig. 3B, left). That is, combinations with DLAH at the *N*-terminus and Leu in the variable motif (DLAH/L) populated three observable conformational states in a similar manner to ETV1 and ETV5 (Fig. 3B, top left), and, analogously, combinations with LPPL at the *N*-terminus and Phe in the variable motif (LPPL/F) populated three sub-states comparably to ETV4 (Fig. 3B, bottom left).

Conversely, when nonnative combinations of the *N*-terminus and variable motif were tested, unique conformational behavior was observed (Fig. 3B, right). Only two bound conformations of DLAH/F variants were detected in kinetics experiments, and displayed exchange kinetics similar to the C1–C2 transitions of the native complexes (Fig. 3B, top right). Calculation of conformational populations indicated that the second conformation had a higher overall population (82 ± 3%) than the C2 conformations of the ETV1 (56 ± 6%), ETV5 (53 ± 9%), or ETV4 complexes (28 ± 3%). Similarly, two bound conformations with C1–C2-like exchange rates were detected with LPPL/L variants (Fig 3B, bottom right), but the initial bound sub-state was preferentially populated. We note that the C3 conformation of these complexes may be undetectable due to a low population (≤10%) or an increase in the exchange kinetics between C2–C3; however, neither of these possibilities are inconsistent with the conclusion that these mutant complexes are conformationally distinct (see SI *Discussion of Kinetic Data Analysis* for further details). Together, these results indicate that the conformational differences between ETV/PEA3•Med25 complexes are dictated by the identity of *both* the hydrophobic residue in the variable motif and the disordered *N*-terminus. The latter result is particularly striking, as disordered regions of the TAD are often removed/ignored in biophysical and structural studies because they typically do not contribute to overall affinity.(9–11, 25, 35–40)

To obtain further evidence for the unexpected role of the disordered *N*-terminus on the conformational behavior of ETV/PEA3•Med25 PPIs, we also tested the effects of removing the four variable *N*-terminal residues (ΔNt) of ETV1 and ETV4 in kinetics experiments. The resulting variants ETV1_ΔNt_ and ETV4_ΔNt_ displayed differential changes in conformational behavior from the parent TADs (Fig. 3C), in addition to a slight (~1.7-fold) gain in affinity for both variants (Fig. 3D). Removal of the ETV1 *N*-terminus resulted in significant conformational redistribution; kinetic analysis indicated that the ETV1_ΔNt_•Med25 complex exchanged between two equally populated conformational sub-states on a similar timescale to the C1–C2 transition of the parent ETV1•Med25 complex. On the other hand, the ETV4_ΔNt_•Med25 complex populated three conformational sub-states in an analogous manner to the parent ETV4•Med25 complex. These results therefore support a direct role of the *N*-terminal residues of the ETV1/ETV5 TADs, but not the ETV4 TAD, in biasing the conformational behavior of the native TAD•Med25 complexes.

### Variable Regions of the ETV/PEA3 TADs Differentially Engage with the Med25 Surface

We next directly examined the structural basis by which the two variable regions in the ETV/PEA3 TADs modulate the bound ETV/PEA3•Med25 conformations using NMR spectroscopy. A conservative mutagenesis strategy was pursued, where minimally perturbing mutations were individually introduced into unlabeled TADs, and then CSP analysis of _1_H,_13_C-HSQC spectra of Med25 bound to the native or mutated TAD was performed to identify Med25 methyl groups affected by the mutation. Analysis of CSP differences in the mutant TAD•Med25 HSQC spectrum can therefore detect Med25 residues in direct proximity to the mutated site in the complex and also has the potential to reveal allosteric connections if the effects of the mutation are propagated from the interaction site.(41) Furthermore, this strategy avoids the significant experimental challenge associated with NMR analysis of Med25-bound ETV/PEA3 TADs, where a significant fraction of peaks are too broad to detect due to conformational exchange.(30) Here, conservative mutations were introduced into the two key variable regions of the ETV1 and ETV4 TADs to detect differences in engagement modes that could explain the effects of these regions on ETV/PEA3•Med25 conformational states.

Mutations were first made within the variable motif of the helical binding region by swapping the variable polar residue between ETV1 and ETV4 to form ETV1_Q52H_ and ETV4_H59Q_ (note: residue numbers for ETV4 are shifted by +7 compared to ETV1 and ETV5), based on the observation that this change did not affect the populations of conformational states in kinetics experiments (SI Table S3). Indeed, the _1_H,_13_C-HSQC spectra of the mutant ETV/PEA3•Med25 complexes were almost identical to those of the native complexes except for single shifts in unique methyl peaks of Ile541 (Fig. 4A): the ETV4_H59Q_ variant produced a large shift (0.032 ppm) in the Med25 Ile541δ peak, whereas the analogous ETV1_Q52H_ variant produced a smaller perturbation (0.015 ppm) in the Ile541γ peak. These highly localized shifts are consistent with the mutations causing proximity-based perturbations near the mutation site in the complex. In addition, the fact that the individual mutations perturb unique methyl groups originating from the same residue suggest that the native ETV1 QL and ETV4 HF motifs are engaged in unique interactions in a similar interface.

**Figure 4.**
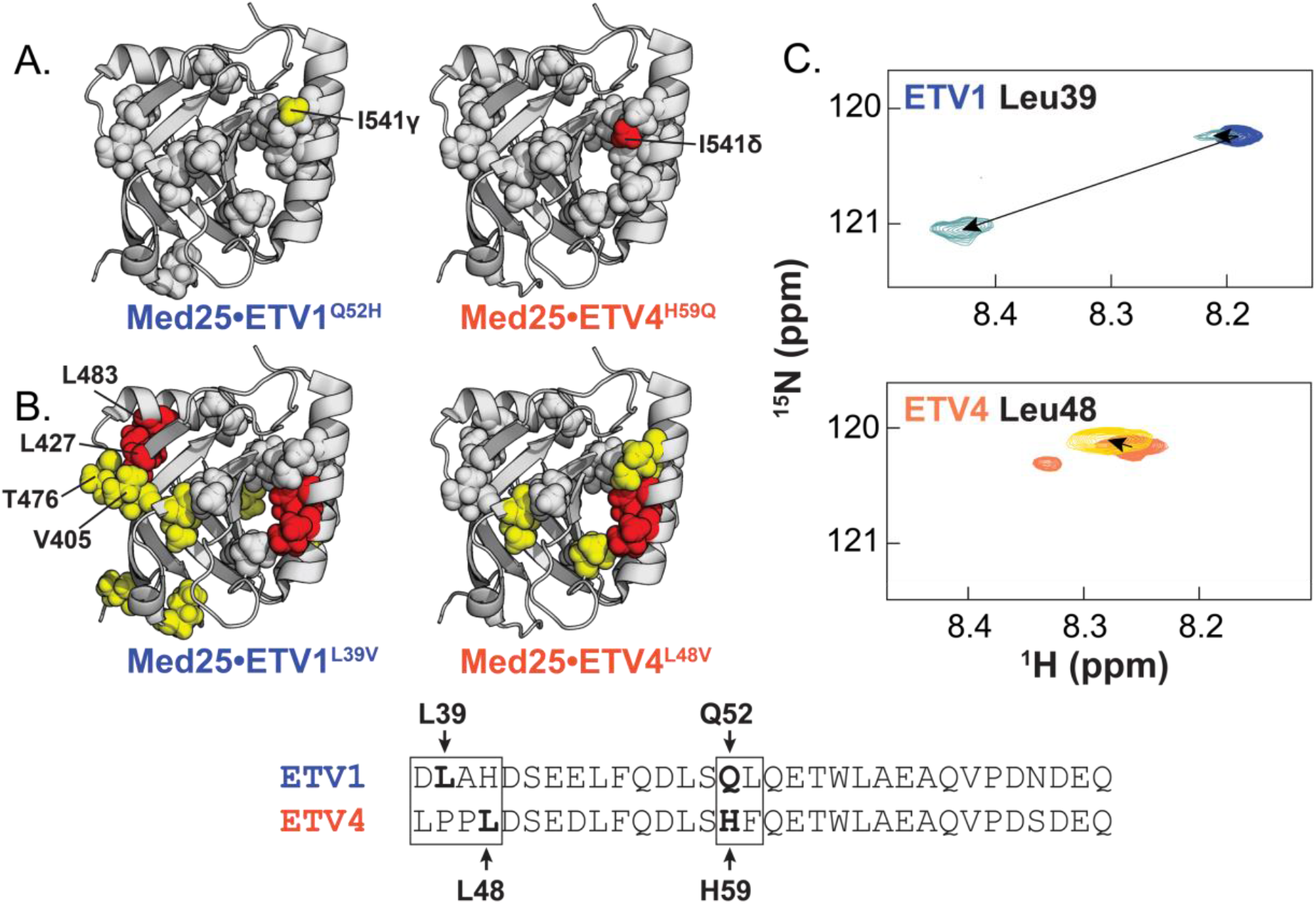
ETV/PEA3 variable regions engage in unique interactions with the Med25 surface. Effects of conservative mutations in the A) helical binding region and B) *N*-termini are plotted on the structure of Med25. Yellow = 0.015 – 0.030 ppm. Red ≥ 0.030 ppm. Residues discussed in the text are labeled. Grey spheres denote residues that undergo identical perturbations in both parent and mutant complexes. Residues chosen for mutation are bolded and labeled in the alignment. C) Chemical shift perturbations of 150 μM ETV1 (above) and ETV4 (below) TADs in the absence (blue and orange, respectively) and presence (light blue and maroon, respectively) of 280 μM unlabeled Med25. TADs were selectively _15_N labeled at the positions noted. Small secondary peaks in free ETV4 spectra were observed and likely arose from isomerization of the two tandem Pro residues in the *N*-terminal region.

Next, we introduced mutations into the disordered *N*-terminus. Specific sites for mutations in this region were not immediately obvious from previous data, so several point mutations were made; Leu to Val mutations ETV1_L39V_ and ETV4_L48V_ sufficed to produce measurable differences in bound _1_H,_13_C-HSQC spectra compared to the native TADs, without significantly altering the overall spectra (Fig. 4B, SI Figs. S22 and S23). In contrast to mutations in the helical binding region, ETV1_L39V_ and ETV4_L48V_ affected a larger overall site on the Med25 structure, indicating possible indirect effects stemming from these mutations. Indeed, both of these *N*-terminal mutants altered an overlapping sub-set of residues in the core binding site, which would be occupied by the TAD helical binding region. Nonetheless, evidence for a direct interaction of the ETV1 *N*-terminus was also apparent from the ETV1_L39V_•Med25 spectrum; several shifts from the native ETV1•Med25 spectrum were observed in a cluster of residues in a distal site involving Val405, Leu427, Thr476, and Leu483. Conversely, this site was unaffected by the ETV4_L48V_ variant, suggesting this interaction is made only by the ETV1 *N*-terminus. Consistent with a functional role for differential engagement of this distal site, our kinetics data demonstrated that removal of the ETV1 *N*-terminus significantly alters the ETV1•Med25 conformational ensemble, whereas removal of the ETV4 *N*-terminus only slightly affected the ETV4•Med25 ensemble (Fig. 3C).

To further test this model, we synthesized ETV1 and ETV4 TADs that were selectively _15_N-labeled at single Leu residues and analyzed CSPs in _1_H,_15_N-HSQC spectra upon binding of unlabeled Med25. Analysis of Leu residues in the helical binding region was attempted by this method; however, in almost all cases these peaks were too broad to detect when the TADs were bound to unlabeled Med25. In contrast, peaks for Leu residues in the *N*-terminus remained relatively sharp upon binding to Med25, likely due to these regions retaining more structural disorder in the complex.(30) Comparison of CSPs of Leu residues in the *N*-termini of ETV1 and ETV4 were consistent with the proposed differential interaction with Med25: Leu39 in the ETV1 *N*-terminus underwent a large (~0.27 ppm) shift whereas Leu48 in the ETV4 *N*-terminus shifted only slightly (0.02 ppm) upon addition of unlabeled Med25 (Fig. 4C). Interestingly, we also observed a minor peak in spectra of bound ETV1 Leu39 that was slightly shifted from the unbound position, perhaps representing one of the lowly-populated ETV1•Med25 conformations (Fig. 2D) where the *N*-terminus is weakly bound to the Med25 surface.

### ETV/PEA3•Med25 Conformational Changes Involve Shifts in Med25 Structure

Altogether, these data support a model where the conformational differences between ETV/PEA3•Med25 complexes are caused by the ETV/PEA3 variable regions engaging with the Med25 surface in unique and sequence-dependent manners. While NMR and mutagenesis data revealed that the variable *N*-terminus affects ETV/PEA3•Med25 conformation via differential engagement with the Med25 surface, the mechanism by which bound conformational behavior is affected by the variable motif in the helical binding region is unclear. Interestingly, NMR analysis demonstrated that the ETV1 and ETV4 variable motifs localize to a similar region (Fig. 4A), which indicates that conformational differences caused by this motif originate from distinct interactions in the same site. We thus reasoned that this would likely involve remodeling of the Med25 ABD. To test this hypothesis, we examined changes in the bound TAD•Med25 _1_H,_13_C-HSQC spectra produced by a point mutation in the variable motif that significantly redistributes the populations of conformational states. The ETV4_F60L_ mutation was selected because this small change in residue identity caused a drastic conformational redistribution (Fig. 3B, compare bottom left and top right) to favor the initial bound sub-state (C1).

Comparison of the _1_H,_13_C-HSQC spectra of ETV4•Med25 and ETV4_F60L_•Med25 revealed several Med25 peaks that are significantly perturbed when bound to native ETV4 shift less drastically when ETV4_F60L_ is bound (Fig. 5A). Significantly, the ETV4_F60L_ variant elicited weaker CSPs around the core binding site than native ETV4 (Fig. 5B), suggesting that several of the large CSPs in this region are tied to the conformational changes. Furthermore, this behavior was observed for peaks representing residues that are buried or in allosteric regions of the protein, including the β-barrel core, the interface between the β-barrel and the allosteric α2 helix, and the interface between the *C*-terminal α3 helix and the allosteric α1 helix (Fig. 5B). These data are therefore consistent with a direct role for ABD conformational plasticity in molecular recognition, which likely enables specific interactions by revealing new topology in the core binding site.

**Figure 5.**
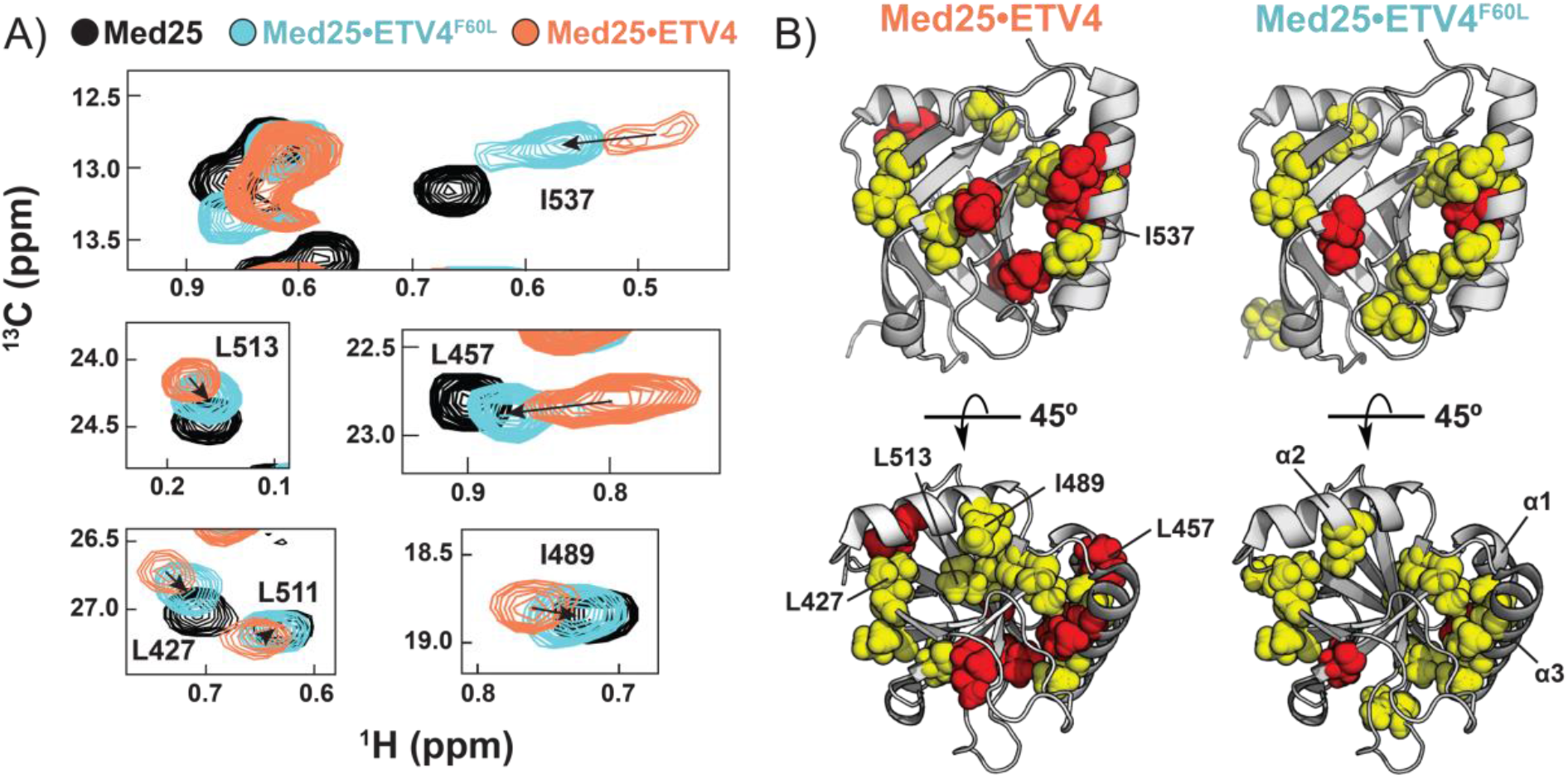
Structural shifts in the Med25 ABD accompany the conformational changes of ETV/PEA3•Med25 complexes. A) Examples of Med25 resonances undergoing shifts toward the unbound position (black) upon ETV4_F60L_ (cyan) mutation. B) Comparison of Chemical Shift Perturbations from binding of ETV4 (left) and ETV4_F60L_ (right) demonstrate that conformational changes involve the binding site and allosteric regions of Med25. Residues shown in A. are labeled on the structures. Yellow = 0.040–0.080 ppm, red > 0.080 ppm.

## Discussion

The exceptional sequence variability of functional TADs—characterized only by a general preponderance of acidic and hydrophobic amino acids—has remained a molecular recognition enigma over the past several decades.(1–5, 10, 12–14) There have been several recognition models advanced to account for the large variety of functional TADs, most of which propose that TAD•ABD recognition occurs via nonspecific intermolecular interactions.(3, 9–12) The major driving force of association in these models is thus the stochastic burial of hydrophobic sidechains rather than the formation of defined intermolecular contacts typical for well-structured PPIs. However, there is limited direct biophysical evidence for nonspecific mechanisms, and the available biophysical data relies almost entirely on a) the observation of similar CSP patterns of unique TAD sequences binding to the same ABD,(11) and b) NMR paramagnetic relaxation enhancement (PRE) patterns showing multiple bound orientations of the TAD on the ABD surface.(9–11) Alternative explanations also exist for both of these observations: similar CSP trajectories are expected if the ABD undergoes a conserved conformational change upon binding,(42–44) and significant PREs can be observed from lowly populated (<10%) sub-states that place the paramagnetic spin label in close proximity to the interacting partner.(45, 46) The degree to which activator•coactivator recognition is truly nonspecific in such examples is therefore unclear. Unfortunately, this widely accepted view of activator•coactivator recognition also represents a primary reason why these PPIs have been traditionally considered “untargetable”.(29)

Here, we scrutinized these recognition models by subjecting the dynamic PPIs formed between the ETV/PEA3 family of TADs and their binding partner Med25 to detailed mechanistic and structural dissection. We found that these interactions exhibited a striking degree of conformational sensitivity to small changes in TAD sequence, which is inconsistent with PPIs driven by nonspecific intermolecular interactions. Instead, both ordered and disordered regions of ETV/PEA3 TADs have the capacity to engage in a finite set of *specific* interactions with the Med25 surface. This recognition mechanism is further enabled by an underappreciated role of ABD plasticity in molecular recognition: in contrast to the shallow and largely featureless ETV/PEA3 binding surface presented by Med25 in the unbound state, the Med25 ABD undergoes significant remodeling upon complex formation and thus likely plays a direct role in enabling different interaction modes.(47)

Taken together, these data reveal alternative *specific* mechanisms of TAD•ABD molecular recognition that rationalize the extreme sequence variability of functional TADs (Fig. 6), which has been proposed to be due to entirely nonspecific recognition mechanisms.(3, 9–11) In contrast to nonspecific mechanisms, the model of TAD•ABD molecular recognition outlined here highlights a direct path to targeting strategies for small molecule therapeutics or probes of activator•coactivator complexes. Specifically, the role of ABD plasticity in recognition suggests that development of molecules that stabilize specific ABD conformational states may be a more effective targeting strategy than directly targeting the topologically challenging TAD•ABD interface. We previously obtained allosteric modulators of Med25 and other ABDs by covalent targeting of dynamic structural elements,(32, 48) which supports the idea that ABD plasticity can be effectively exploited by small molecules.

**Figure 6.**
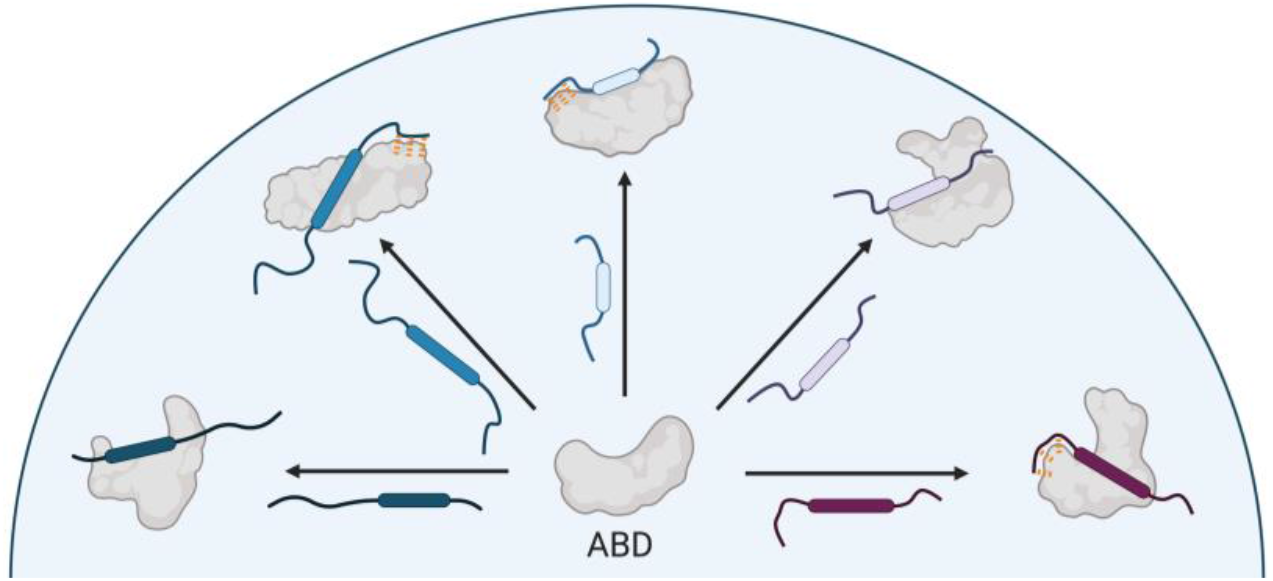
Specific recognition model emerging from this work. Activator binding domains recognize a diversity of activators via conformational plasticity and unique specific intermolecular interactions. Rounded boxes in the TADs represent the “canonical” recognition motifs. Orange dashes indicate specific 734) 975-2902 734) 975-2902interactions made by disordered regions outside the recognition motifs.

A fundamental biological question that emerges from our study is how molecular recognition mechanisms affect function. In general, it is still exceptionally challenging to relate PPI function and affinity; in processes where dynamic PPIs serve as critical functional events, such as transcription and proteostasis, there is often little or no correlation between affinity and functional activity.(49, 50) Several factors play into this observation, such as subcellular localization and concentration, but a potentially significant factor is the mechanism by which the complex forms. For example, in our current study we observed that the lifetimes of individual ETV/PEA3•Med25 conformations varied up to two orders of magnitude (SI Table S2), therefore shifting the conformational ensemble towards longer-lived sub-states could have significant functional outcomes without necessitating changes in affinity.(51) Thus, there is a critical need to reevaluate the nonspecific recognition models that are common for dynamic biomolecular interactions—such as those made by intrinsically disordered proteins and RNA(11, 52)—to develop a deeper understanding into the relationship between the function of biomolecular interactions and the mechanisms by which they are formed.

## Methods

### Protein Expression and Purification

Med25 AcID was expressed and purified from a pET21b-Med25(394-543)-His6 plasmid from *E. coli* BL21 (DE3) cells as described previously (full details in the Supporting Information).(32) Uniformly _13_C,_15_N labeled Med25 for NMR experiments was expressed identically except using M9 minimal media supplemented with 1 g/L _15_NH_4_Cl, 2 g/L _13_C_-D-_glucose, and 0.5% _13_C,_15_N-labelled Bioexpress media. Protein identity was confirmed by mass spectrometry (Agilent Q-TOF).

### Peptide Synthesis

The peptides used in this study were prepared using standard Fmoc solid-phase peptide synthesis on a Liberty Blue Microwave Peptide Synthesizer (CEM). _15_N-Leu labeled peptides were synthesized using Fmoc-_15_N-Leu in place of unlabeled Fmoc-Leu at the specified positions. Details of synthesis and peptide characterization data (HPLC) are included in the Supporting Information.

### Stopped-flow kinetics

Stopped-flow kinetic assays were performed using a Kintek SF-2001 stopped flow instrument equipped with a 100-W Xe arc lamp in two-syringe mode. All experiments were completed at 10 °C in stopped-flow buffer (10 mM sodium phosphate, 100 mM NaCl, 2% DMSO, 1% glycerol, 0.001% NP-40, pH 6.8). All concentrations reported are after mixing. The 4-DMN fluorophore was excited at 440 nm, and fluorescence intensity was measured at wavelengths >510 nm using a long-pass filter (Corion). Further details, including an explanation of how reported values were determined, are in the Supporting Information.

### NMR Spectroscopy

Constant time _1_H,_13_C-HSQC experiments were performed with uniformly _13_C,_15_N-labeled Med25 in NMR buffer (20 mM sodium phosphate pH 6.5, 150 mM NaCl, 3 mM DTT, 10% D_2_O, and 2% DMSO) on a Bruker 600 MHz instrument equipped with a cryoprobe. HSQC experiments were processed in NMRPipe(53) and visualized with NMRFAM-Sparky.(54) Chemical shift perturbation analyses were performed on samples with 1.1 equivalents of unlabeled binding partner, which results in ≥96% bound Med25 based on measured *K*_d_ values. Assignments of side-chain methyl resonances of free Med25 were achieved through 3D H(CCCO)NH and (H)CC(CO)NH TOCSY experiments (23 ms TOCSY mixing time) performed with a sample of 600 μM _13_C,_15_N Med25 on a Bruker 800 MHz instrument equipped with a cryoprobe. Further details about assignment and data analysis can be found in the Supporting Information.

## Acknowledgements

Financial support for this work was received from NIH R01 GM65530, R35 GM136356 (to A.K.M.) and CA207272 (to T.C.). M.J.H. was supported by a fellowship from the Department of Education (Graduate Assistance in Areas of National Need), and B.S.M. by a fellowship from the Michigan Life Sciences May-Walt Fellowship Fund. We thank the Mapp lab for helpful comments and D. Sahu for assistance with performing the NMR assignment experiments. Figures 1 and 6 were created using BioRender.com.

## Author Contributions

M.J.H. and A.K.M. conceived and designed research. M.J.H., B.M.L., and B.S.M. performed research. M.J.H. analyzed the data. M.J.H. and A.K.M. wrote the manuscript with input from all coauthors.

## Supporting Information for “Unexpected Specificity within Dynamic Transcriptional Protein-Protein Complexes”

### Protein Expression and Purification

Med25 AcID was expressed and purified from a pET21b-Med25(394-543)-His_6_ plasmid from E. coli BL21 (DE3) cells as described previously.(1) Briefly, 50 mL starter cultures in LB were grown overnight in the presence of 0.1 mg/mL ampicillin (Gold Bio Technology) at 37 °C at 150 rpm. The next day, 5 mL of the starter culture was used to inoculate 1 L of TB media (24 g yeast extract, 12 g tryptone, 4 mL glycerol, 100 mL 0.17 M KH_2_PO_4_/0.72 M K_2_HPO_4_, 900 mL water) with 0.1 mg/mL ampicillin, which was grown at 37 °C, 250 rpm, to an OD_600_ of 0.8. The incubator temperature was lowered to 21 °C and the culture was allowed to recover for 30 min, at which point isopropyl β-D1-thiogalactopyranoside (IPTG, Research Products International) was added to a final concentration of 500 μM to induce expression. The protein was allowed to express overnight (~18 hr), after which the cells were harvested via centrifugation (6000 rpm, 20 min), and then frozen and stored at −80 °C. Uniformly ^13^C,^15^N labeled Med25 for NMR experiments was expressed identically except using M9 minimal media supplemented with 1 g/L ^15^NH_4_Cl, 2 g/L ^13^C-_D_-glucose, and 0.5% ^13^C,^15^N-labelled Bioexpress media for the 1 L growth (all labeled components were purchased from Cambridge Isotopes).

To purify Med25, cell pellets were resuspended in 25 mL lysis buffer (50 mM phosphate, 300 mM NaCl, 10 mM imidazole, pH 7.2, 1.4 μL/mL β-mercaptoethanol, 1 Roche complete mini protease inhibitor tablet) and lysed by sonication. Insoluble material was then pelleted by centrifugation (9500 rpm, 20 min), the supernatant was removed and re-sonicated, and then filtered using a 0.45 syringe filter (CellTreat) and loaded onto an AKTA Pure FPLC equipped with a Ni HisTrap HP column (GE Healthcare) pre-equilibrated with wash buffer (50 mM phosphate, 300 mM NaCl, 10 mM imidazole, pH 7.2). Med25 was then purified using a gradient of 10–300 mM imidazole (other buffer components were constant), and fractions containing Med25 were pooled and subjected to secondary purification using a HiTrap SP HP cation exchange column (GE Healthcare) using a gradient of 0–1 M NaCl (50 mM sodium phosphate, 1 mM DTT, pH 6.8). Pooled fractions were dialyzed into stopped-flow buffer (10 mM sodium phosphate, 100 mM NaCl, 1% glycerol, 0.001% NP-40, pH 6.8) or NMR buffer (20 mM sodium phosphate, 150 mM NaCl, pH 6.5). Concentration was determined by a NanoDrop instrument using an extinction coefficient at 280 nm of 22,460 M^−1^cm^−1^. Aliquots were flash frozen in liquid N_2_ and stored at −80 °C until use. Protein identity was confirmed by mass spectrometry (Agilent Q-TOF).

### Peptide Synthesis

The peptides used in this study were prepared using standard Fmoc solid-phase peptide synthesis on a Liberty Blue Microwave Peptide Synthesizer (CEM). Deprotection was accomplished by 20% piperidine (ChemImpex) in DMF supplemented with 0.2 M Oxyma Pure (CEM), with irradiation at 90 °C for 1 min. Coupling reactions were completed with 5 equivalents of Fmoc-amino acid (CEM), 7 equivalents of diisopropylcarbodiimide (ChemImpex), and 5 equivalents of Oxyma Pure in DMF, with irradiation at 90 °C for 4 min. Between steps, the resin was rinsed four times with an excess of DMF. Selectively ^15^N-labeled peptides were synthesized using Fmoc-^15^N-Leu in place of Fmoc-Leu in the specified positions.

Unlabeled peptides were acetylated at the *N*-terminus through reaction of a mixture of acetic anhydride (Sigma), triethylamine (Sigma), and dichloromethane in a 1:1:8 ratio for 30 min after the conclusion of the synthesis. 4-DMN labeled peptides were coupled with ~1.5 equivalents of 4-DMN-β-Alanine, as described previously.(1) All peptides were cleaved from the resin using a cocktail of 95:2.5:2.5 trifluoroacetic acid (Sigma), ethanedithiol (Sigma), and water for 3 hr, followed by filtration. The peptide solution was concentrated under a stream of N_2_, then precipitated with cold diethyl ether and pelleted by centrifugation (4,500 rpm, 5 min). The ether was then discarded and the pellet was taken up in 7:3 100 mM Ammonium acetate-acetonitrile. The peptide was purified by an Agilent 1260 preparatory HPLC using a 40 minute gradient of 10-50% acetonitrile, with 100 mM ammonium acetate as the stationary phase. The flow rate was 40 mL/min. Fractions containing the correct peptide were pooled and lyophilized, and the resulting powders were dissolved in minimal DMSO and stored at −20 °C. Concentrations were taken with a 1:100 dilution of the DMSO stock into 6 M Guanidinium Chloride on a NanoDrop instrument, using extinction coefficients of 5,690 M^−1^cm^−1^ (280 nm, unlabeled) or 10,800 M^−1^cm^−1^ (450 nm, 4-DMN-labeled). Identity of the peptides were confirmed by mass spectrometry.

**Table S1.**
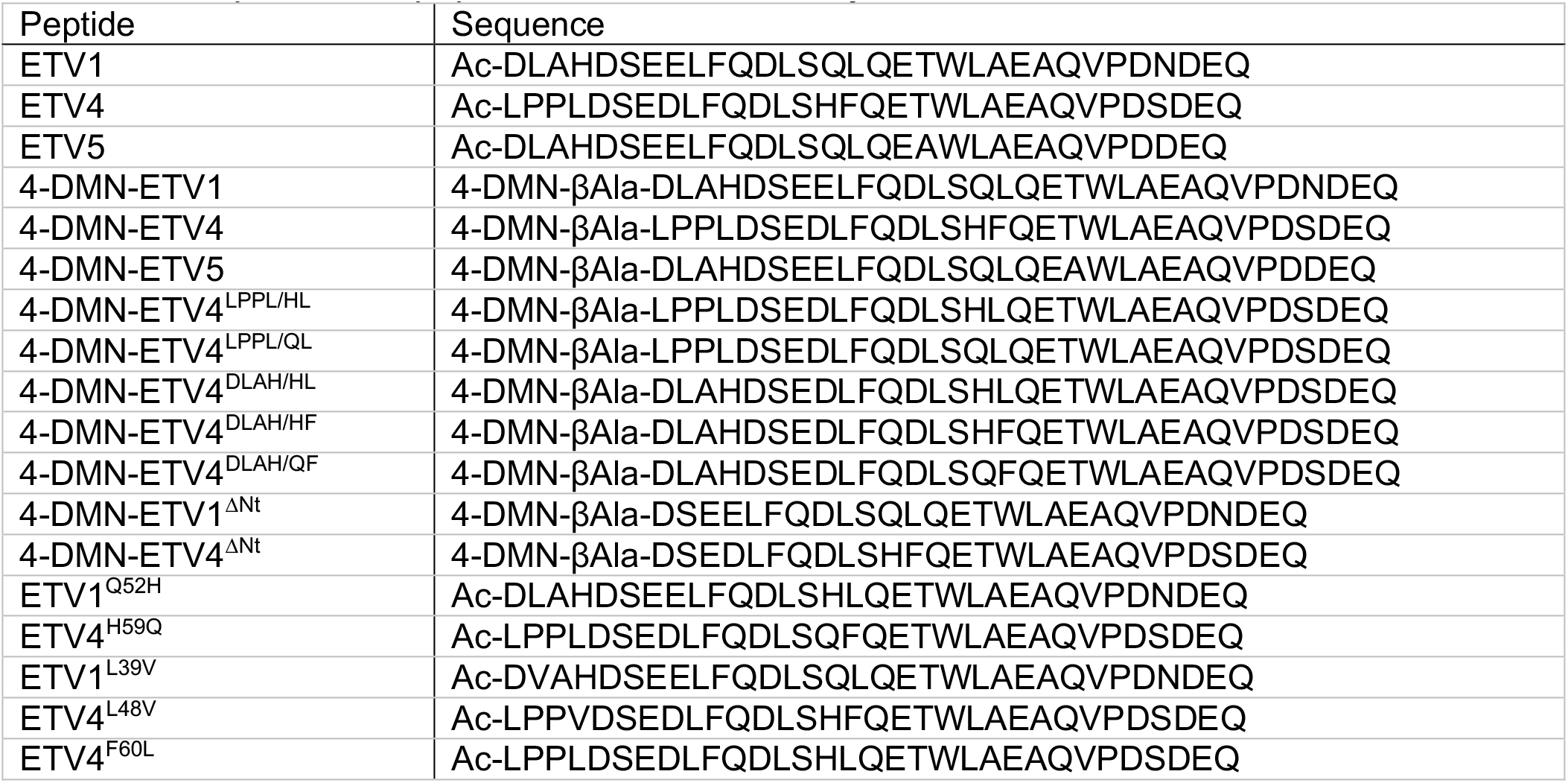
Sequences of peptides used for this study

| Peptide | Sequence |
| --- | --- |
| ETV1 | Ac-DLAHDSEELFQDLSQLQETWLAEAQVPDNDEQ |
| ETV4 | Ac-LPPLDSEDLFQDLSHFQETWLAEAQVPDSDEQ |
| ETV5 | Ac-DLAHDSEELFQDLSQLQEAWLAEAQVPDDEQ |
| 4-DMN-ETV1 | 4-DMN-βAla-DLAHDSEELFQDLSQLQETWLAEAQVPDNDEQ |
| 4-DMN-ETV4 | 4-DMN-βAla-LPPLDSEDLFQDLSHFQETWLAEAQVPDSDEQ |
| 4-DMN-ETV5 | 4-DMN-βAla-DLAHDSEELFQDLSQLQEAWLAEAQVPDDEQ |
| 4-DMN-ETV4<sup>LPPL/HL</sup> | 4-DMN-βAla-LPPLDSEDLFQDLSHLQETWLAEAQVPDSDEQ |
| 4-DMN-ETV4<sup>LPPL/QL</sup> | 4-DMN-βAla-LPPLDSEDLFQDLSQLQETWLAEAQVPDSDEQ |
| 4-DMN-ETV4<sup>DLAH/HL</sup> | 4-DMN-βAla-DLAHDSEDLFQDLSHLQETWLAEAQVPDSDEQ |
| 4-DMN-ETV4<sup>DLAH/HF</sup> | 4-DMN-βAla-DLAHDSEDLFQDLSHFQETWLAEAQVPDSDEQ |
| 4-DMN-ETV4<sup>DLAH/QF</sup> | 4-DMN-βAla-DLAHDSEDLFQDLSQFQETWLAEAQVPDSDEQ |
| 4-DMN-ETV1<sup>ΔNt</sup> | 4-DMN-βAla-DSEELFQDLSQLQETWLAEAQVPDNDEQ |
| 4-DMN-ETV4<sup>ΔNt</sup> | 4-DMN-βAla-DSEDLFQDLSHFQETWLAEAQVPDSDEQ |
| ETV1<sup>Q52H</sup> | Ac-DLAHDSEELFQDLSHLQETWLAEAQVPDNDEQ |
| ETV4<sup>H59Q</sup> | Ac-LPPLDSEDLFQDLSQFQETWLAEAQVPDSDEQ |
| ETV1<sup>L39V</sup> | Ac-DVAHDSEELFQDLSQLQETWLAEAQVPDNDEQ |
| ETV4<sup>L48V</sup> | Ac-LPPVDSEDLFQDLSHFQETWLAEAQVPDSDEQ |
| ETV4<sup>F60L</sup> | Ac-LPPLDSEDLFQDLSHLQETWLAEAQVPDSDEQ |

**Figure S1.**
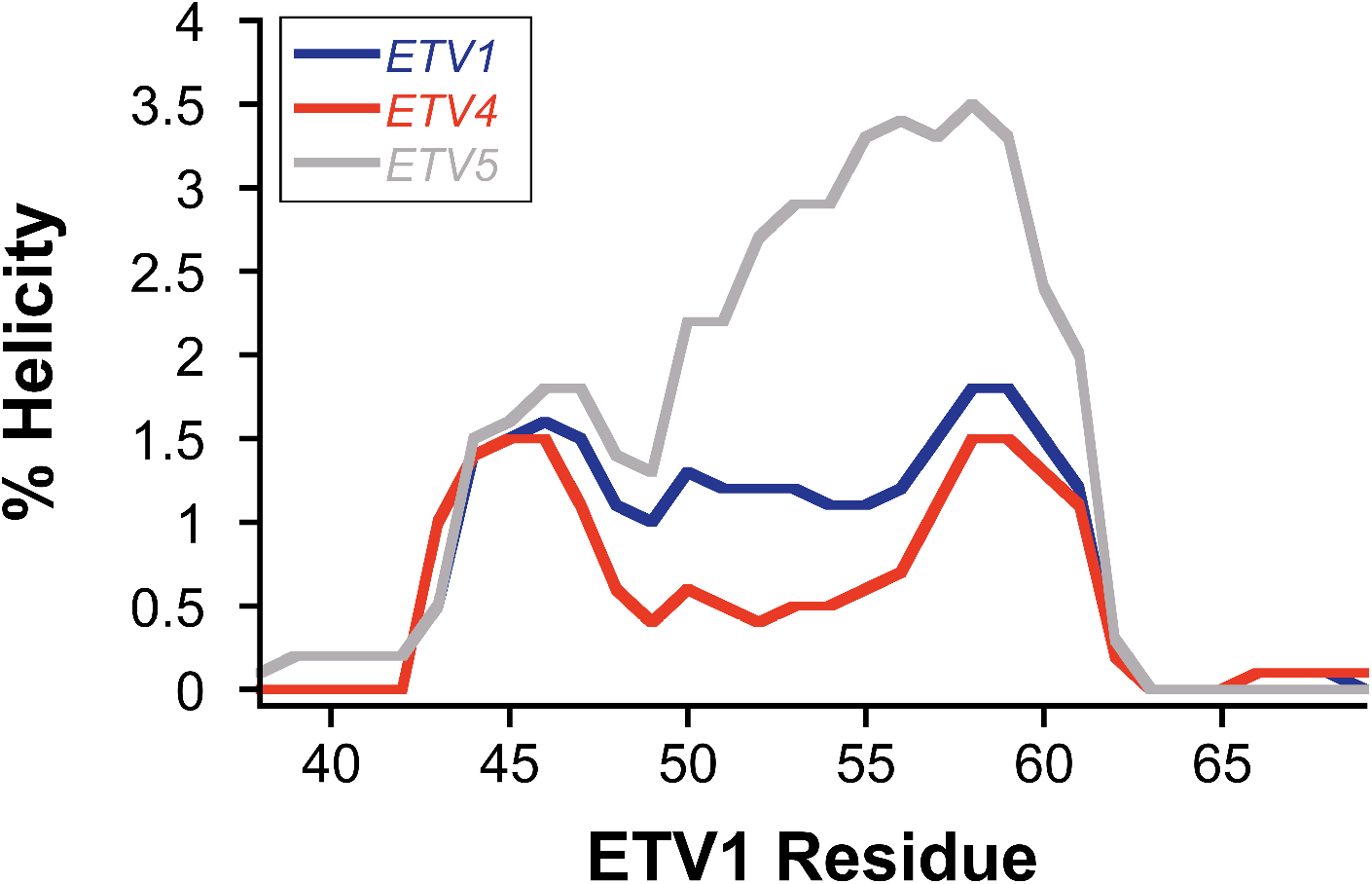
Predicted helical propensity of PEA3 activators from Agadir,(2) aligned to ETV1 sequence.

### Stopped-flow kinetics

Stopped-flow kinetic assays were performed using a Kintek SF-2001 stopped flow instrument equipped with a 100-W Xe arc lamp in two-syringe mode. All experiments were completed at 10 °C in stopped-flow buffer (10 mM sodium phosphate, 100 mM NaCl, 2% DMSO, 1% glycerol, 0.001% NP-40, pH 6.8). All concentrations reported are after mixing. The 4-DMN fluorophore was excited at 440 nm, and fluorescence intensity was measured at wavelengths >510 nm using a long-pass filter (Corion). Association experiments were completed by 1:1 mixing of a constant concentration of 0.25 μM 4-DMN-labeled peptide with variable concentrations of Med25. Dissociation experiments were performed by mixing 50 μM unlabeled peptide with a preformed complex of 0.5-1 μM Med25 and 0.25 μM labeled peptide. Unlabeled peptides for dissociation experiments mutants were typically the parent peptide, but no unique effects were observed from using different competitors. Typically, 30-40 traces were averaged before fitting.

Traces were fitted using a series of exponential equations (first equation below), where *F(t)* is the fluorescence at time *t*, *F*_∞_ is the endpoint fluorescence, Δ*F*_*n*_ are the fluorescence amplitudes, and *k*_obs,n_ are the observed rate constants. Equilibrium dissociation constants (*K*_d_) were determined by fitting the concentration dependence of *F*_∞_ values to a standard hyperbolic equation. The individual *k*_obs,n_ values were plotted as a function of concentration and fit to square hyperbola (second equation below) to determine the maximal observed rate constant (*k*_obs,n,max_), and the half maximal concentration (*K*_1/2,n_). The value of *k*_obs,n,min_ was included for fitting purposes, but the value itself is defined by the corresponding *k*_obs,n,off_ value from dissociation experiments and thus the *k*_obs,n,min_ value from fitting was not used for calculations. The microscopic rate constants were calculated using a combined rapid equilibrium and steady-state approximation, detailed in the next section. This approach was enabled by optimized conditions for dissociation experiments from our previous report,(1) as the *k*_obs,n,off_ phases were more clearly defined under the conditions used. Values of all microscopic rate and equilibrium constants were calculated for single datasets and averaged across 2-3 results from independent datasets. All errors reported are the standard deviation between the results from separate datasets. Examples of datasets for all mutants are reported at the end of this document.

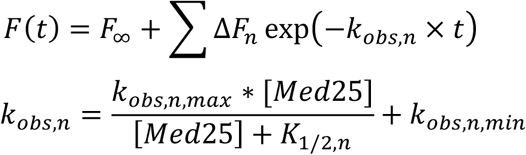

### Calculation of Microscopic Rate Constants

**Figure S2.**
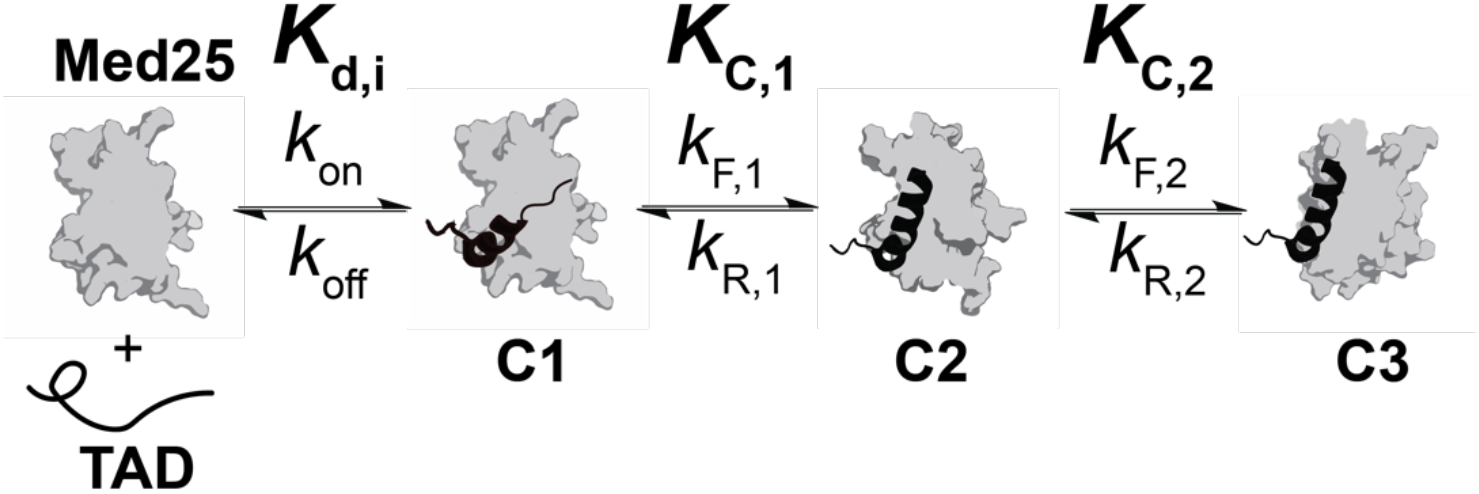
Kinetic mechanism of Med25 binding to PEA3 TADs, as determined previously.(1) Microscopic rate and equilibrium constants are labeled as they appear in this supporting information document.

Here, we chose to use a combined rapid equilibrium and steady state approach to determine rate parameters as a straightforward way to handle the large amount of complex kinetic data collected. This method for calculating the microscopic rate constants is split into two “sections”. First, the mechanism is considered as only the first two steps: the initial binding step to form conformer C1 and its transition to C2. After calculation of all first order microscopic rate constants from this “section” of the mechanism, the transition from C2 to C3 is considered. This is similar to treating the first two steps of the binding mechanism as a rapid equilibrium before the final conformational transition to C3. For all mechanisms where only a single conformational change was observed, the calculations followed the same procedure except that the final step to calculate rate constants for the C2–C3 transition was left out.

Beginning from the first two steps, the maximal observed rate constant of the first conformational change phase (*k*_obs,2,max_) is:

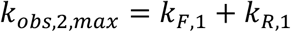

And, by the steady-state approximation, the corresponding observed rate constant for dissociation (*k*_obs,2,off_) is:

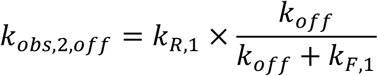

The value of *k*_off_ can be retrieved from the observed rate constants from dissociation experiments (*k*_obs,n,off_) in an analogous way to our previous work (1). That is, by the exact expression for a two step induced fit mechanism, *k*_off_ is equivalent to:

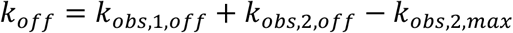

In all cases except for ETV4, the fast dissociation phase (*k*_obs,1,off_) is well defined in dissociation experiments, enabling the use of this method. For ETV4, where it is not well defined, this value was set to the minimal value we observed for other variants tested, 300 s^−1^. To substitute a directly measurable value for *k*_F,1_, the equation defined above for *k*_obs,2,max_ can be used. Thus, by substitution and simplification:

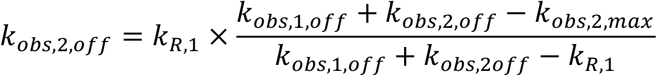

And by rearrangement:

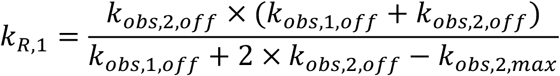

Calculation of *k*_F,1_ is then obtained by subtracting the *k*_R,1_ value from *k*_obs,2,max_. Next, the transition from C2 to C3 was considered. In dissociation experiments, the observed rate constant of the kinetic phase corresponding to this step (*k*_obs,3,off_) is given below by the steady-state approximation:

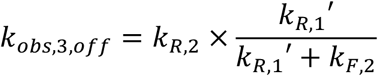

Here, *k*_R,1_’ is the net rate constant for successful dissociation from the C2 conformer and is identical to the expression for *k*_obs,2,off_ above. Thus, the value of *k*_obs,2,off_ that is obtained from fitting is used in this calculation. Similarly, *k*_obs,3,max_ is the sum of *k*_F,2_ and *k*_R,2_. Thus, substitution in a analogous manner as above gives:

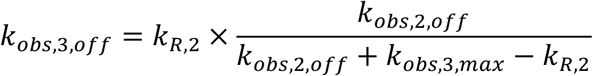

And by rearrangement:

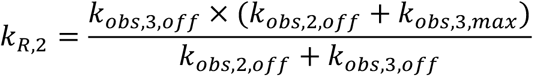

Again, *k*_F,2_ is obtained by subtracting the calculated value of *k*_R,2_ from *k*_obs,3,max_. The calculated values for all tested variants in this study are shown in Table S2. Relative populations of C1, C2, and C3 at equilibrium were then determined by the conformational equilibrium constants (*K*_C,n_ = *k*_F,n_/*k*_R,n_), which by definition are ratios between the conformational states. Below are the equations used to calculate the population of each conformational state, using C1 as the reference state. Only the relative conformational populations of the bound state were considered, thus the values are concentration-independent.

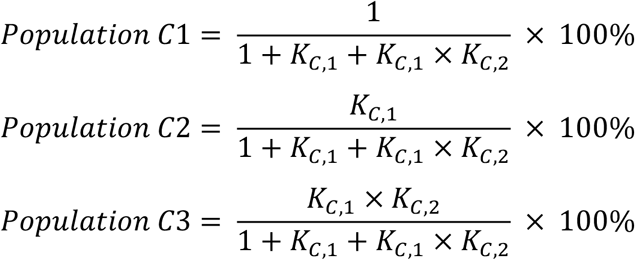

**Table S2.** Calculated rate and equilibrium constants for all 4-DMN-labeled PEA3 TADs

| ETV Variant | $K_{C,1}$ | $k_{F,1}$<br>(s<sup>-1</sup>) | $k_{R,1}$<br>(s<sup>-1</sup>) | $K_{C,2}$ | $k_{F,2}$<br>(s<sup>-1</sup>) | $k_{R,2}$<br>(s<sup>-1</sup>) | $K_d$<br>(μM) |
| --- | --- | --- | --- | --- | --- | --- | --- |
| ETV1 | 3.6 ± 0.7 | 52 ± 5 | 15 ± 3 | 0.5 ± 0.1 | 2.4 ± 0.3 | 4.9 ± 0.8 | 1.2 ± 0.2 |
| ETV4 | 3.2 ± 0.5 | 56 ± 8 | 18 ± 1 | 2.3 ± 0.5 | 3.3 ± 0.7 | 1.4 ± 0.1 | 0.7 ± 0.2 |
| ETV5 | 3.4 ± 0.6 | 46 ± 4 | 14 ± 2 | 0.6 ± 0.3 | 2.5 ± 1.0 | 4.1 ± 0.3 | 0.9 ± 0.1 |
| ETV4<sup>DLAH/HL</sup> | 2.7 ± 0.4 | 101 ± 7 | 38 ± 3 | 0.6 ± 0.1 | 4.6 ± 0.5 | 7.8 ± 0.1 | 1.2 ± 0.2 |
| ETV4<sup>DLAH/QF</sup> | 5.4 ± 0.5 | 54 ± 6 | 10 ± 0.3 | – | – | – | 0.9 ± 0.1 |
| ETV4<sup>DLAH/HF</sup> | 4.2 ± 0.2 | 63 ± 8 | 15 ± 2 | – | – | – | 0.8 ± 0.3 |
| ETV4<sup>LPPL/QL</sup> | 0.7 ± 0.1 | 27 ± 4 | 37 ± 1 | – | – | – | 2.0 ± 0.1 |
| ETV4<sup>LPPL/HL</sup> | 0.9 ± 0.1 | 34 ± 2 | 38 ± 1 | – | – | – | 1.7 ± 0.1 |
| ETV4<sup>LPPL/QF</sup> | 2.8 ± 0.4 | 52 ± 1 | 13 ± 1 | 1.6 ± 0.2 | 13 ± 0.7 | 8.1 ± 0.8 | 0.6 ± 0.1 |
| ETV1<sup>ΔNt</sup> | 1.0 ± 0.3 | 21 ± 3 | 21 ± 3 | – | – | – | 0.7 ± 0.1 |
| ETV4<sup>ΔNt</sup> | 3.9 ± 0.7 | 62 ± 1 | 16 ± 3 | 4.9 ± 1.5 | 4.9 ± 0.9 | 1.0 ± 0.1 | 0.4 ± 0.1 |

**Table S3.** Calculated conformational populations for all 4-DMN-labeled PEA3 TADs

| ETV Variant | C1 Population (%) | C2 Population (%) | C3 Population (%) |
| --- | --- | --- | --- |
| ETV1 | 16 ± 2 | 56 ± 6 | 28 ± 5 |
| ETV4 | 9 ± 1 | 28 ± 4 | 63 ± 5 |
| ETV5 | 16 ± 2 | 53 ± 8 | 31 ± 9 |
| ETV4<sup>DLAH/HL</sup> | 19 ± 2 | 51 ± 3 | 30 ± 2 |
| ETV4<sup>DLAH/QF</sup> | 16 ± 1 | 84 ± 1 | – |
| ETV4<sup>DLAH/HF</sup> | 19 ± 1 | 81 ± 1 | – |
| ETV4<sup>LPPL/QL</sup> | 58 ± 4 | 42 ± 4 | – |
| ETV4<sup>LPPL/HL</sup> | 53 ± 1 | 47 ± 1 | – |
| ETV4<sup>LPPL/QF</sup> | 12 ± 1 | 34 ± 3 | 54 ± 3 |
| ETV1<sup>ΔNt</sup> | 50 ± 7 | 50 ± 7 | – |
| ETV4<sup>ΔNt</sup> | 4 ± 2 | 17 ± 4 | 79 ± 6 |

### Discussion of Kinetic Data Analysis

The major strengths of the calculation method outlined above is that it is operationally simple, and that it performs very well when tested against simulated data in the relative rate ranges we observed experimentally. Specifically, calculated values of microscopic rate constants obtained from simulated data were well within 20% of the input values, which is on the level of normal experimental error. However, we must note that this methodology performs poorly under conditions where *k*_R,2_ approaches the value of *k*_R,1_. In kinetic simulations, the range where this becomes problematic is when *k*_R,2_ ≥ 0.5\**k*_R,1_; above this value of *k*_R,2_ the equilibrium constant *K*_C,2_ can be significantly underestimated. In the current study, this scenario only occurred for the variant ETV4^LPPL/QF^ (see Tables S2 and S3), and this likely led to an underestimated *K*_C,2_ value. Nonetheless, this variant was still detected by kinetic analysis as conformationally similar to wild-type ETV4, which is further supported by NMR analysis (see Fig. 3A in main text). As the value of *k*_R,2_ increases beyond *k*_R,1_ it becomes especially challenging to fit all three phases accurately, in this case because the amplitude of the corresponding kinetic phase shrinks considerably. This specific issue is not necessarily a drawback of our analysis approach, as other calculation methods and global fitting strategies cannot typically detect a fast step that follows a slow step. This type of kinetic behavior also may be why the some variants only have one observable conformational change. In the case where the second conformational change phase becomes undetectable due to an increase in k_R,2_, the apparent population of C2 (as presented in the figures) would be the sum of the true populations of C2 and C3 conformers. We note that this possibility does not affect any of the conclusions presented in this work.

### NMR Spectroscopy

Constant time _1_H,^13^C-HSQC experiments were performed with 50-75 μM uniformly ^13^C,^15^N-labeled Med25 in NMR buffer (20 mM sodium phosphate pH 6.5, 150 mM NaCl, 3 mM DTT, 10% D_2_O, and 2% DMSO) on a Bruker 600 MHz instrument equipped with a cryoprobe. HSQC experiments were processed in NMRPipe(3) and visualized with NMRFAM-Sparky.(4) All chemical shift perturbation analyses were performed on samples with 1.1 equivalents of unlabeled binding partner, which results in ≥96% bound Med25 based on measured *K*_d_ values. Peak assignments of PEA3-bound complexes were achieved by titration experiments with the unlabeled TAD with titration points of 0.2, 0.5, 0.8, and 1.1 equivalents of TAD, and assignments of mutations were made based on the parent TAD complexes. Native ETV1- and ETV4-bound spectra were representative of at least three biological replicates using different protein and peptide stocks. Chemical shift perturbations (Δδ) were calculated from the proton (Δδ_H_) and carbon (Δδ_C_) chemical shifts by:

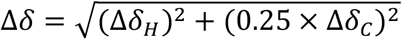

Assignments of side-chain methyl resonances of free Med25 were achieved through 3D H(CCCO)NH and (H)CC(CO)NH TOCSY experiments (23 ms TOCSY mixing time) performed with a sample of 600 μM ^13^C,^15^N Med25 on a Bruker 800 MHz instrument equipped with a cryoprobe. An additional non-uniformly sampled 4D HCC(CO)NH TOCSY experiment(5) (12 ms TOCSY mixing time and 25% sampling) with a sample of 400 μM ^13^C,^15^N Med25 was performed to supplement the 3D experiments. _1_H,^13^C resonance assignment from these experiments was enabled by a previous assignment of the Med25 _1_H,^15^N-HSQC spectrum,(1) however of 141 possible assignable non-proline amide resonances only 117 had been assigned, which precluded full assignment of the methyl spectrum. Thus, a supplemental set of TROSY-based HNCACB and CBCA(CO)NH experiments were performed to enable more complete assignment of _1_H,^15^N resonances using a 600 μM sample of ^13^C,^15^N Med25 on a Bruker 800 MHz instrument. From these experiments, 132 of 141 amide resonances along with 83 of 89 methyl resonances were assigned. Unassigned methyl resonances include one diastereotopic methyl of Leu399 and both methyls of Leu544, the latter of which is a cloning artifact and not part of the native Med25 sequence. Three residues, Leu427, Ile453, and Ile526 are not possible to assign from the TOCSY experiments as they are directly before Pro residues in the primary sequence. However, at least one methyl from each of these residues were previously assigned by another group(6) and were well separated from other resonances. The full _1_H,^13^C-HSQC spectral assignment is shown below (Fig. S3) along with a structural representation of the assignment (Fig. S4).

**Figure S3.**
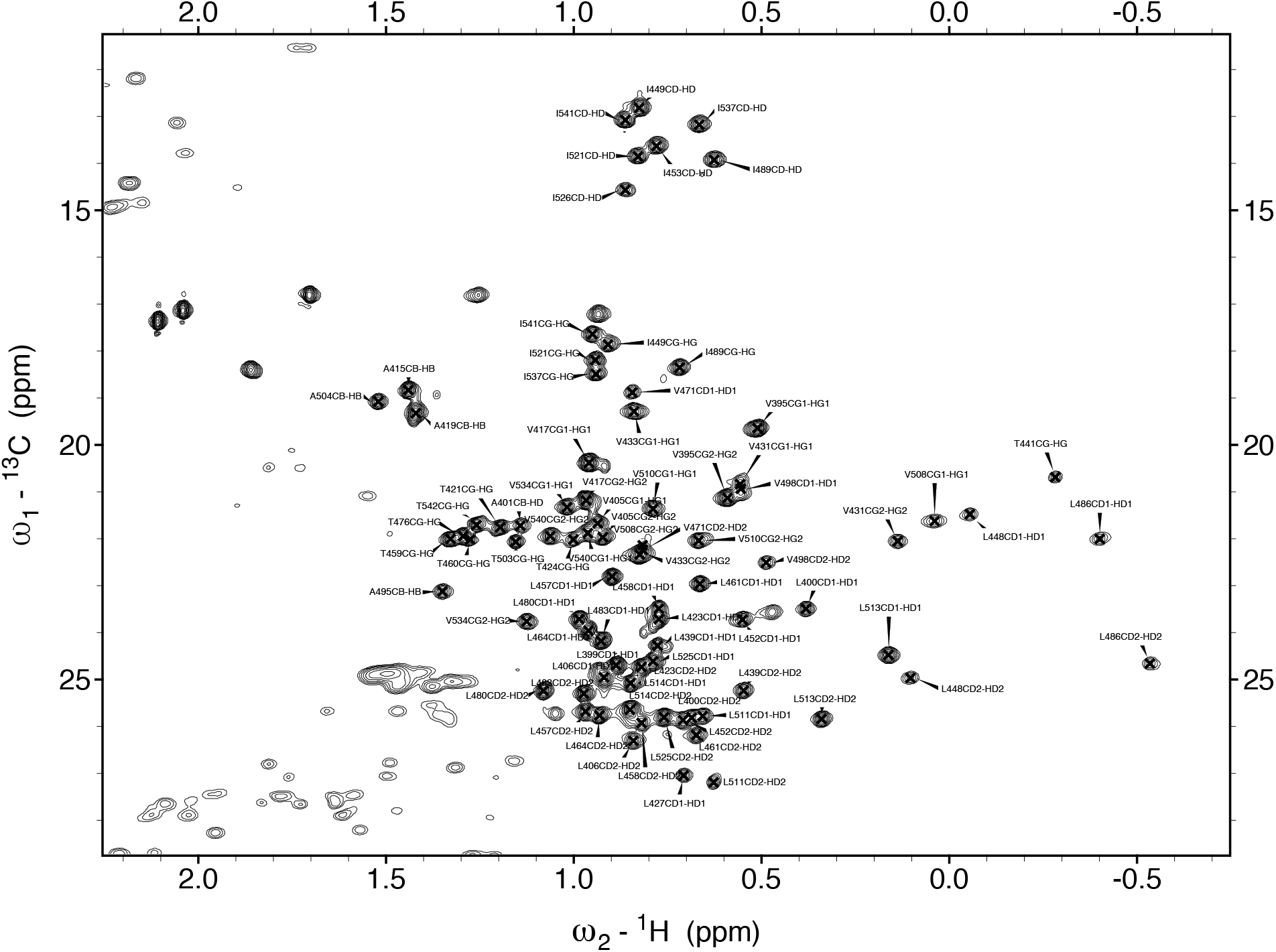
Assigned methyl ^1^H,^13^C-HSQC spectrum of Med25. Experiment was conducted at 600 μM Med25 and 800 MHz field strength.

**Figure S4.**
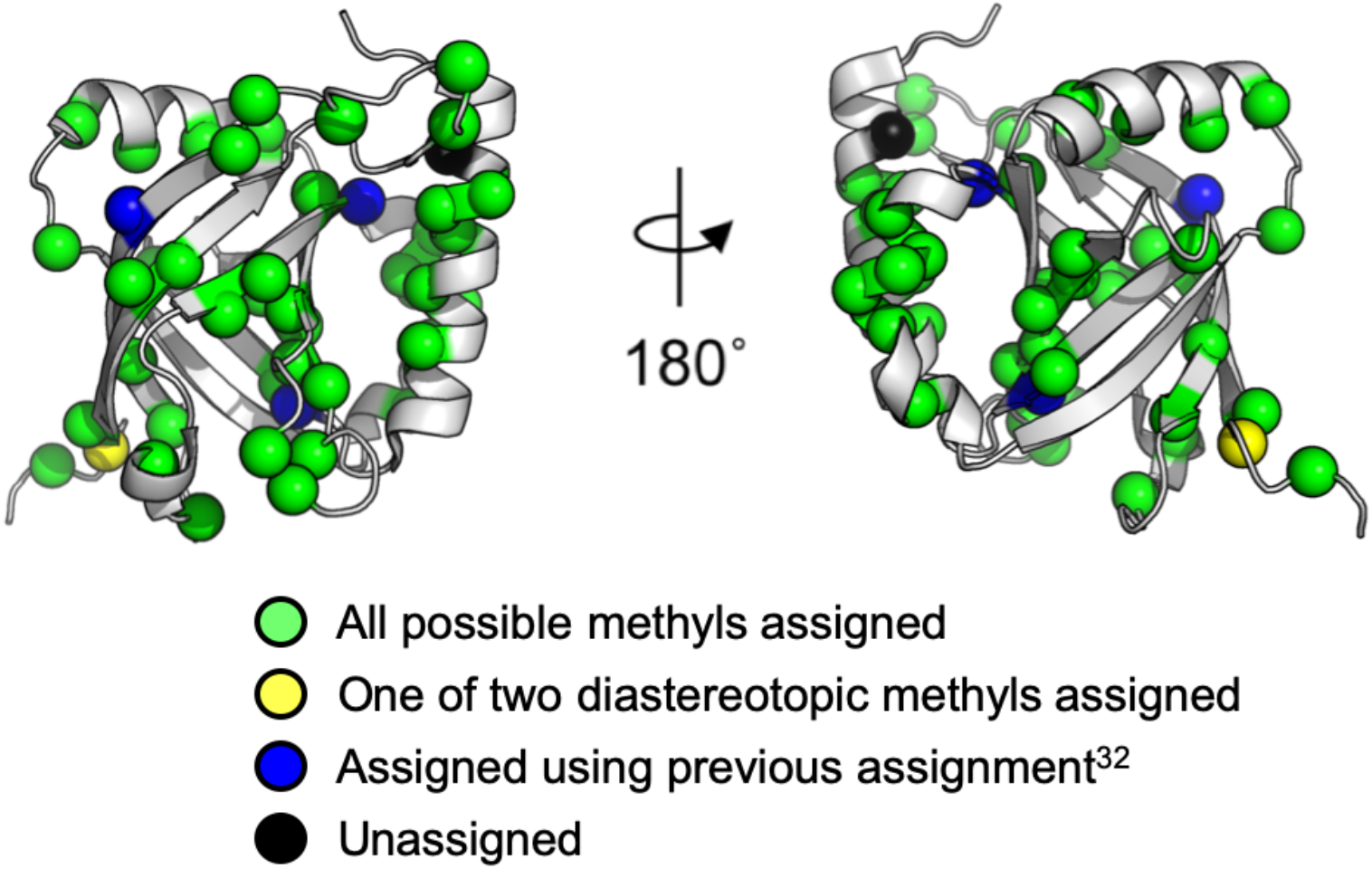
Assigned residues with methyl groups plotted on structure of Med25 (PDB ID 2XNF).(7) Green: all possible methyl groups for the residue were assigned using either the two 3D HCC(CO)NH TOCSY experiments, or the HNCACB experiment (for remaining Ala residues before Pro in the sequence). Yellow: only one of two diastereotopic methyls were assigned from the TOCSY experiments. Blue: well-dispersed methyls for the residue were assigned using a previous methyl assignment.*(6)* In all cases, these residues were before Pro residues in the sequence and thus were not observed in the HCC(CO)NH TOCSY experiments. Black: unassigned.

### Kinetic Data

The full kinetic datasets used to generate the figures in the manuscript are shown below

**Figure S5.**
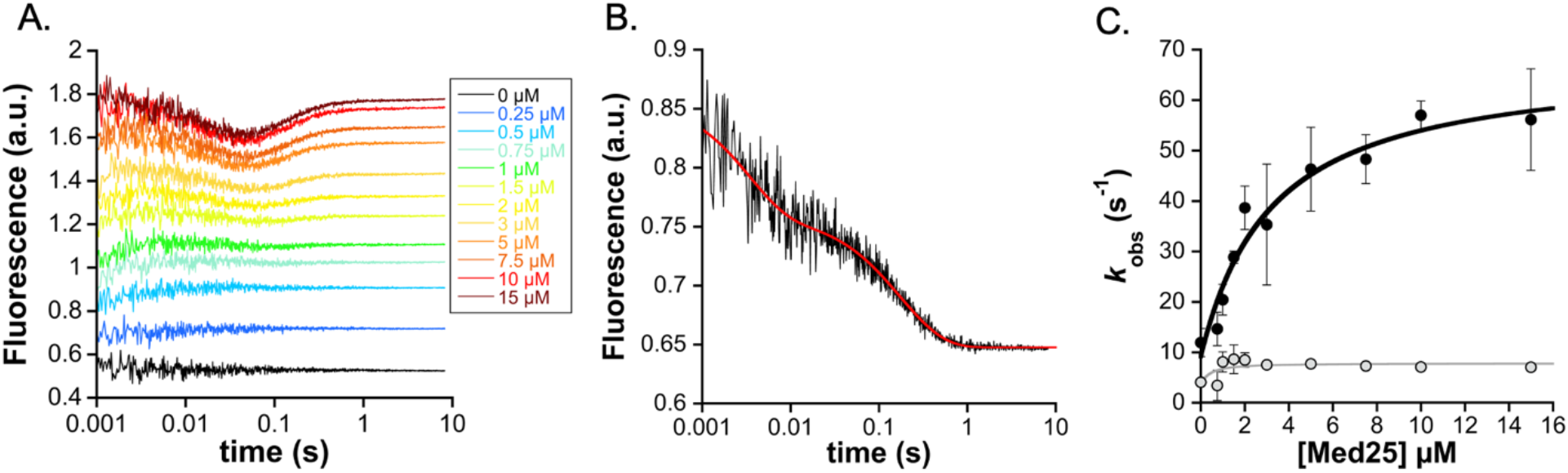
A. Association traces for binding of 0.25 μM 4-DMN-ETV1(38-69) to Med25. Concentrations of Med25 are as noted. B. Dissociation trace from 0.25 μM 4-DMN-ETV1(38-69) prebound to 0.5 μM Med25, mixed with 50 μM unlabeled ETV1(38-69). C. Observed rate constants for conformational change phases from curve fitting (black=fast phase, grey=slow phase). Values are average of *n*=3 biological replicates, and error represents the standard deviation.

**Figure S6.**
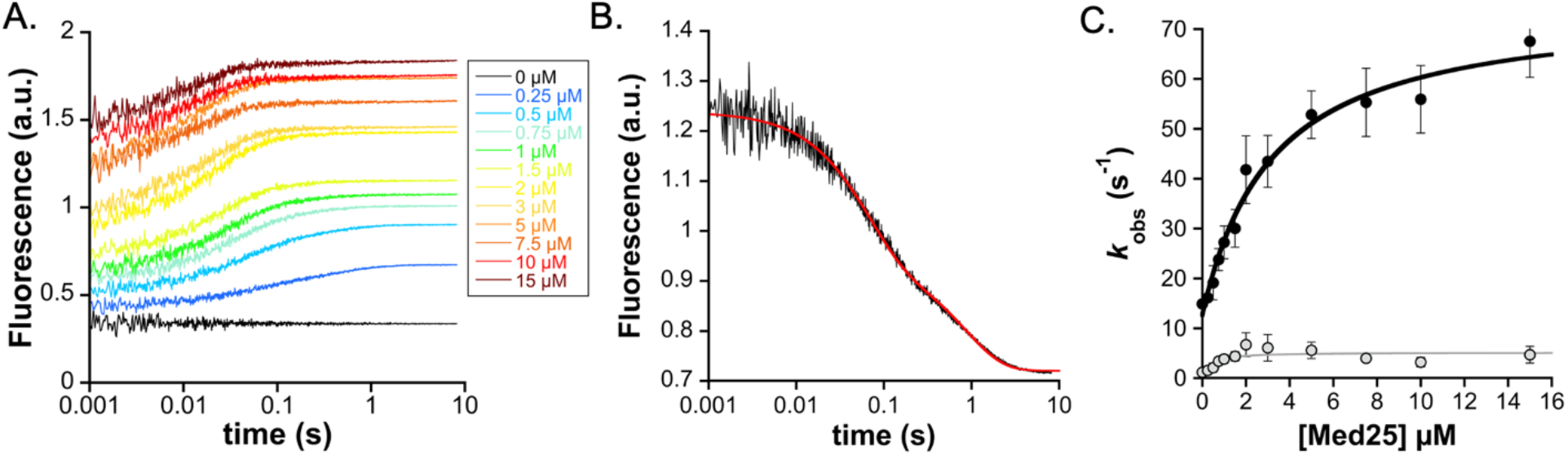
A. Association traces for binding of 0.25 μM 4-DMN-ETV4(45-76) to Med25. Concentrations of Med25 are as noted. B. Dissociation trace from 0.25 μM 4-DMN-ETV4(45-76) prebound to 0.5 μM Med25, mixed with 50 μM unlabeled ETV4(45-76). C. Observed rate constants for conformational change phases from curve fitting (black=fast phase, grey=slow phase). Values are average of *n*=4 biological replicates, and error represents the standard deviation.

**Figure S7.**
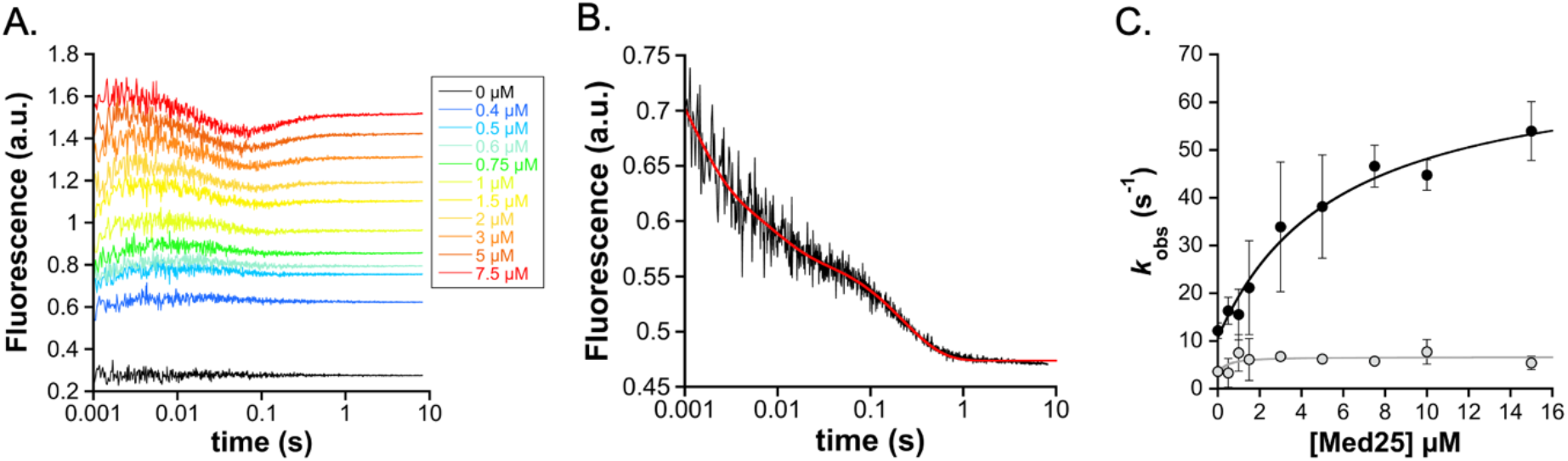
A. Association traces for binding of 0.25 μM 4-DMN-ETV5(38-68) to Med25. Concentrations of Med25 are as noted. B. Dissociation trace from 0.25 μM 4-DMN-ETV5(38-68) prebound to 0.5 μM Med25, mixed with 50 μM unlabeled ETV5(38-68). C. Observed rate constants for conformational change phases from curve fitting (black=fast phase, grey=slow phase). Values are average of *n*=3 biological replicates, and error represents the standard deviation.

**Figure S8.**
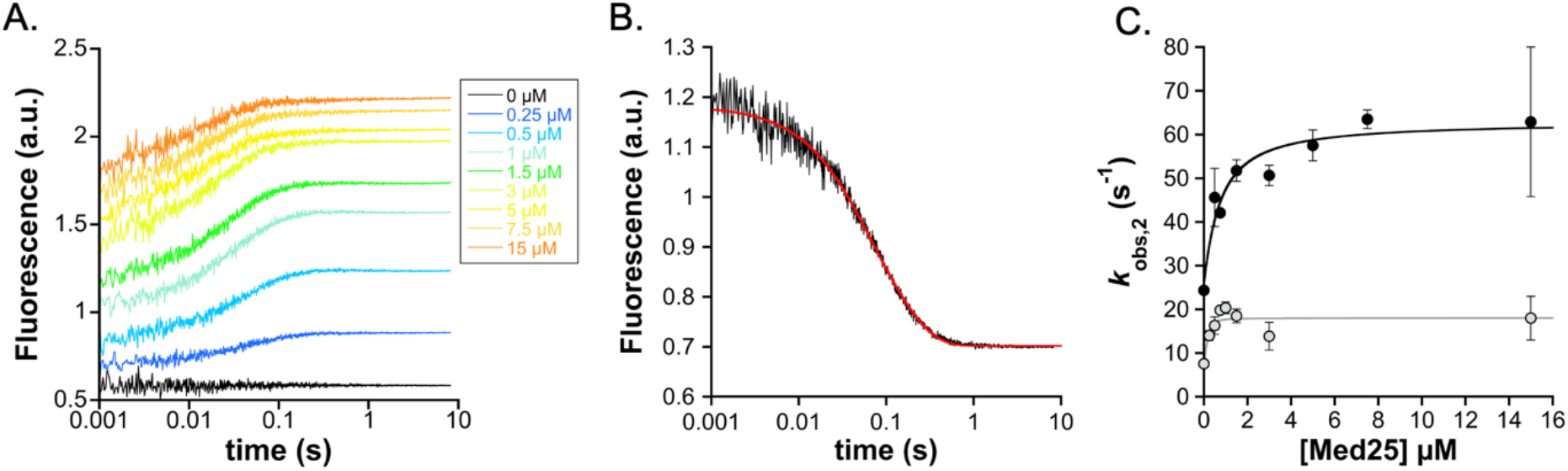
A. Association traces for binding of 0.25 μM 4-DMN-ETV4(45-76)^QF^ to Med25. Concentrations of Med25 are as noted. B. Dissociation trace from 0.25 μM 4-DMN-ETV4(45-76)^QF^ prebound to 0.5 μM Med25, mixed with 50 μM unlabeled ETV4(45-76). C. Observed rate constants for conformational change phases from curve fitting (black=fast phase, grey=slow phase). Values are average of *n*=2 biological replicates, and error represents the standard deviation.

**Figure S9.**
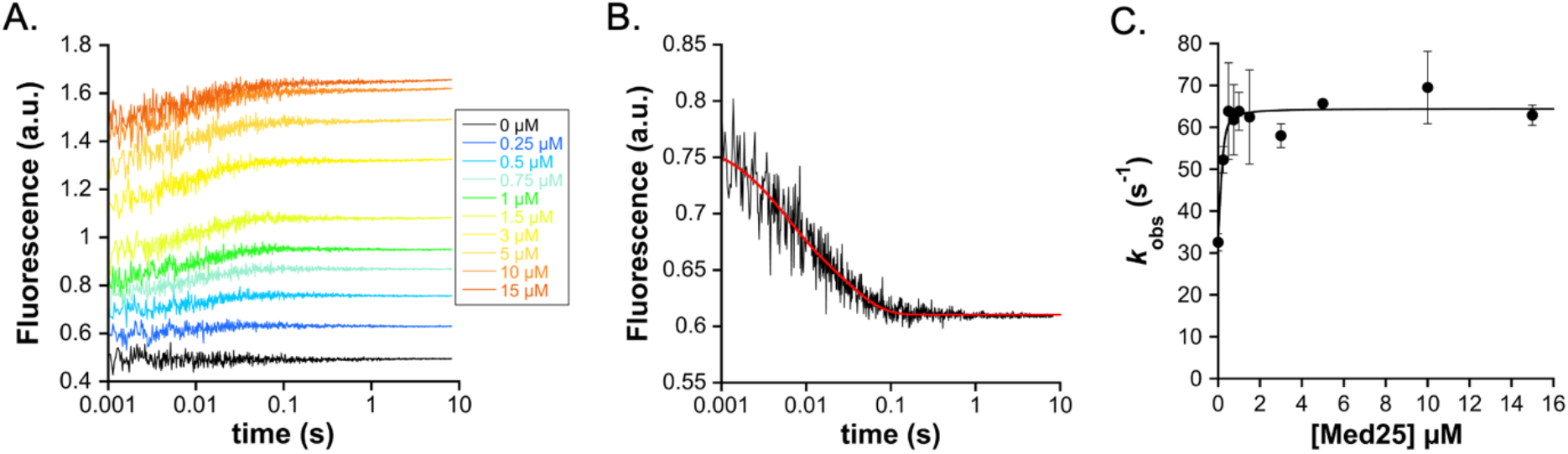
A. Association traces for binding of 0.25 μM 4-DMN-ETV4(45-76)^QL^ to Med25. Concentrations of Med25 are as noted. B. Dissociation trace from 0.25 μM 4-DMN-ETV4(45-76)^QL^ prebound to 0.5 μM Med25, mixed with 50 μM unlabeled ETV4(45-76). C. Observed rate constants for conformational change phases from curve fitting (black=fast phase, grey=slow phase). Values are average of *n*=2 biological replicates, and error represents the standard deviation.

**Figure S10.**
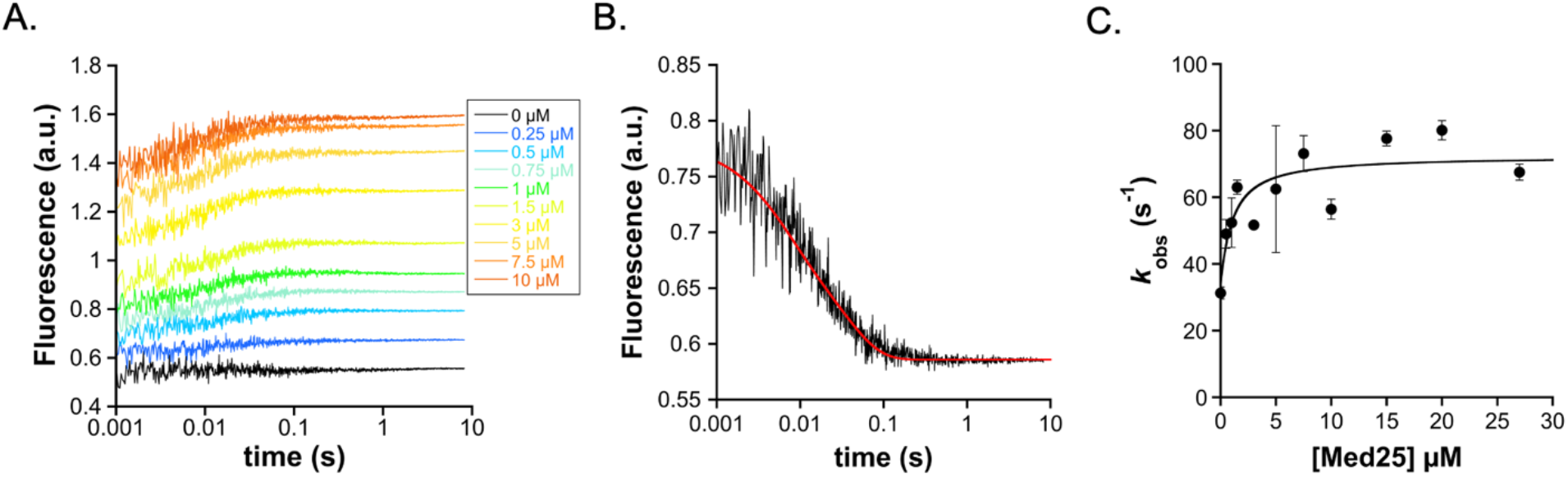
A. Association traces for binding of 0.25 μM 4-DMN-ETV4(45-76)^HL^ to Med25. Concentrations of Med25 are as noted. B. Dissociation trace from 0.25 μM 4-DMN-ETV4(45-76)^HL^ prebound to 0.5 μM Med25, mixed with 50 μM unlabeled ETV4(45-76). C. Observed rate constants for conformational change phases from curve fitting (black=fast phase, grey=slow phase). Values are average of *n*=2 biological replicates, and error represents the standard deviation.

**Figure S11.**
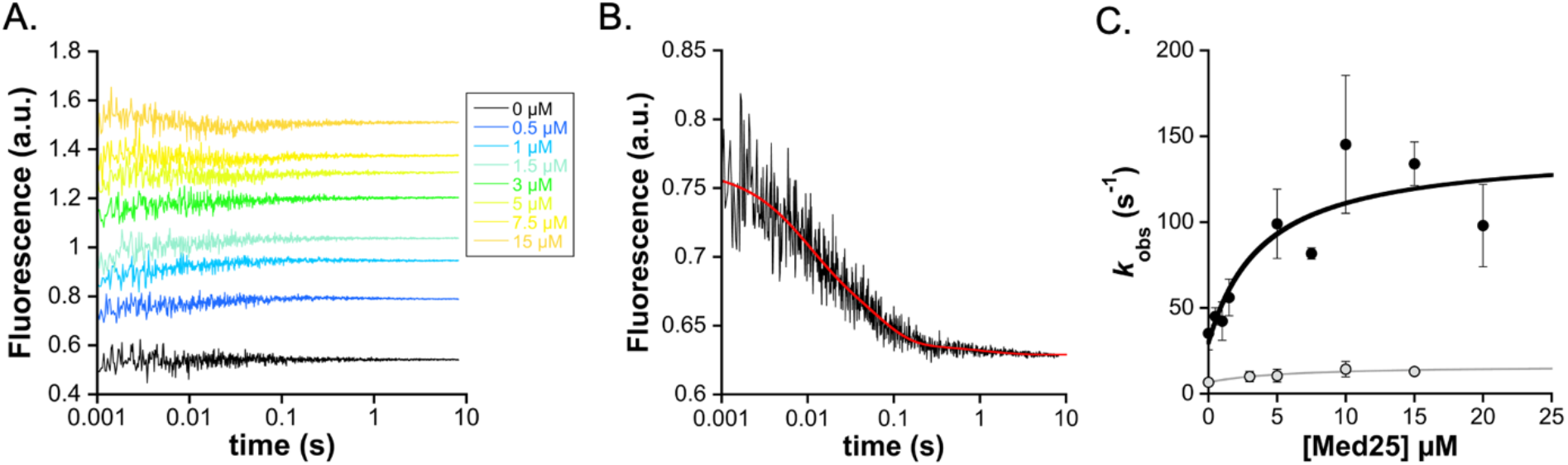
A. Association traces for binding of 0.25 μM 4-DMN-ETV4(45-76)^DLAH/HL^ to Med25. Concentrations of Med25 are as noted. B. Dissociation trace from 0.25 μM 4-DMN-ETV4(45-76)^DLAH/HL^ prebound to 0.5 μM Med25, mixed with 50 μM unlabeled ETV4(45-76). C. Observed rate constants for conformational change phases from curve fitting (black=fast phase, grey=slow phase). Values are average of *n*=2 biological replicates, and error represents the standard deviation.

**Figure S12.**
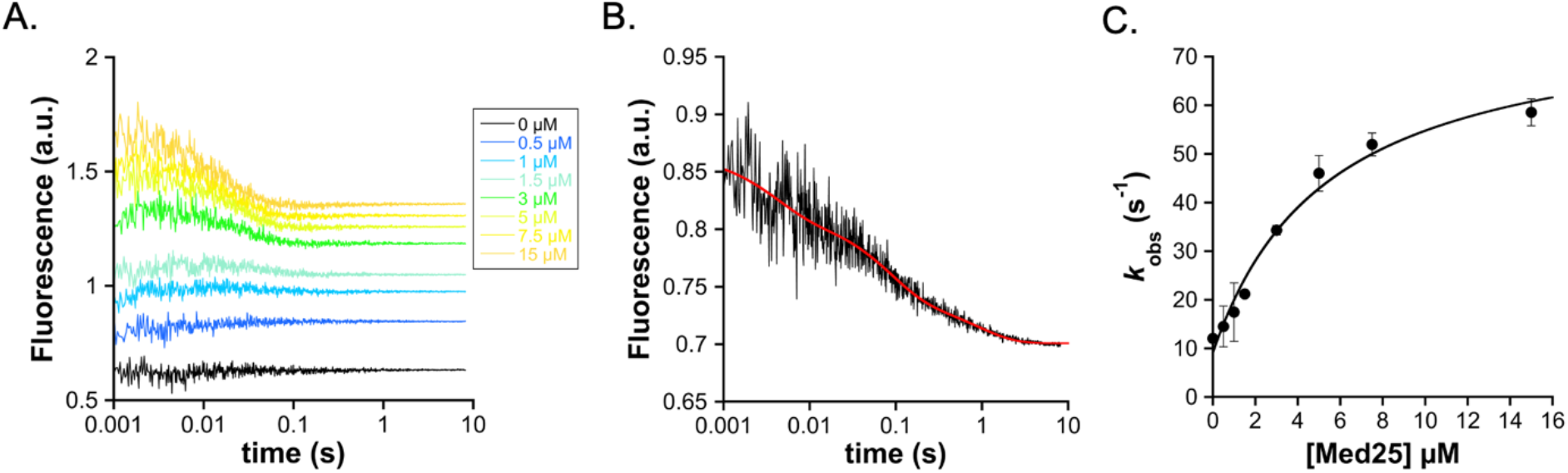
A. Association traces for binding of 0.25 μM 4-DMN-ETV4(45-76)^DLAH/HF^ to Med25. Concentrations of Med25 are as noted. B. Dissociation trace from 0.25 μM 4-DMN-ETV4(45-76)^DLAH/HF^ prebound to 0.5 μM Med25, mixed with 50 μM unlabeled ETV4(45-76). C. Observed rate constants for conformational change phases from curve fitting (black=fast phase, grey=slow phase). Values are average of *n*=2 biological replicates, and error represents the standard deviation.

**Figure S13.**
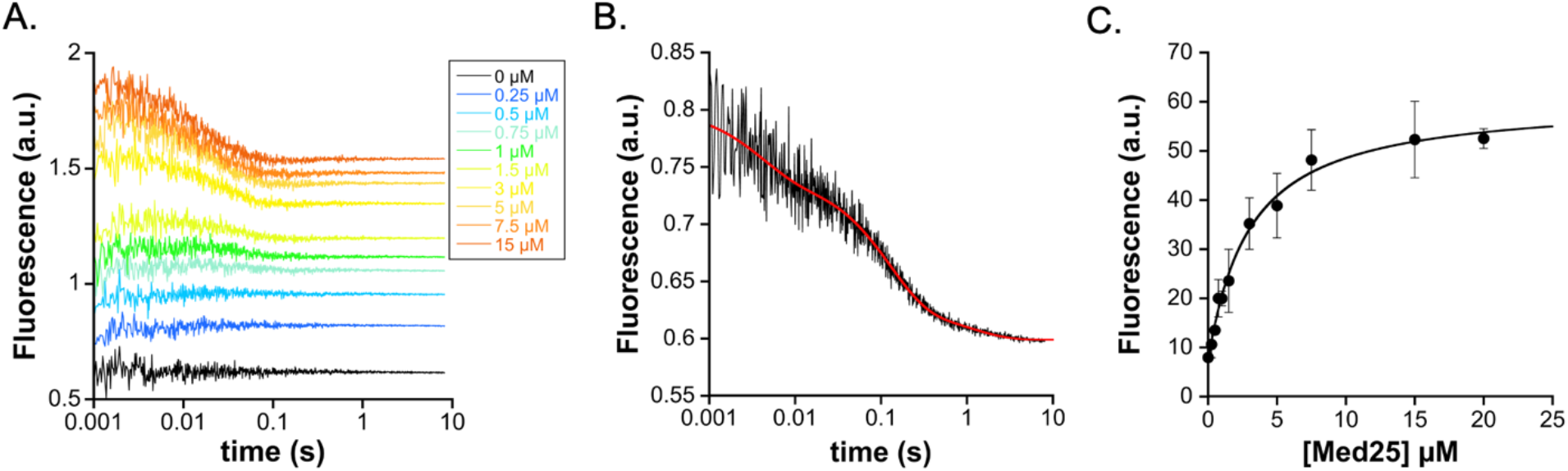
A. Association traces for binding of 0.25 μM 4-DMN-ETV4(45-76)^DLAH/QF^ to Med25. Concentrations of Med25 are as noted. B. Dissociation trace from 0.25 μM 4-DMN-ETV4(45-76)^DLAH/QF^ prebound to 0.5 μM Med25, mixed with 50 μM unlabeled ETV4(45-76). C. Observed rate constants for conformational change phases from curve fitting (black=fast phase, grey=slow phase). Values are average of *n*=2 biological replicates, and error represents the standard deviation.

**Figure S14.**
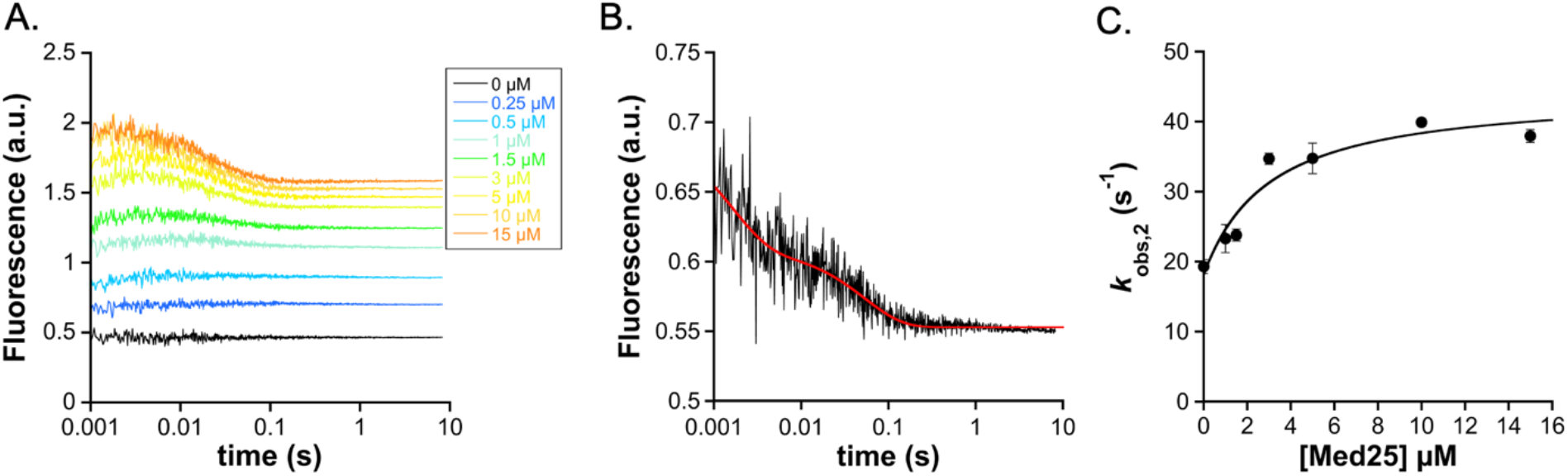
A. Association traces for binding of 0.25 μM 4-DMN-ETV1(42-69) to Med25. Concentrations of Med25 are as noted. B. Dissociation trace from 0.25 μM 4-DMN-ETV1(42-69) prebound to 0.5 μM Med25, mixed with 50 μM unlabeled ETV1(38-69). C. Observed rate constants for conformational change phases from curve fitting (black=fast phase, grey=slow phase). Values are average of *n*=2 biological replicates, and error represents the standard deviation.

**Figure S15.**
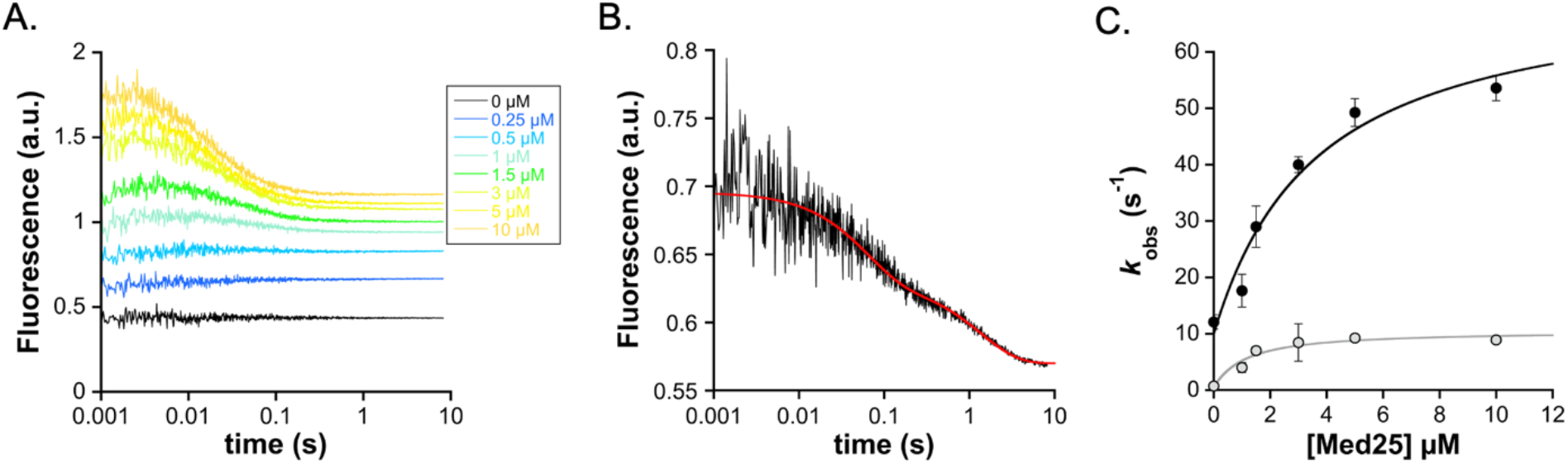
A. Association traces for binding of 0.25 μM 4-DMN-ETV4(49-76) to Med25. Concentrations of Med25 are as noted. B. Dissociation trace from 0.25 μM 4-DMN-ETV4(49-76) prebound to 0.5 μM Med25, mixed with 50 μM unlabeled ETV4(45-76). C. Observed rate constants for conformational change phases from curve fitting (black=fast phase, grey=slow phase). Values are average of *n*=2 biological replicates, and error represents the standard deviation.

### NMR Data

The full NMR datasets used to generate the figures in the manuscript are shown below.

**Figure S16.**
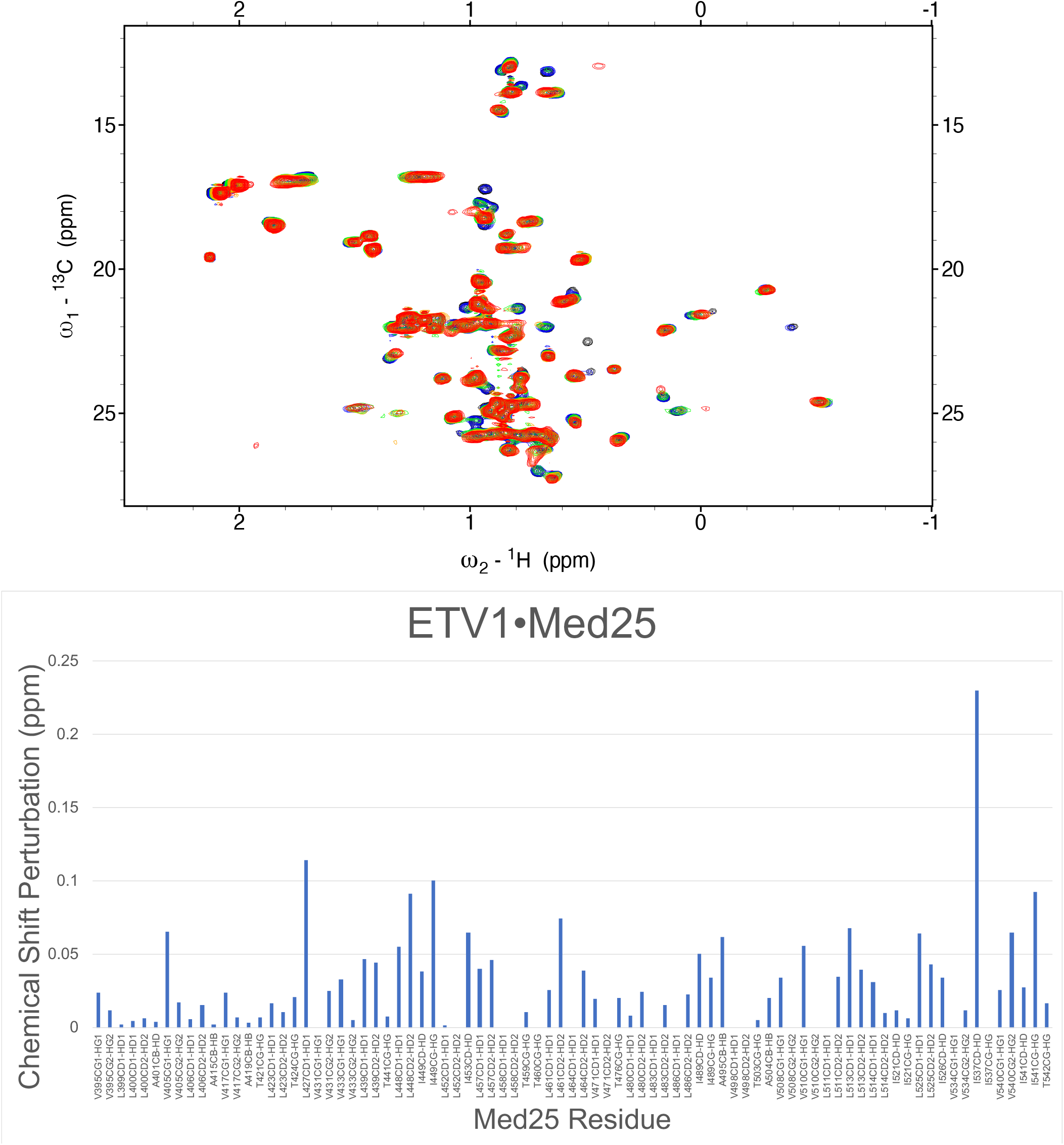
Above: Overlay of ^1^H,^13^C HSQC spectra of Med25 upon titration with unlabeled ETV1. Spectra are free Med25 (50 μM, black), 0.2 eq ETV1 (blue), 0.5 eq ETV1 (green), 0.8 eq ETV1 (orange) 1.1 eq ETV1 (red). Below: CSP mapping of Med25 bound to 1.1 eq ETV1.

**Figure S17.**
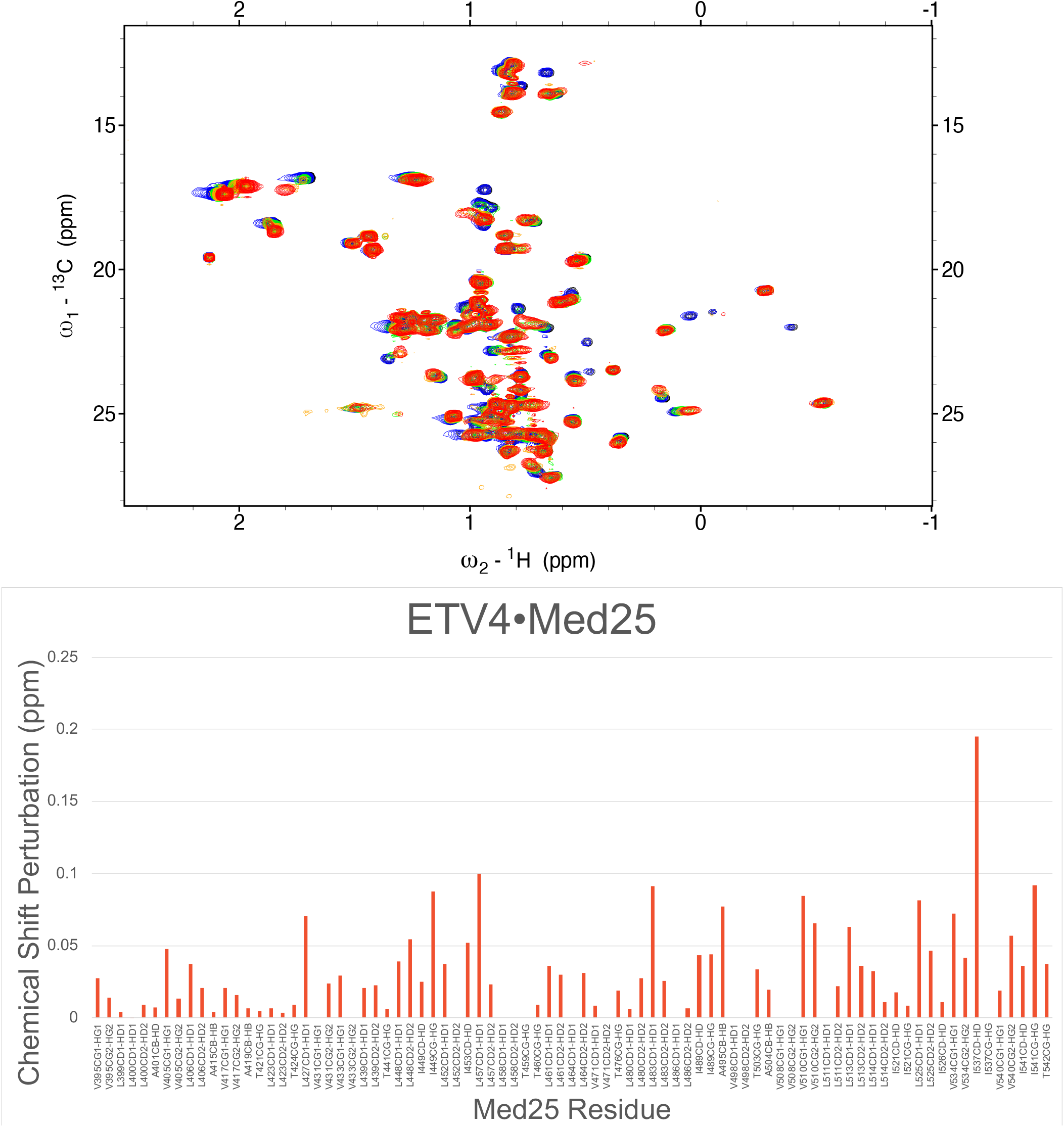
Above: Overlay of ^1^H,^13^C HSQC spectra of Med25 upon titration with unlabeled ETV4. Spectra are free Med25 (50 μM, black), 0.2 eq ETV4 (blue), 0.5 eq ETV4 (green), 0.8 eq ETV4 (orange) 1.1 eq ETV4 (red). Below: CSP mapping of Med25 bound to 1.1 eq ETV4.

**Figure S18.**
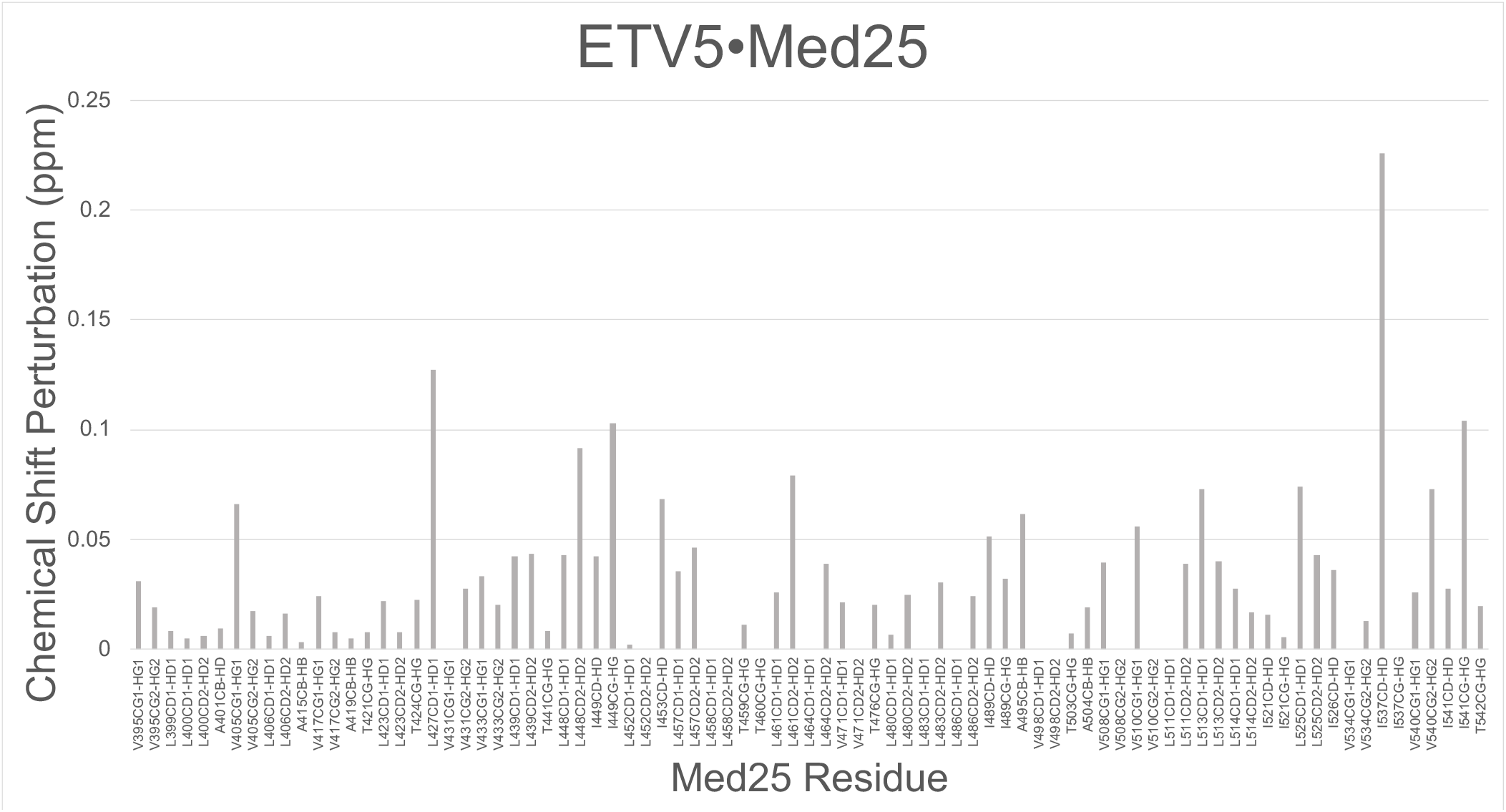
CSP mapping of Med25 bound to 1.1 eq ETV5. Assignments were generated by comparison to the essentially identical spectra of Med25 bound to ETV1.

**Figure S19.**
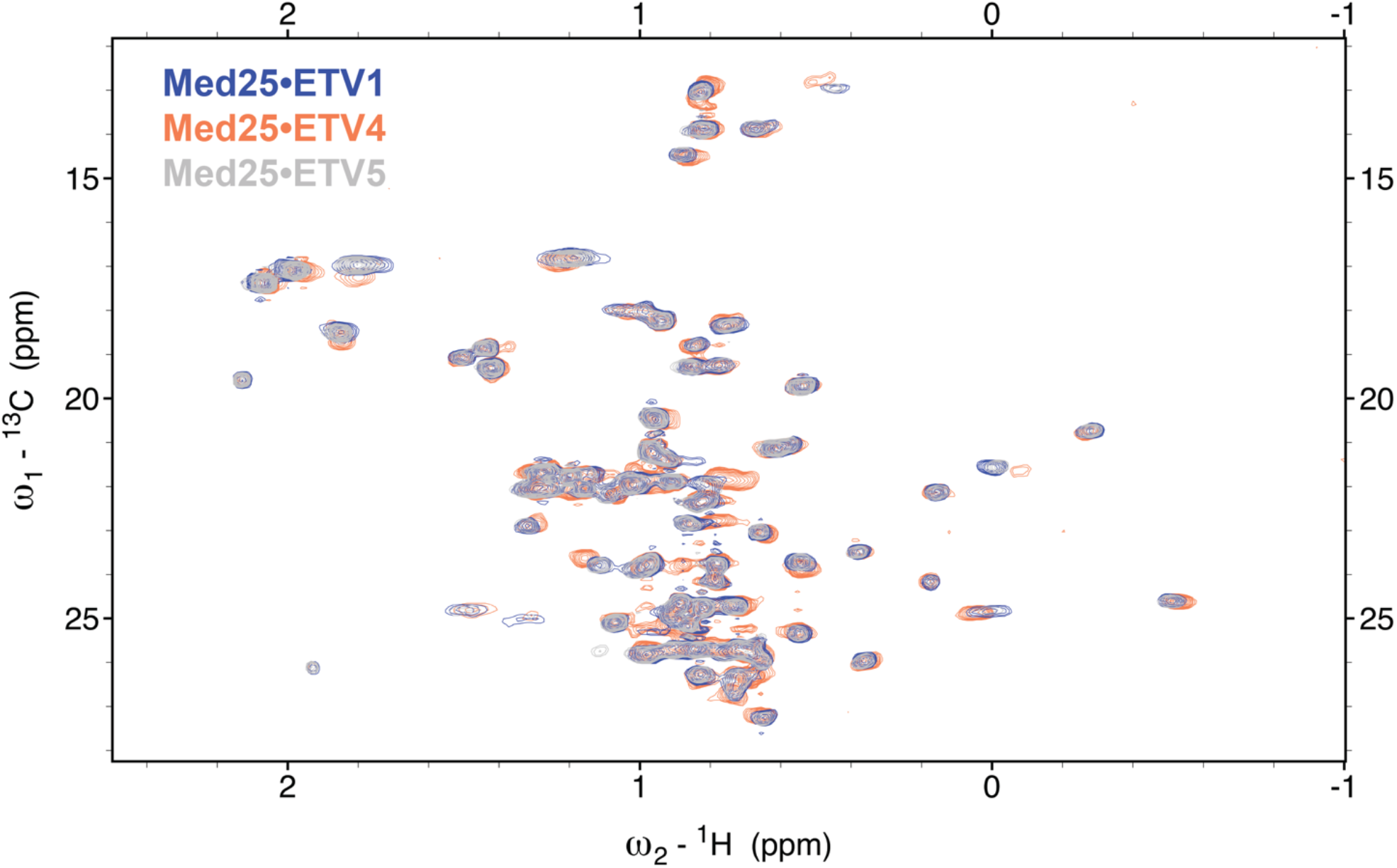
Overlay of ^1^H,^13^C HSQC spectra of Med25 bound to ETV1 (blue), ETV4 (orange), and ETV5 (grey).

**Figure S20.**
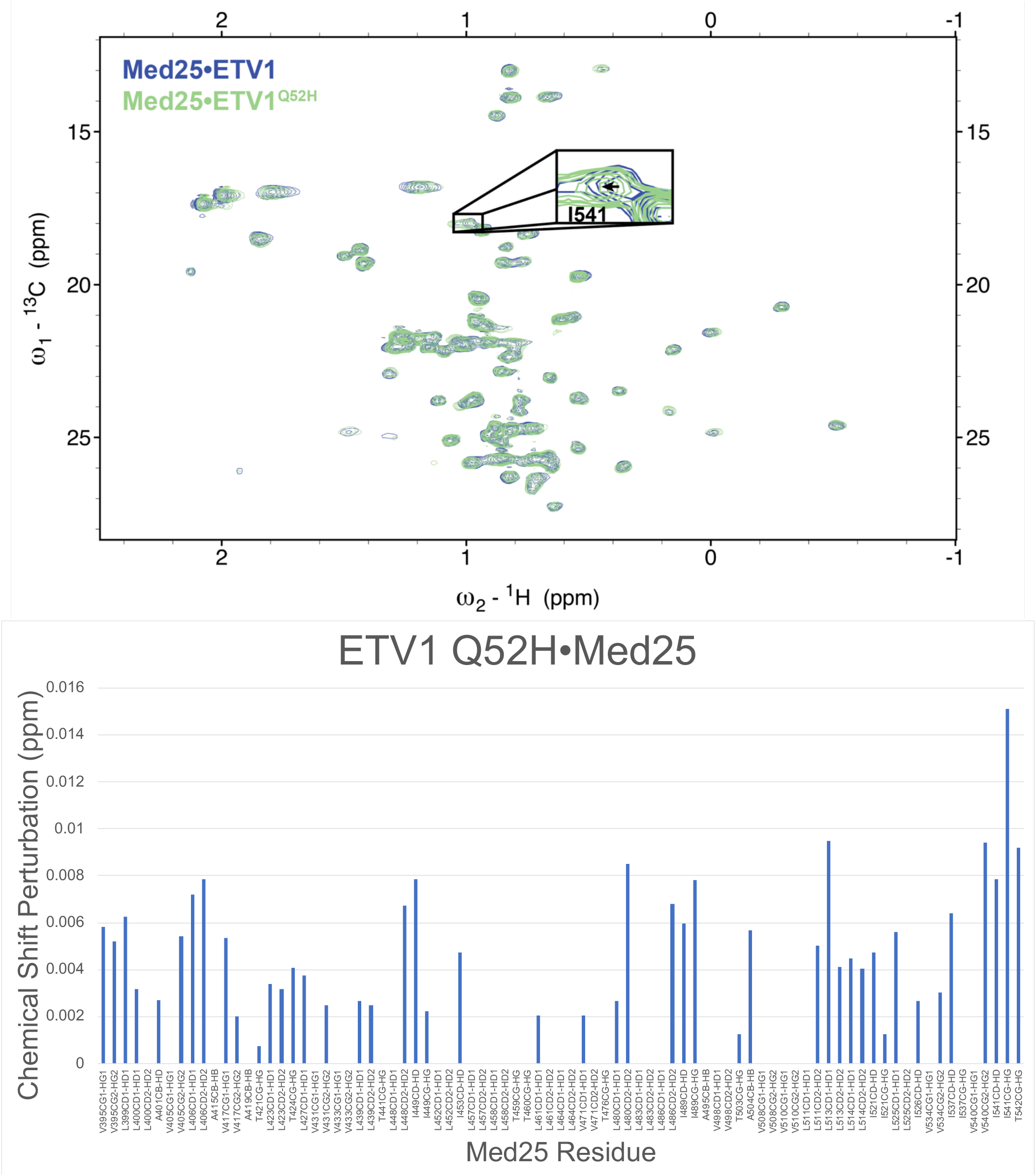
Above: Overlay of ^1^H,^13^C HSQC spectra of Med25 bound to ETV1 helical region soft mutation variant. Spectra are Med25 (60 μM) bound to 1.1 eq ETV1 (blue) or ETV1^Q52H^ (green). Below: CSP mapping of differences between ETV1- and ETV1^Q52H^-bound Med25 spectra.

**Figure S21.**
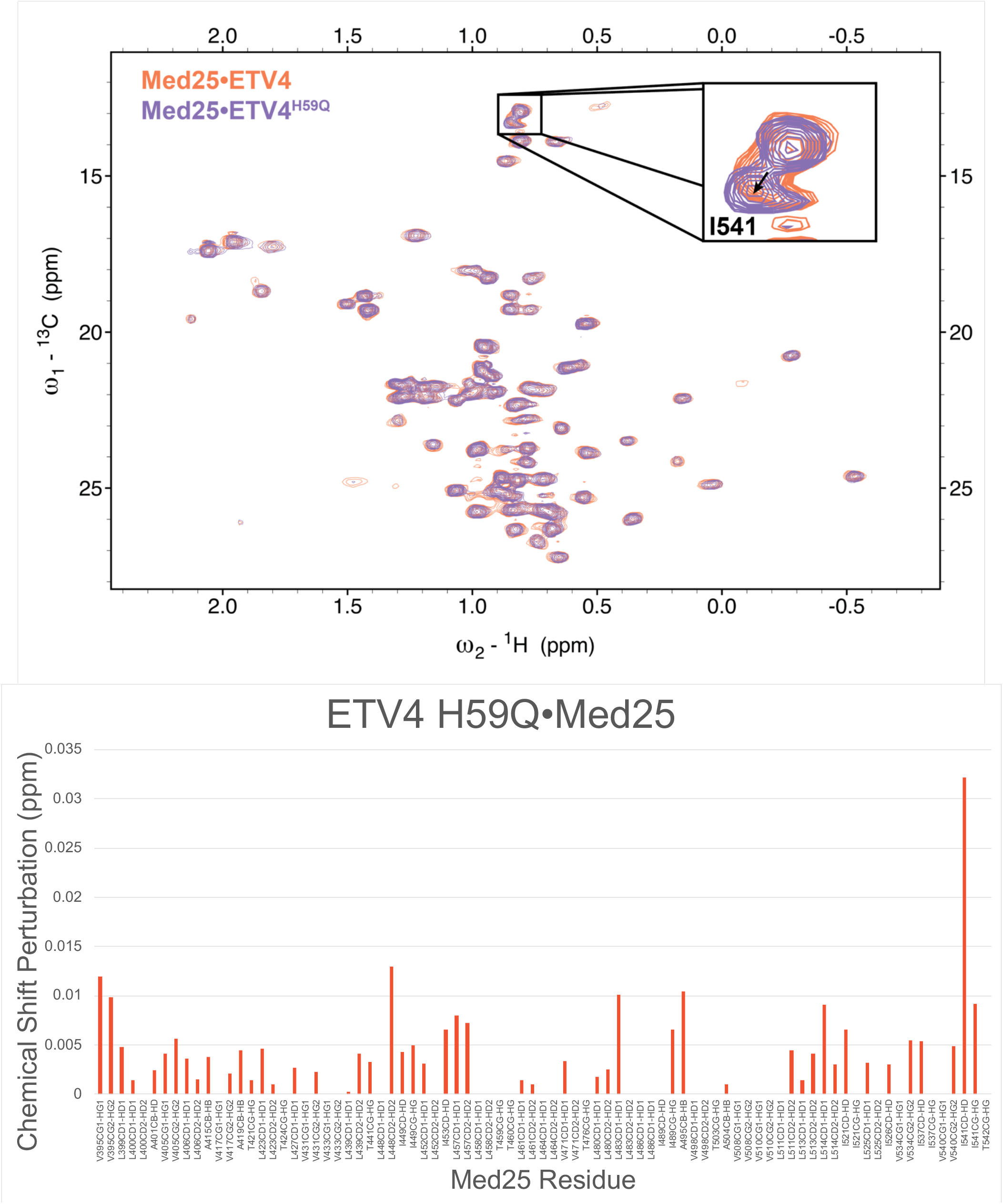
Above: Overlay of ^1^H,^13^C HSQC spectra of Med25 bound to ETV4 helical region soft mutation variant. Spectra are Med25 (60 μM) bound to 1.1 eq ETV4 (orange) or ETV4^H59Q^ (light purple). Below: CSP mapping of differences between ETV4- and ETV4^H59Q^-bound Med25 spectra.

**Figure S22.**
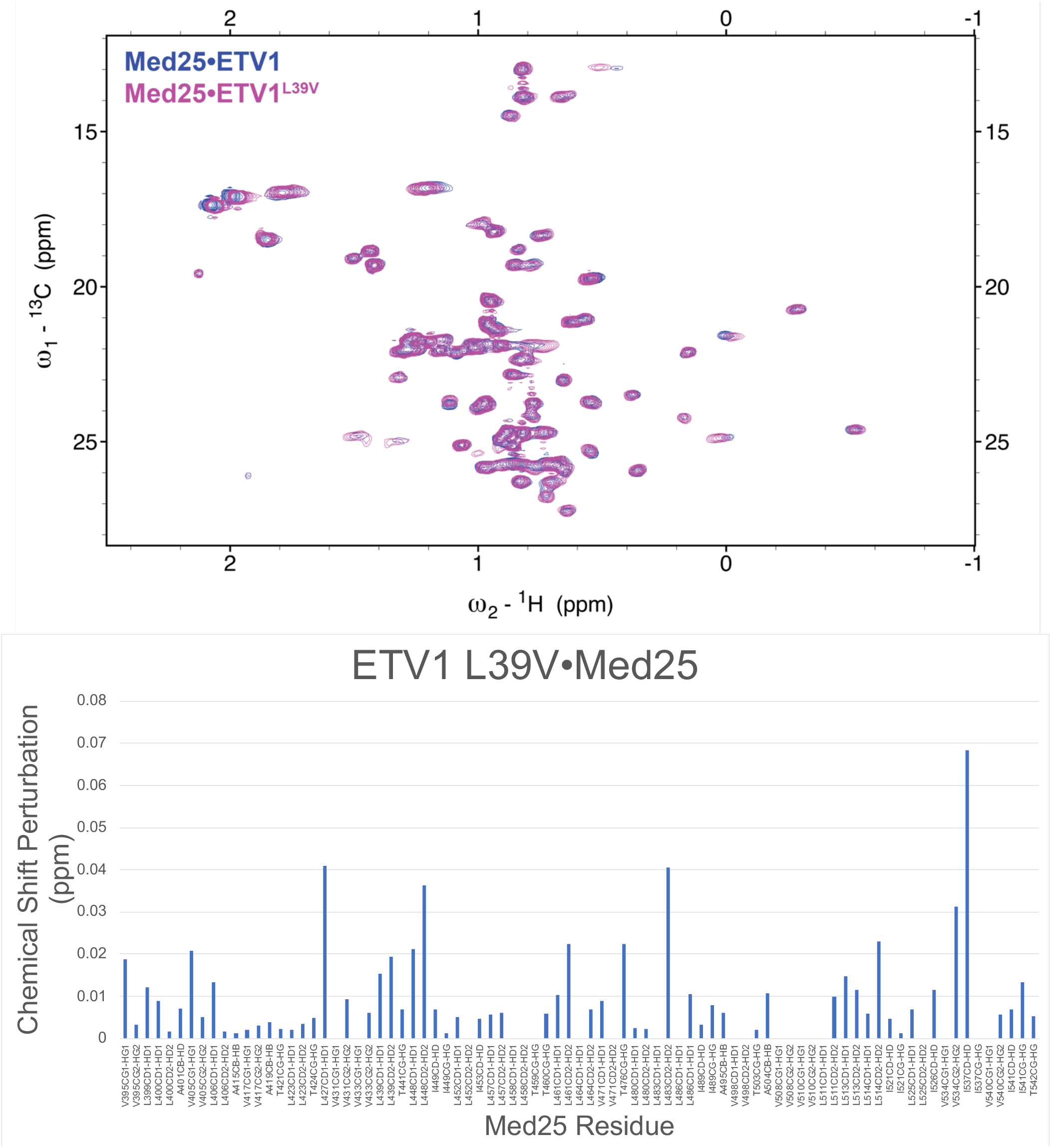
Above: Overlay of ^1^H,^13^C HSQC spectra of Med25 bound to ETV1 *N*-terminal soft mutation variant. Spectra are Med25 (70 μM) bound to 1.1 eq ETV1 (blue) or ETV1^L39V^ (purple). Below: CSP mapping of differences between ETV1- and ETV1^L39V^-bound Med25 spectra.

**Figure S23.**
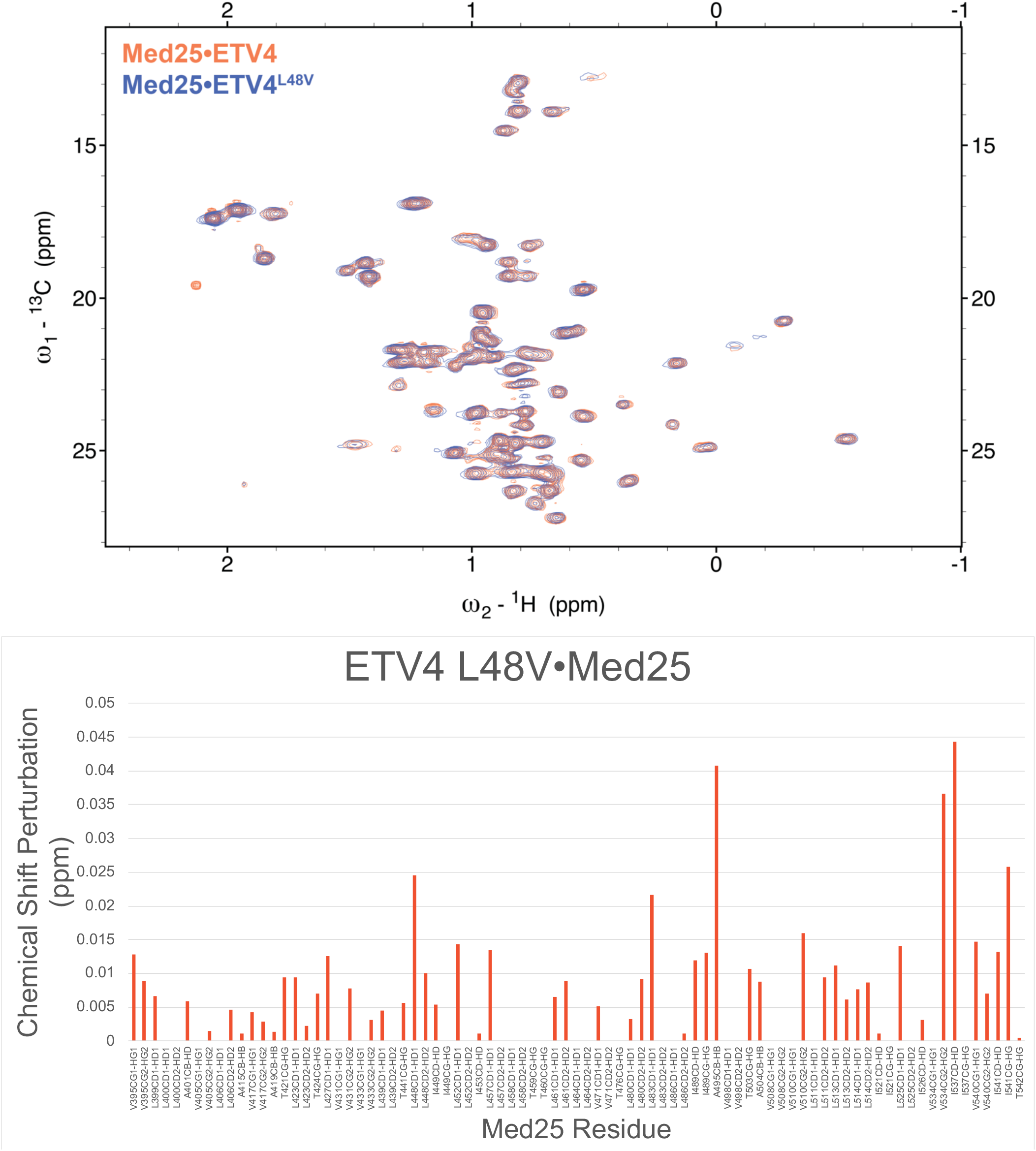
Above: Overlay of ^1^H,^13^C HSQC spectra of Med25 bound to ETV4 *N*-terminal soft mutation variant. Spectra are Med25 (70 μM) bound to 1.1 eq ETV4 (orange) or ETV4^L48V^ (blue). Below: CSP mapping of differences between ETV4- and ETV4^L48V^-bound Med25 spectra.

**Figure S24.**
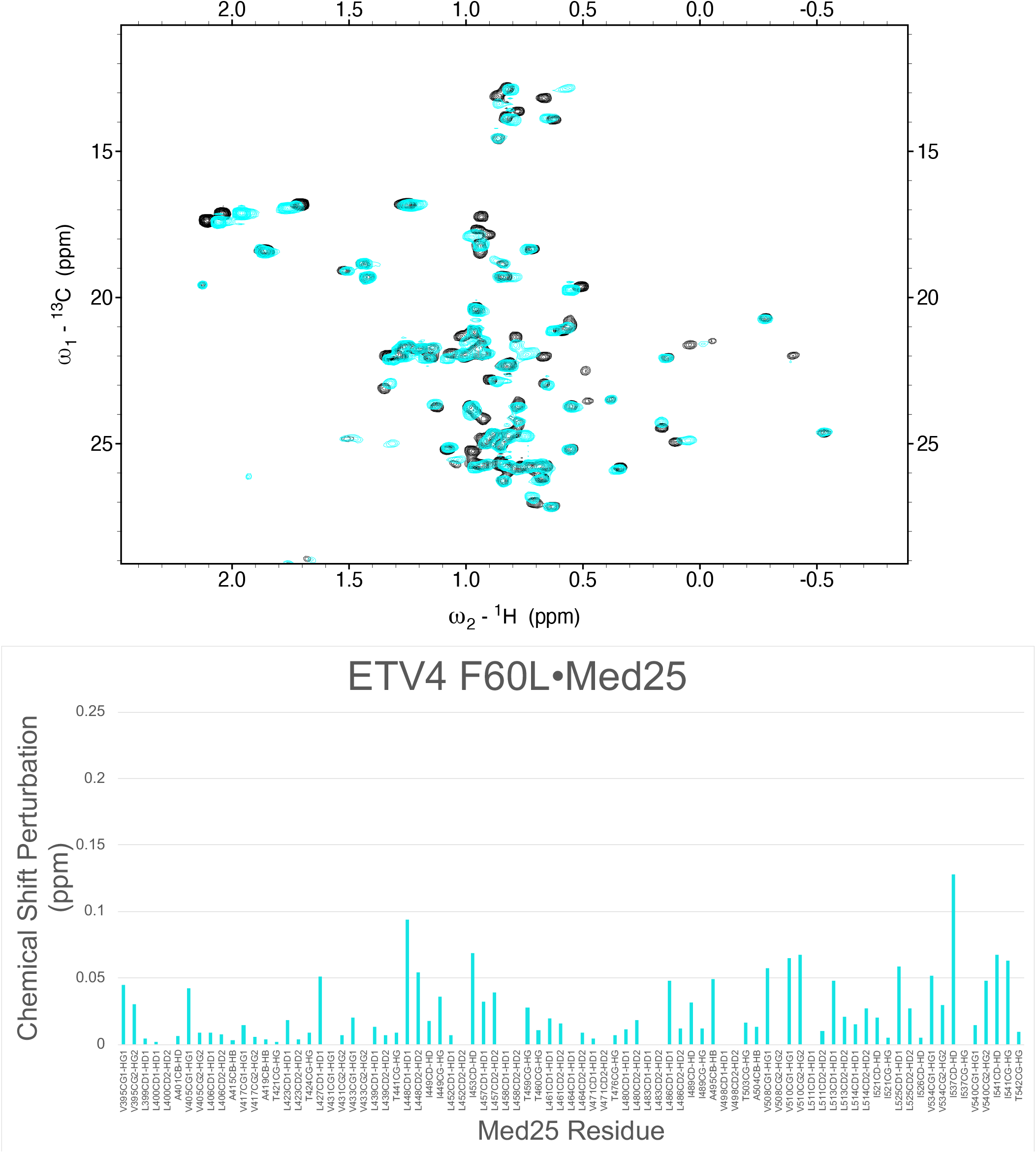
Above: Overlay of ^1^H,^13^C HSQC spectra of Med25 bound to ETV4 helical region variant that redistributes conformational ensemble. Spectra are Med25 (60 μM, black) and Med25 bound to 1.1 eq ETV4^F60L^ (cyan). Below: CSP mapping of Med25 bound to ETV4^F60L^.

**Figure S25.**
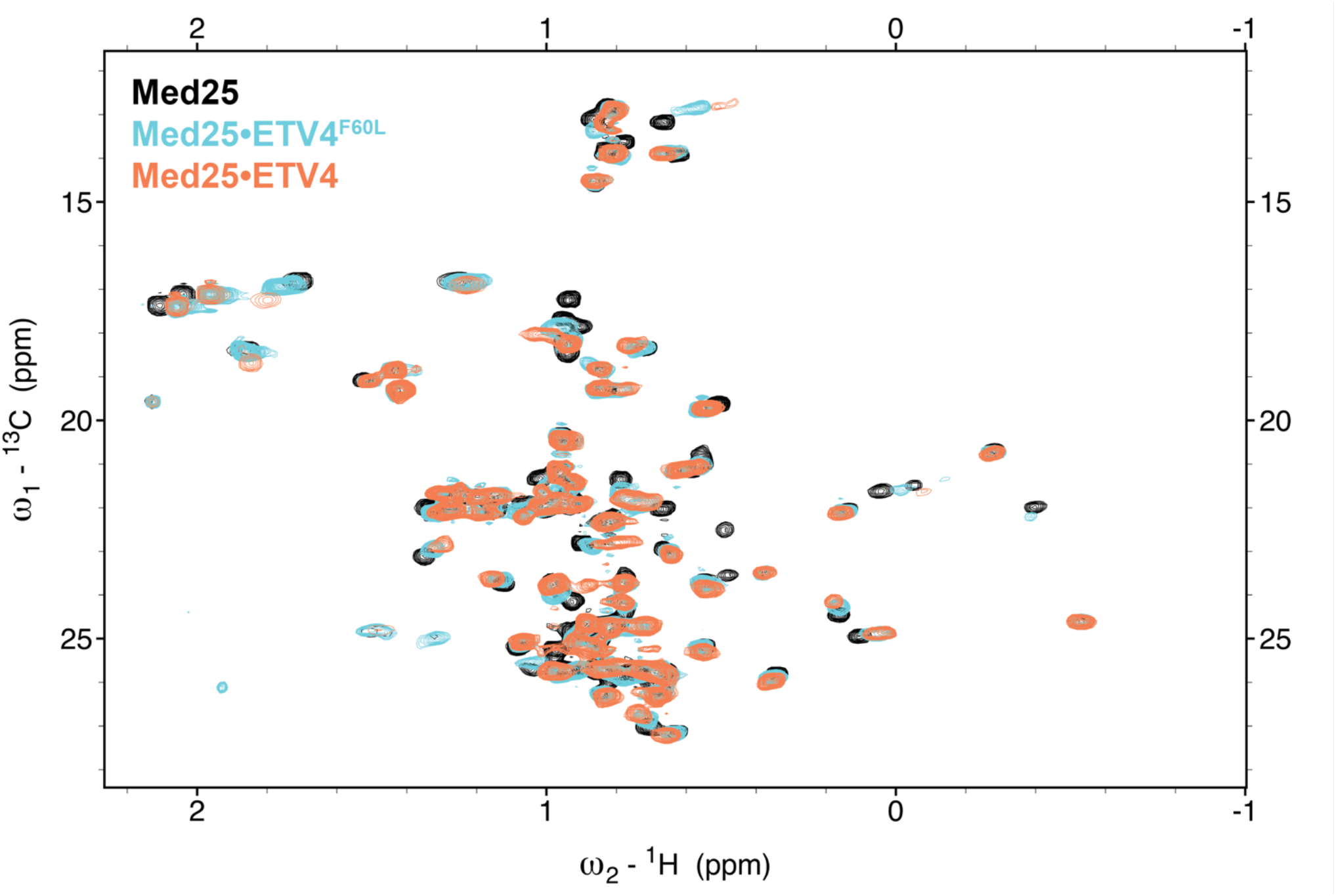
Overlay of ^1^H,^13^C HSQC spectra of free Med25 (60 μM, black), 1.1 eq ETV4^F60L^ (cyan), 1.1 eq ETV4 (orange).

### Peptide Characterization

The characterization data for all peptides in this manuscript is shown below.

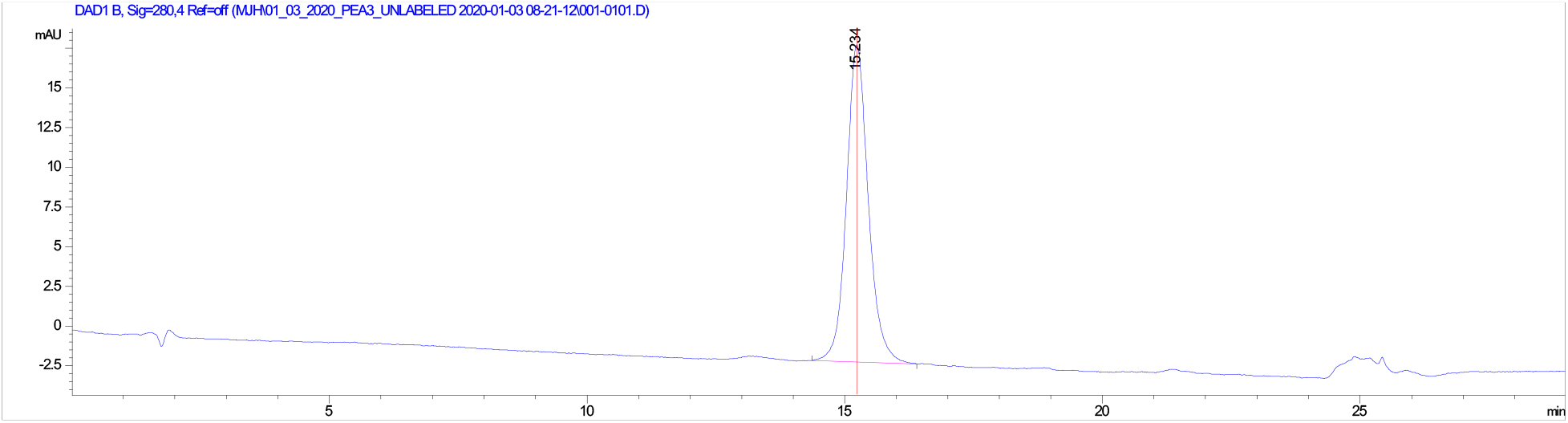

Analytical HPLC trace of **ETV1(38-69)**, monitored at 280 nm. Analytical sample was run in a water (with 100 mM ammonium acetate)/ acetonitrile system. The sample was injected with an isocratic flow of 70% water (with 100 mM ammonium acetate) and 30% acetonitrile. After 2 mins, the solvent gradient was increased from 10-35% acetonitrile over 20 mins.

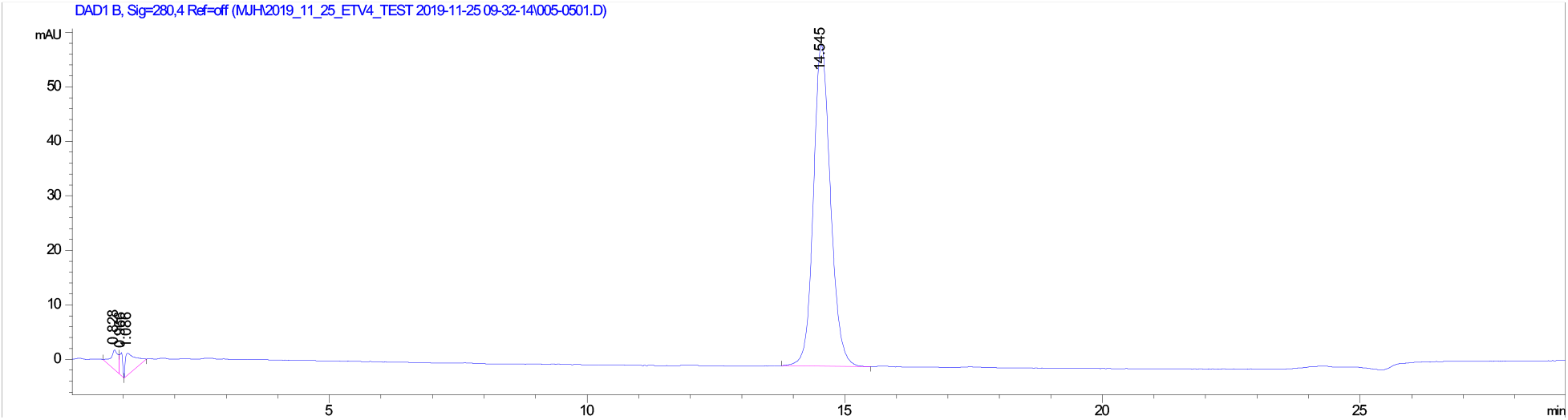

Analytical HPLC trace of **ETV4(45-76)**, monitored at 280 nm. Analytical sample was run in a water (with 100 mM ammonium acetate)/ acetonitrile system. The sample was injected with an isocratic flow of 70% water (with 100 mM ammonium acetate) and 30% acetonitrile. After 2 mins, the solvent gradient was increased from 10-35% acetonitrile over 20 mins.

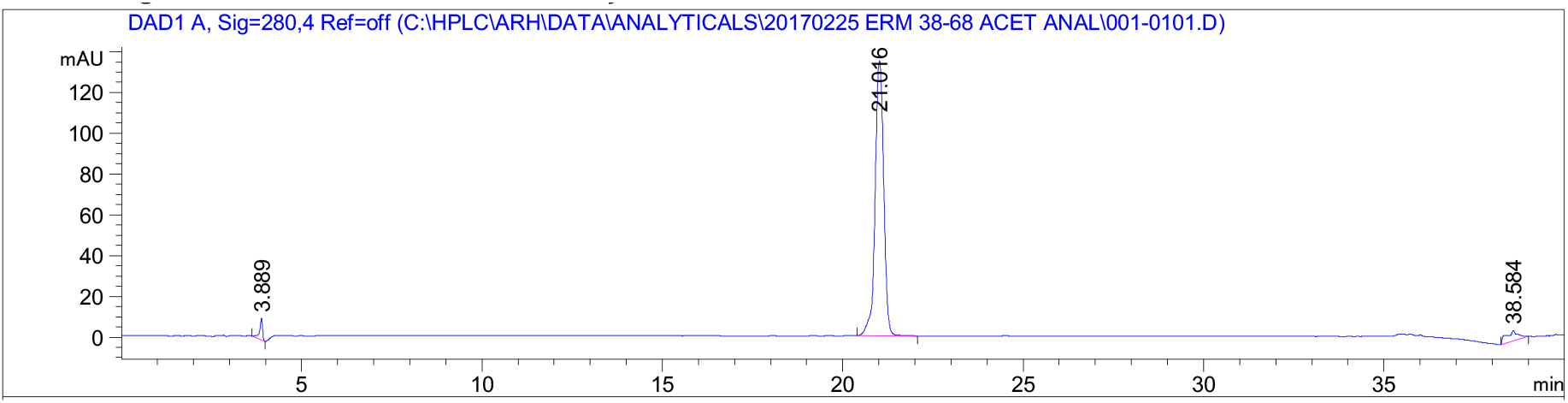

Analytical HPLC trace of **ETV5(38-68)**, monitored at 280 nm. Analytical sample was run in a water (with 100 mM ammonium acetate)/acetonitrile system. The sample was injected with an isocratic flow of 85% water (with 100 mM ammonium acetate) and 15% acetonitriile. After 2 mins, the solvent gradient was increased from 15-30% acetonitrile over 20 mins.

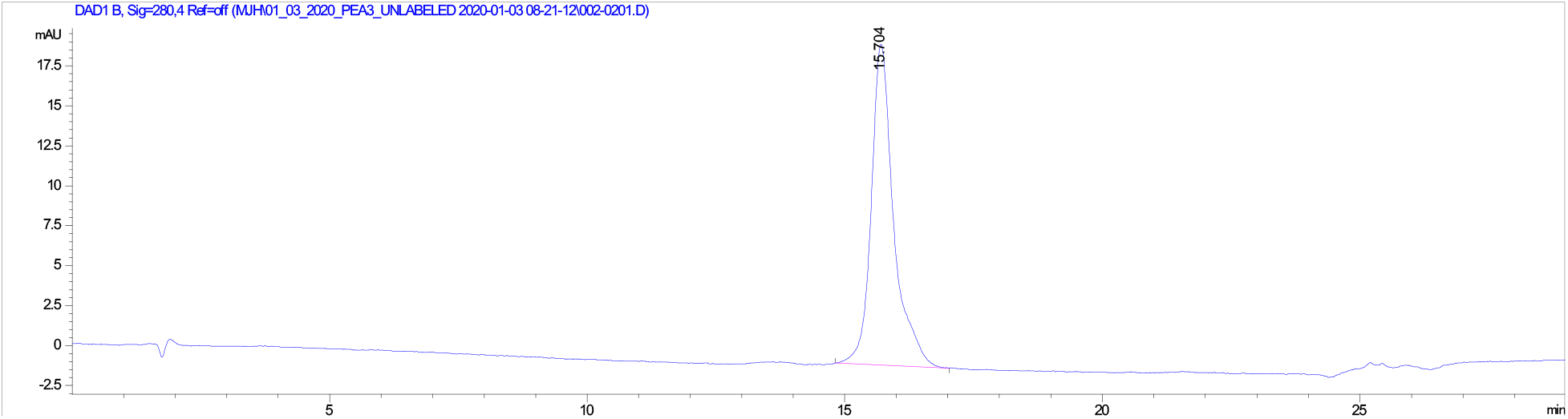

Analytical HPLC trace of **ETV1(38-69)^Q52H^**, monitored at 280 nm. Analytical sample was run in a water (with 100 mM ammonium acetate)/ acetonitrile system. The sample was injected with an isocratic flow of 70% water (with 100 mM ammonium acetate) and 30% acetonitrile. After 2 mins, the solvent gradient was increased from 10-35% acetonitrile over 20 mins.

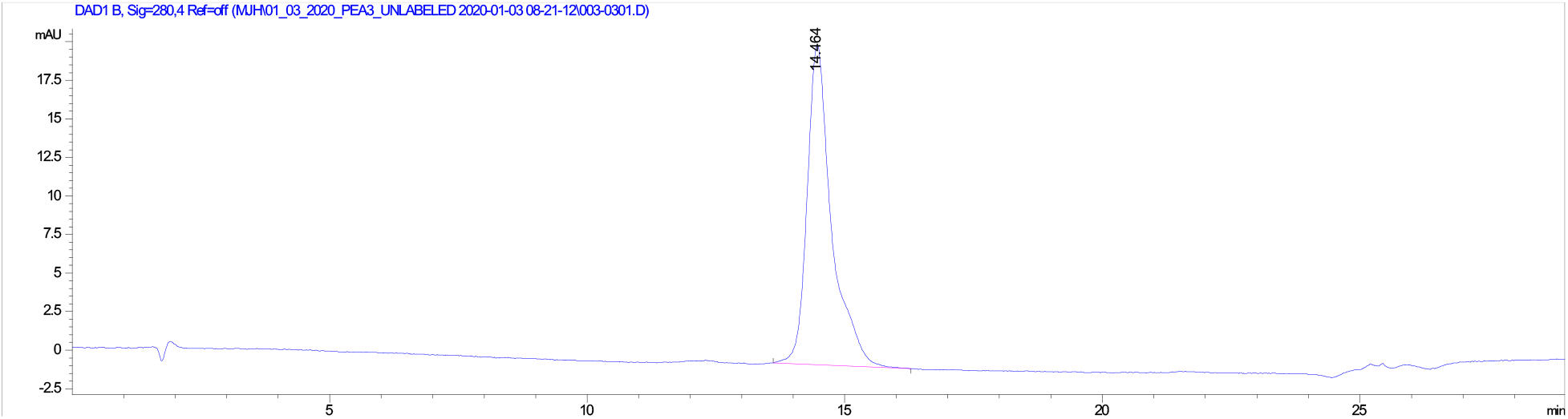

Analytical HPLC trace of **ETV1(38-69)^L39V^**, monitored at 280 nm. Analytical sample was run in a water (with 100 mM ammonium acetate)/ acetonitrile system. The sample was injected with an isocratic flow of 70% water (with 100 mM ammonium acetate) and 30% acetonitrile. After 2 mins, the solvent gradient was increased from 10-35% acetonitrile over 20 mins.

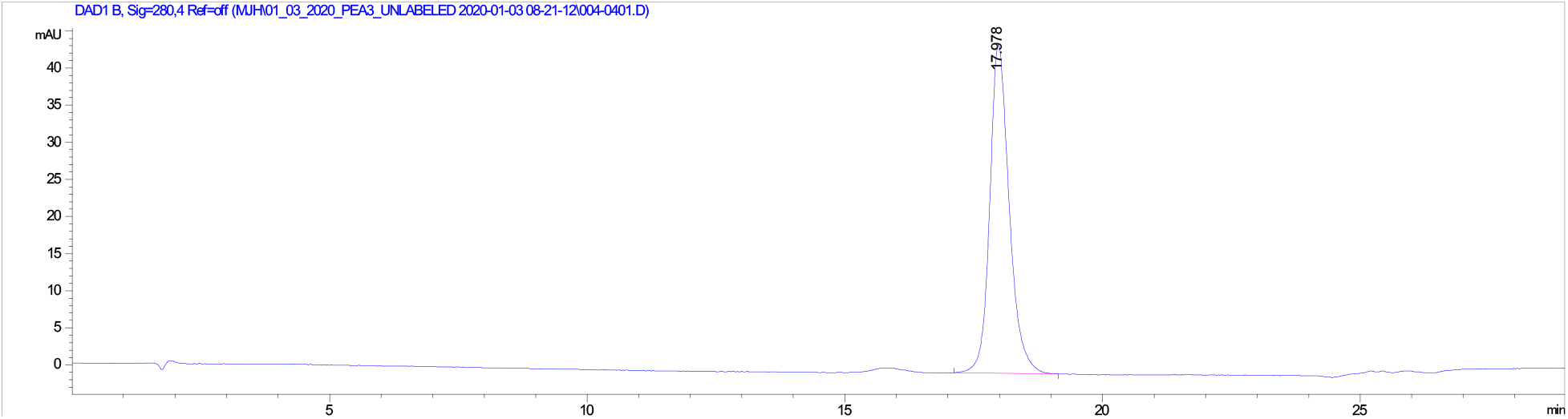

Analytical HPLC trace of **ETV4(45-76)^F60L^**, monitored at 280 nm. Analytical sample was run in a water (with 100 mM ammonium acetate)/ acetonitrile system. The sample was injected with an isocratic flow of 70% water (with 100 mM ammonium acetate) and 30% acetonitrile. After 2 mins, the solvent gradient was increased from 10-35% acetonitrile over 20 mins.

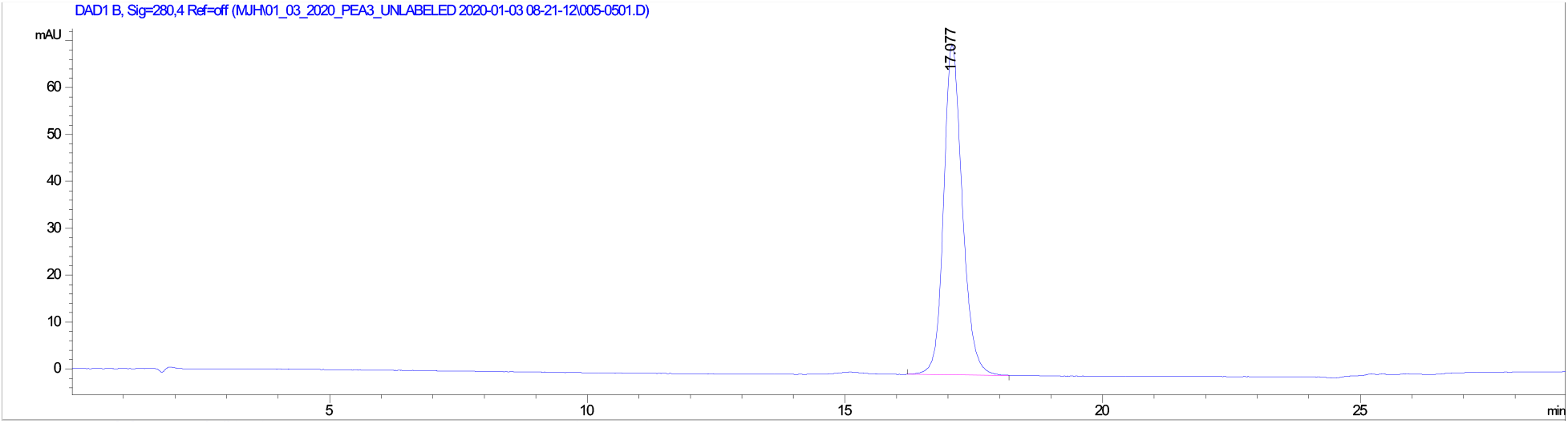

Analytical HPLC trace of **ETV4(45-76)^H59Q^**, monitored at 280 nm. Analytical sample was run in a water (with 100 mM ammonium acetate)/ acetonitrile system. The sample was injected with an isocratic flow of 70% water (with 100 mM ammonium acetate) and 30% acetonitrile. After 2 mins, the solvent gradient was increased from 10-35% acetonitrile over 20 mins.

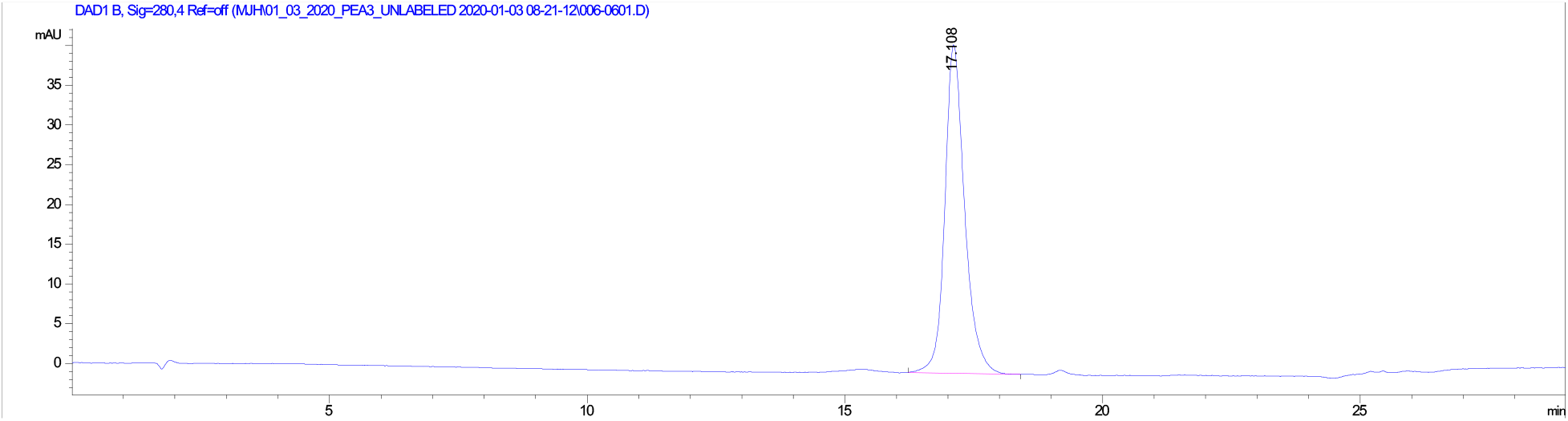

Analytical HPLC trace of **ETV4(45-76)^L48V^**, monitored at 280 nm. Analytical sample was run in a water (with 100 mM ammonium acetate)/ acetonitrile system. The sample was injected with an isocratic flow of 70% water (with 100 mM ammonium acetate) and 30% acetonitrile. After 2 mins, the solvent gradient was increased from 10-35% acetonitrile over 20 mins.

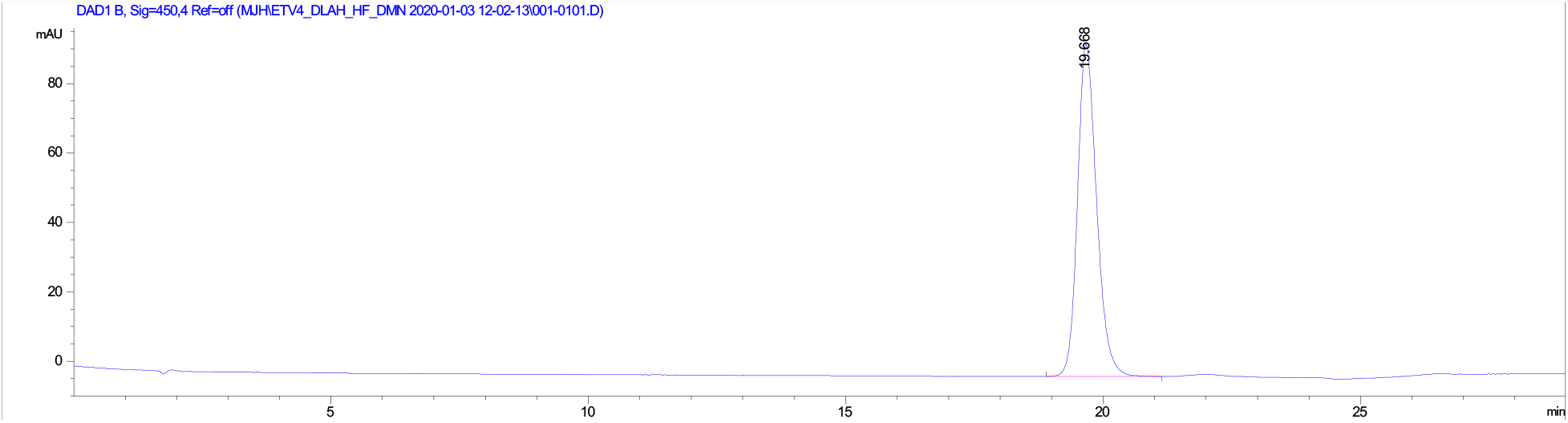

Analytical HPLC trace of **4-DMN-ETV1(38-69)**, monitored at 450 nm. Analytical sample was run in a water (with 100 mM ammonium acetate)/ acetonitrile system. The sample was injected with an isocratic flow of 70% water (with 100 mM ammonium acetate) and 30% acetonitrile. After 2 mins, the solvent gradient was increased from 10-35% acetonitrile over 20 mins.

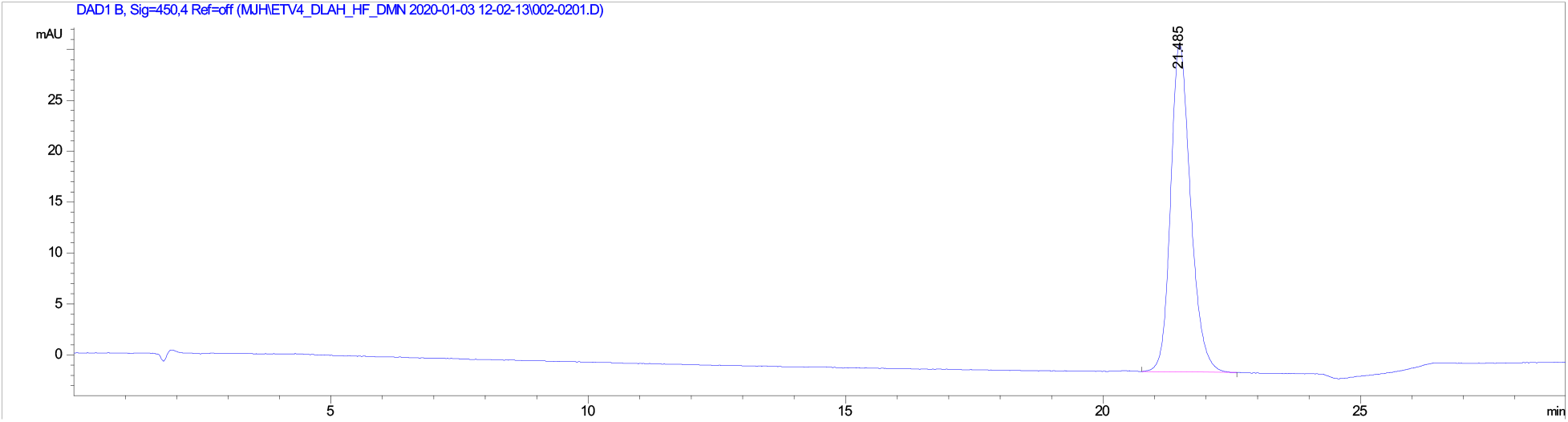

Analytical HPLC trace of **4-DMN-ETV4(45-76)**, monitored at 450 nm. Analytical sample was run in a water (with 100 mM ammonium acetate)/ acetonitrile system. The sample was injected with an isocratic flow of 70% water (with 100 mM ammonium acetate) and 30% acetonitrile. After 2 mins, the solvent gradient was increased from 10-35% acetonitrile over 20 mins.

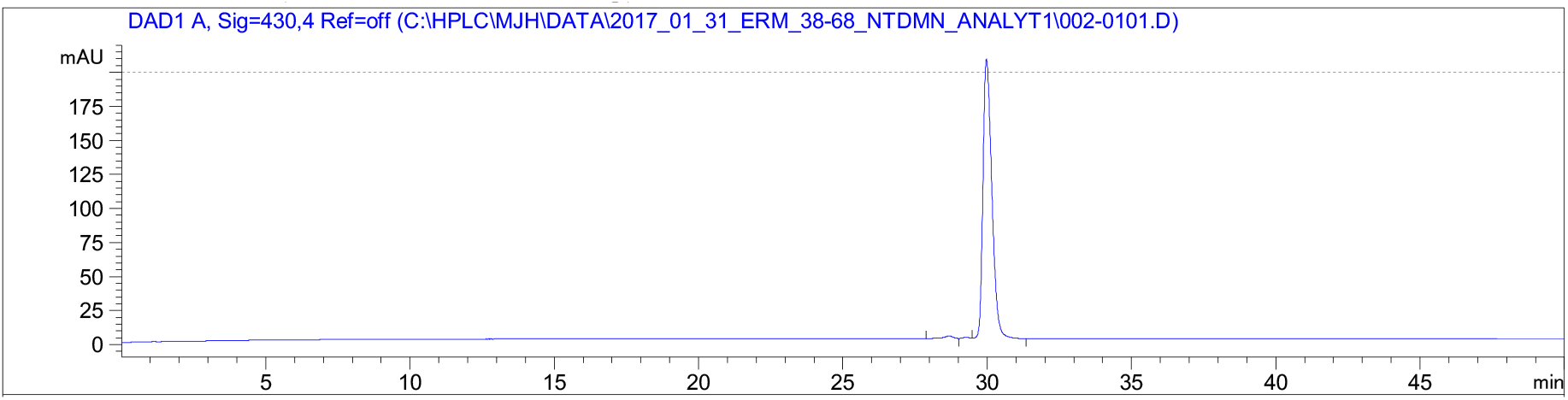

Analytical HPLC trace of **4-DMN-ETV5(38-68)**, monitored at 430 nm. Analytical sample was run in a water (with 100 mM ammonium acetate)/ acetonitrile system. The sample was injected with an isocratic flow of 90% water (with 100 mM ammonium acetate) and 10% acetonitrile. After 2 mins, the solvent gradient was increased from 10-40% acetonitrile over 40 mins.

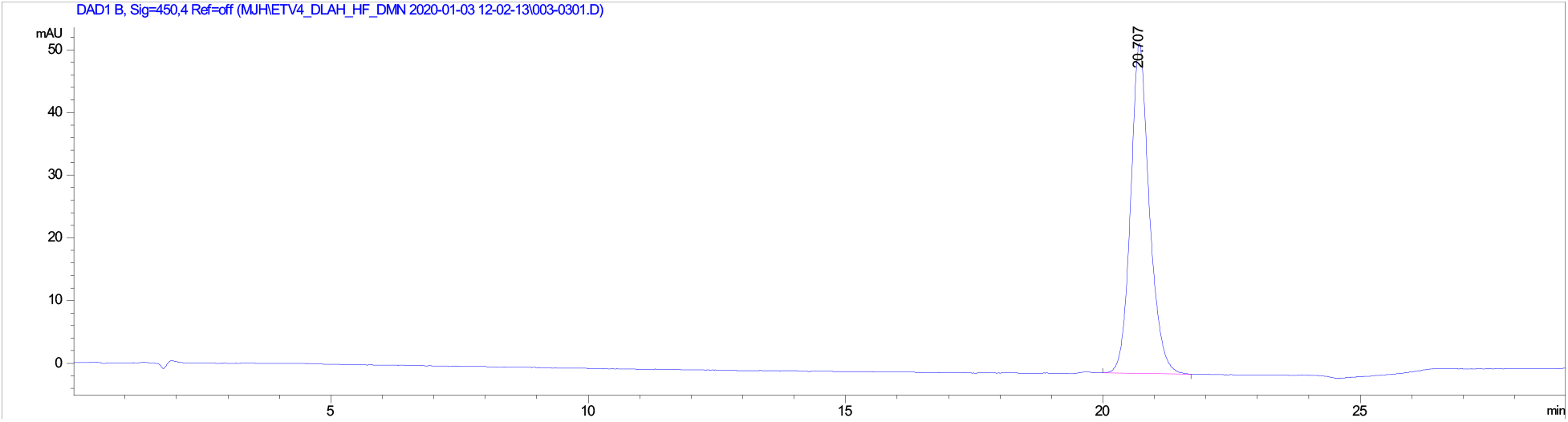

Analytical HPLC trace of **4-DMN-ETV4(45-76)^LPPL/QF^**, monitored at 450 nm. Analytical sample was run in a water (with 100 mM ammonium acetate)/ acetonitrile system. The sample was injected with an isocratic flow of 70% water (with 100 mM ammonium acetate) and 30% acetonitrile. After 2 mins, the solvent gradient was increased from 10-35% acetonitrile over 20 mins.

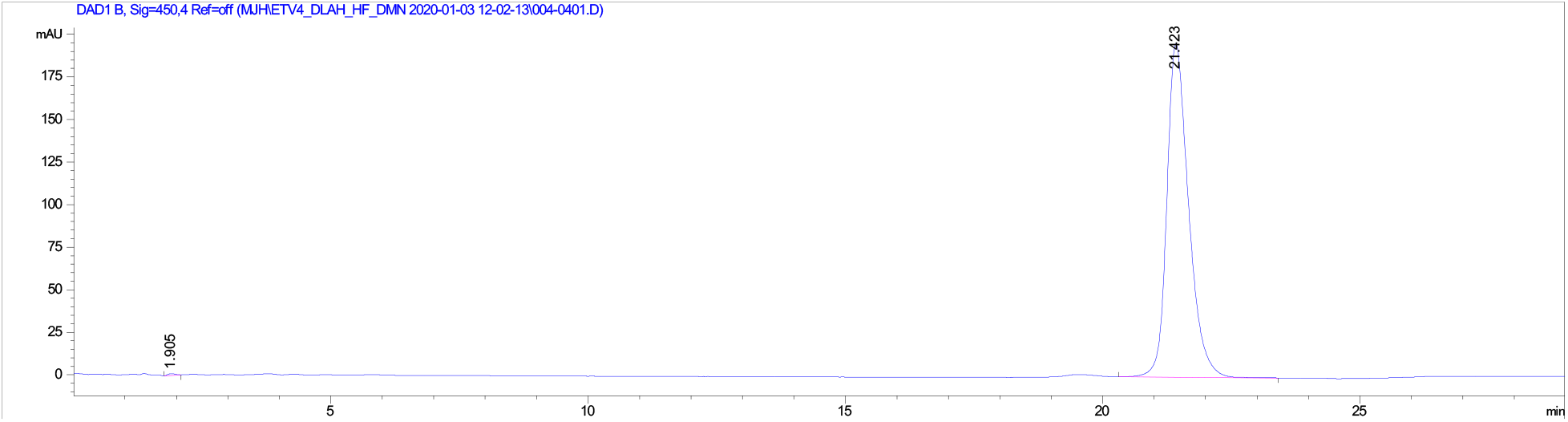

Analytical HPLC trace of **4-DMN-ETV4(45-76)^LPPL/HL^**, monitored at 450 nm. Analytical sample was run in a water (with 100 mM ammonium acetate)/ acetonitrile system. The sample was injected with an isocratic flow of 70% water (with 100 mM ammonium acetate) and 30% acetonitrile. After 2 mins, the solvent gradient was increased from 10-35% acetonitrile over 20 mins.

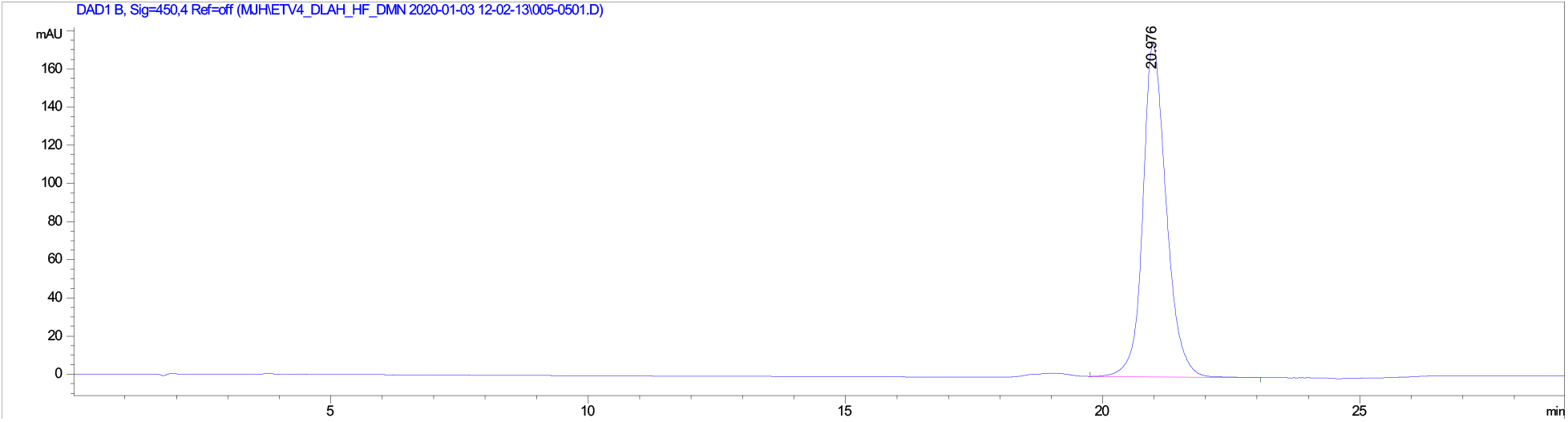

Analytical HPLC trace of **4-DMN-ETV4(45-76)^LPPL/QL^**, monitored at 450 nm. Analytical sample was run in a water (with 100 mM ammonium acetate)/ acetonitrile system. The sample was injected with an isocratic flow of 70% water (with 100 mM ammonium acetate) and 30% acetonitrile. After 2 mins, the solvent gradient was increased from 10-35% acetonitrile over 20 mins.

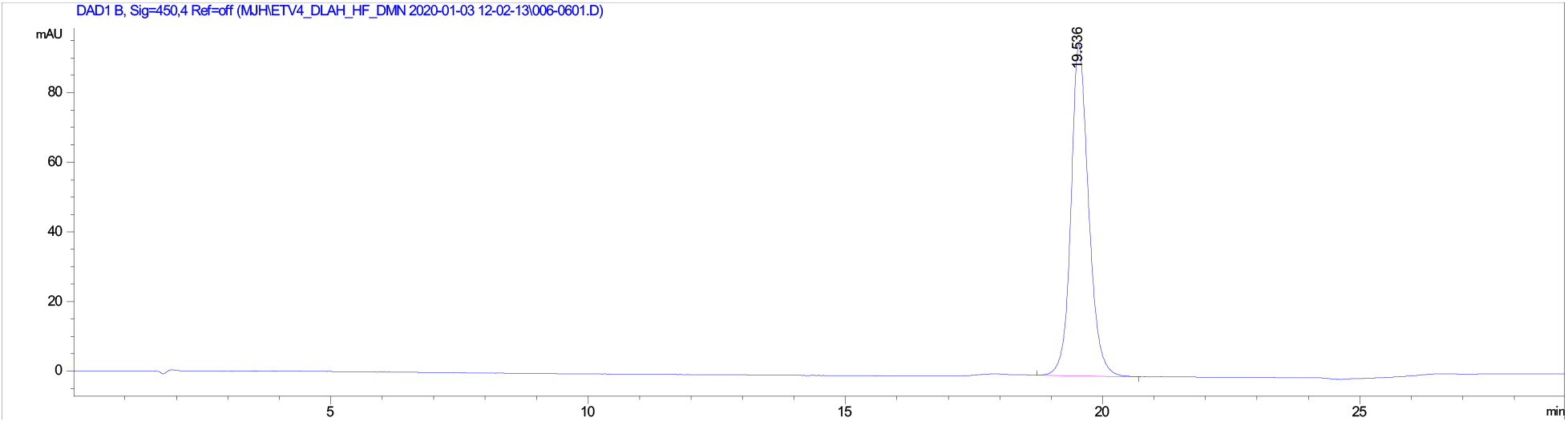

Analytical HPLC trace of **4-DMN-ETV4(45-76)^DLAH/HF^**, monitored at 450 nm. Analytical sample was run in a water (with 100 mM ammonium acetate)/ acetonitrile system. The sample was injected with an isocratic flow of 70% water (with 100 mM ammonium acetate) and 30% acetonitrile. After 2 mins, the solvent gradient was increased from 10-35% acetonitrile over 20 mins.

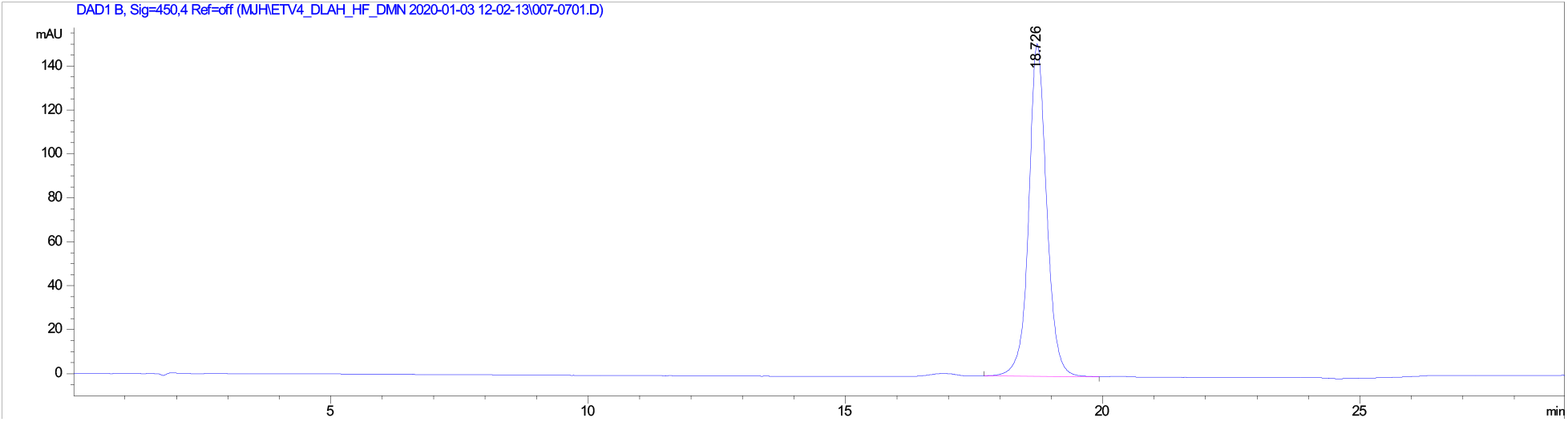

Analytical HPLC trace of **4-DMN-ETV4(45-76)^DLAH/QF^**, monitored at 450 nm. Analytical sample was run in a water (with 100 mM ammonium acetate)/ acetonitrile system. The sample was injected with an isocratic flow of 70% water (with 100 mM ammonium acetate) and 30% acetonitrile. After 2 mins, the solvent gradient was increased from 10-35% acetonitrile over 20 mins.

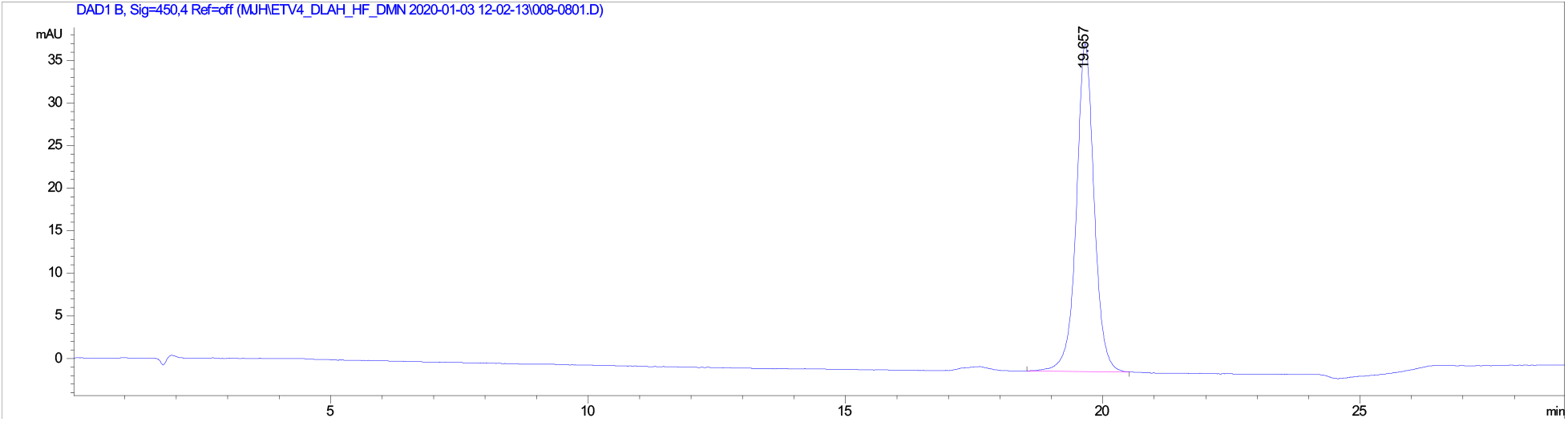

Analytical HPLC trace of **4-DMN-ETV4(45-76)^DLAH/HL^**, monitored at 450 nm. Analytical sample was run in a water (with 100 mM ammonium acetate)/ acetonitrile system. The sample was injected with an isocratic flow of 70% water (with 100 mM ammonium acetate) and 30% acetonitrile. After 2 mins, the solvent gradient was increased from 10-35% acetonitrile over 20 mins.

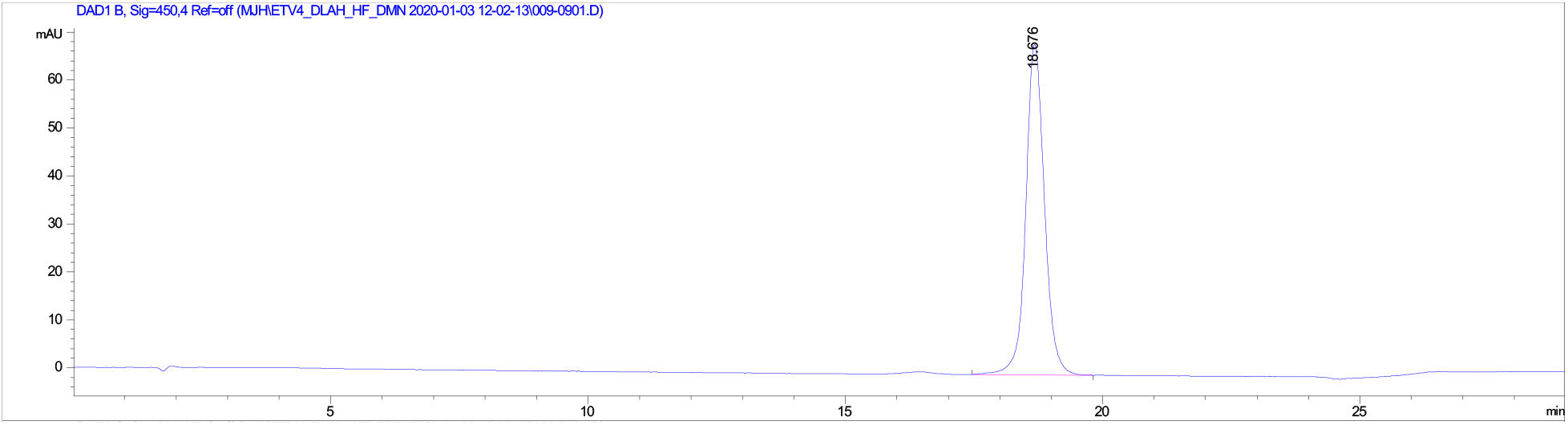

Analytical HPLC trace of **4-DMN-ETV1(42-69)**, monitored at 450 nm. Analytical sample was run in a water (with 100 mM ammonium acetate)/ acetonitrile system. The sample was injected with an isocratic flow of 70% water (with 100 mM ammonium acetate) and 30% acetonitrile. After 2 mins, the solvent gradient was increased from 10-35% acetonitrile over 20 mins.

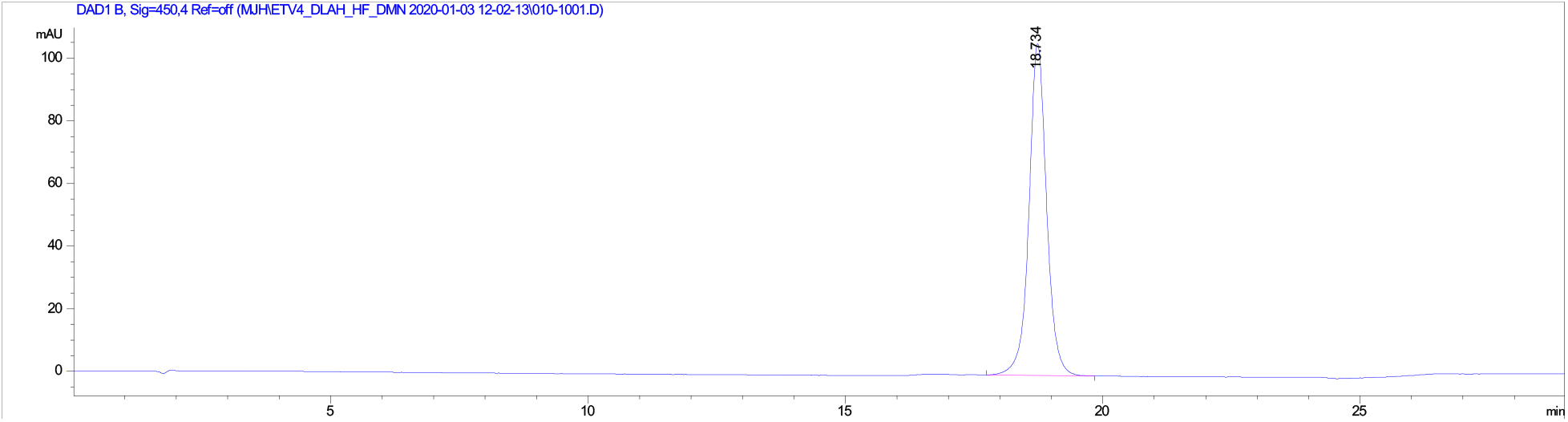

Analytical HPLC trace of **4-DMN-ETV4(49-76)**, monitored at 450 nm. Analytical sample was run in a water (with 100 mM ammonium acetate)/ acetonitrile system. The sample was injected with an isocratic flow of 70% water (with 100 mM ammonium acetate) and 30% acetonitrile. After 2 mins, the solvent gradient was increased from 10-35% acetonitrile over 20 mins.

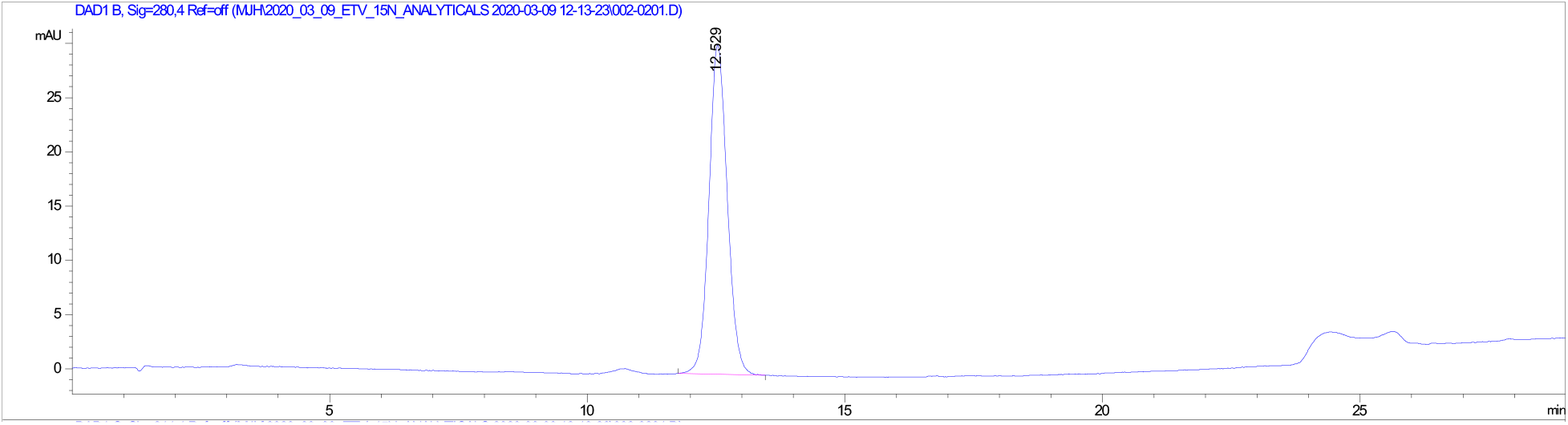

Analytical HPLC trace of **ETV1(38-69) ^15^N-Leu39**, monitored at 280 nm. Analytical sample was run in a water (with 100 mM ammonium acetate)/ acetonitrile system. The sample was injected with an isocratic flow of 70% water (with 100 mM ammonium acetate) and 30% acetonitrile. After 2 mins, the solvent gradient was increased from 10-35% acetonitrile over 20 mins.

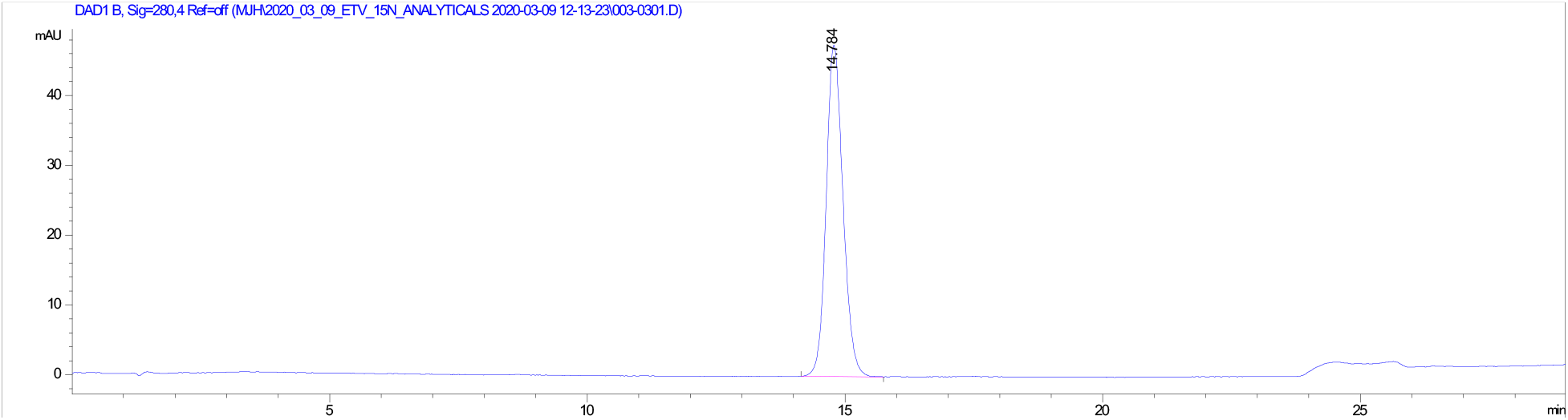

Analytical HPLC trace of **ETV4(45-76) ^15^N-Leu48**, monitored at 280 nm. Analytical sample was run in a water (with 100 mM ammonium acetate)/ acetonitrile system. The sample was injected with an isocratic flow of 70% water (with 100 mM ammonium acetate) and 30% acetonitrile. After 2 mins, the solvent gradient was increased from 10-35% acetonitrile over 20 mins.

